# Decoding Rice Seed Storage Proteins: From Gene Identification to Structural Prediction

**DOI:** 10.1101/2025.10.15.682491

**Authors:** Antima Yadav, Priya Jaiswal, Iny Elizebeth Mathew, Akanksha Panwar, Pinky Agarwal

## Abstract

Rice seed storage proteins (SSPs) serve as the primary source of dietary protein, energy, and nutrition; however, our understanding of these proteins remains limited, hindering efforts aimed at enhancing grain nutritional quality. To address this, we conducted a comprehensive genome-wide analysis of rice SSPs, encompassing phylogenetic relationships, domain and motif characterization, promoter *cis*-element mapping, expression profiling, and 3D structural modeling. Through homology- and domain-based searches, we identified 65 SSP-encoding genes, including 19 previously uncharacterized members. Phylogenetic analysis and domain comparisons revealed evolutionary proximity between albumins and prolamins; and between globulins and glutelins. Albumins, glutelins and prolamins clustered tandemly implying that duplication has driven SSP expansion. Transcriptomic evidence indicated that albumins, globulins, and glutelins were transcriptionally active from S2 seed development stage, whereas prolamins initiated expression from S3 stage. Promoter analysis revealed several seed-specific *cis*-regulatory elements, with CAATBOX1, EBOXBNNAPA, and DOFCOREZM being the most prevalent. Structural modeling showed that albumins and prolamins predominantly adopted α-helical conformations, whereas globulins and glutelins were enriched in β-strands and coils. This integrated analysis offers valuable insights into the classification, evolution, regulation, and structure of rice SSPs, providing a foundational resource for advancing grain nutritional improvement strategies.

## INTRODUCTION

Cereals are the most important food crops with an annual global production of 2859.2 million tons, considerably higher than the legume production worldwide (Fao, 2023). Despite less protein content in cereals compared to legumes, they provide more protein in the human diet as they are staple food crops, and grow on large cultivated areas. Seed storage proteins (SSPs) of cereals account for ∼10-12% of total seed dry weight and greatly impact seeds’ nutritional quality (Shewry and Halford, 2002; Shewry, 2023). Amongst cereals, rice forms a principal food and hence, is a crucial protein source for the majority of the world’s populations, especially in Asia and Africa (Bin Rahman and Zhang, 2023). As the nutritional value of cereal grains is affected by storage proteins, understanding the regulatory mechanisms involved in SSP biosynthesis and accumulation can help in increasing the protein content and nutritional quality of rice as well as other cereal grains by modulating the expression levels and proportion of different SSPs.

SSP synthesis is controlled spatially and temporally with high levels of regulation occurring at the transcriptional level in developing seeds (Yang et al., 2023). Rice SSPs are synthesized in the endosperm and a large amount is accumulated in storage organelles during later stages of seed development (Mahto et al., 2017). It is the second most important nutrient in rice after starch that plays a major role in determining grain quality (Gacek et al., 2018; Li et al., 2022). SSPs are broadly classified into four fractions, namely, albumins (ALBs), globulins (GLBs), glutelins (GLUs), and prolamins (PROs) (Shewry and Halford, 2002; Shewry, 2023). Albumins are non-glycosylated single-chain proteins of 10-15 kDa, usually present in dicots that are later processed into larger and smaller subunits attached by an interchain disulfide bond (Shewry and Halford, 2002; Shewry, 2023). Globulins are present in both monocots and dicots and occur either as 11S or 7S subunits. Globulins are present as pro-globulins, which are later processed by proteolysis and glycosylation (Sano et al., 2014; Gil-González et al., 2024). Glutelins are the most abundant SSPs that account for ∼ 60-80% of total protein, while prolamins constitute ∼ 20-30% of total seed protein in rice (Kawakatsu et al., 2008; Chen et al., 2018). Previous studies have revealed the role of SSPs in seed protection from oxidative stress, which is required for proper seed germination and seedling formation (Nguyen et al., 2015). They also play an important role in auxin homeostasis, and seed longevity and provide nutrition to the germinating seedling (Nguyen et al., 2015; Kumar et al., 2017; He et al., 2021). SSP content and composition determine the end-use nutritional value of the seed (Chen et al., 2018). Hence, to produce nutritionally superior cereal grains there is a need to explore new SSPs, and their regulatory pathways and further target them for molecular breeding and genetic engineering.

SSPs are highly expressed during the seed developmental process and regulated at different levels. The first level of regulation involves the biosynthesis of SSP which occurs majorly at the transcriptional level by specific binding of different transcription factors (TFs) to the *cis*-elements of *SSP* promoters (Shen et al., 2021). The second level of regulation involves synergistic or additive interaction among different transcription factors. The arrangement of *cis*-elements on the *SSP* promoter and the presence of interacting TFs are crucial for efficient SSP gene regulation (Yang et al., 2023). Some *cis*-regulatory motifs, for instance, GLM-GATA boxes are known to regulate *GLU-B1* expression and three *cis*-regulatory modules CCRM1, CCRM2, and CCRM3 are essential for *Glu-1* expression during wheat seed development (Li et al., 2019; Li et al., 2021). Several TFs as endosperm-specific bZIP factors OsSMF1/OsbZIP58/RISBZ1(Kawakatsu et al., 2009; Kim et al., 2017), REB/RISBZ2 (Nakase et al., 1997), and RITA-1/RISBZ3 (Izawa et al., 1994) have been identified as potential regulators of SSP gene transcription in rice. DOF factor RPBF (Kawakatsu et al., 2009), MYB5 (Suzuki et al., 1998), GZF1 (Chen et al., 2014; McDonald et al., 2024), and NAC20/26 (Wu et al., 2023) also play redundant role in expression regulation of rice *SSPs*. Reports on rice glutelin genes, *GluB-1* and *GluD-1*, have shown that combinatorial interactions of some *cis*-regulatory elements like GCN4, Prolamin-box, AACA, and ACGT motifs are essential for proper expression of glutelin genes (Kawakatsu et al., 2008; Qu et al., 2023). Another report shows the requirement of 5’ distal and proximal *cis*-elements for the proper expression of *Gt1* (Glutelin1) (Zheng et al., 1993; Queiroz et al., 2024). The bifactorial endosperm box consisting of the conserved prolamin box and the GCN4 motif has been identified as the necessary region for significant expression of *SSPs*. in many monocots (Marzábal et al., 1998; Cao et al., 2024). SSP synthesis is also regulated at post-transcriptional and post-translational levels, forming the third level of regulation in SSP biosynthesis. SSP accumulation is also affected by the presence of other biomolecules, metabolites, and transcript pools (Zhang et al., 2019). Previous reports have shown that reducing SSP synthesis or accumulation adversely affects nutrient quality and storage organelle formation in seeds (Kawakatsu et al., 2010). It has been shown that a consistent compensatory mechanism also exists that maintains the optimal level of different SSP fractions in rice. A reduction in an individual SSP fraction is accompanied by a subsequent increase in the level of all the other SSPs at both mRNA and protein levels (Kawakatsu et al., 2010). Several reports have suggested the involvement of other regulatory proteins in *SSP* gene expression. One such example is OsFIE1, which forms a complex with the other polycomb group/PcG members in the inner seed coat and suppresses the paternal *OsFIE1* in the endosperm (Cheng et al., 2020). This further regulates target TFs including MADS, NAC, MYB, and zinc finger family members by H3K27me3 modification and also modulates *SSP* gene expression (Huang et al., 2016; Badoni et al., 2023). Hence, these studies suggest the critical role of different TFs and their binding to specific *cis*-regulatory elements in the regulation and accumulation of SSPs in rice seed endosperm.

The information available thus far on rice SSPs in totality is scarce about their abundance in the rice genome, evolution and diversification, regulation of expression, accumulation pattern during seed development, and protein structure. To further this cause, we have performed extensive *in silico* analyses to identify and characterize total SSPs encoded in the rice genome, elaborating their genomic organizations, evolutionary relationships, functional domains, motif compositions, and *cis-*regulatory elements. In addition, we have deciphered the secondary and tertiary protein structures of all four classes of rice SSPs to underpin their structure-function relationships. We have also used microarray and qPCR data to map their expression levels and patterns during seed development. Our study is the first comprehensive portrayal of all regulatory aspects of rice SSPs that will facilitate future research on rice grain quality enhancement.

## MATERIALS AND METHODS

### Genome-wide *in silico* identification of SSP encoding genes in rice

For the genome-wide identification of *SSP* genes in rice (*Oryza sativa*), previously known SSPs belonging to more than ten plant species were searched in the existing literature. The SSP sequences of rice and other plant species were retrieved from MSU-RGAP 7 (MSU Rice Genome Annotation Project) (Ouyang et al., 2007) and NCBI (National Centre for Biotechnology Information) (Wheeler et al., 2007), respectively. These sequences were aligned using the Clustal X 2.0.11 program (Thompson et al., 1997), and were manually checked for removing the truncated and repeated sequences. The final sequences, thus obtained, were used to create a Hidden Markov Model (HMM) profile using the HMMER 3.0 software (http://hmmer.org/). This profile was used to search homologous protein sequences at MSU-RGAP 7. Identified rice sequences with cut-off E value ≤ 1e^-2^ were taken for further analysis. These sequences were searched for the presence of different seed storage protein-identifying domains in different databases like INTERPRO (Hunter et al., 2009), NCBI CDD (Conserved Domains Database) (Marchler-Bauer et al., 2015), ExPASY PROSITE (Gasteiger et al., 2003) and SMART (Simple Modular Architecture Research Tool) (Schultz et al., 1998) and classified into albumins, globulins, glutelins and prolamins based on defining domains. Sequences resulting from the HMMER search not having any SSP domain were discarded. The initial sets of protein sequences were further confirmed by motif and clustering analysis by MEME suite 4.12.0 (Multiple Em for Motif Elicitation) (Bailey et al., 2015) for verifying the presence of specific minimum motifs in them. Proteins that showed the conserved motif pattern were considered as SSPs and analyzed further.

### Chromosomal localization of rice *SSPs*

As per the position coordinates given in the RGAP database, rice *SSPs* were marked onto the twelve rice chromosomes. Rice *SSPs* were named with the letter word ALB, GLB, GLU, and PRO for albumins, globulins, glutelins, and prolamins respectively, followed by a number in the ascending order, corresponding to their positions on the chromosomes. Genes were depicted on the chromosome depending on the direction of transcription. Further, *SSPs* were analyzed for tandem and segmental duplication events as mentioned previously (Agarwal et al., 2007). SSPs belonging to the same family present on the same chromosome separated by 5 or a lesser number of genes were considered as tandemly duplicated. The *SSP* loci were compared with the rice segmental duplicated data from RGAP generated using 500 kb as the maximum distance allowed between two collinear gene pairs.

### Phylogenetic analysis of SSPs from rice and Poaceae

For analyzing the evolutionary relatedness amongst SSPs, two phylogenetic trees were constructed. A phylogenetic tree consisting of 65 SSPs from rice was constructed to analyze the evolutionary relatedness amongst the various classes of rice SSPs. However, for analyzing the evolution of the rice SSPs within the family *Poaceae*, a tree was constructed using 126 amino acid sequences including the previously reported protein sequences from other species as well (Table S1). Phylogenetic trees were generated and visualized through MEGA6 software, using the Maximum Likelihood method based on the Jones-Taylor-Thomson (JTT) matrix-based model (Jones et al., 1992). A discrete gamma distribution was used to model the evolutionary rate difference amongst sites. Both the trees were drawn by performing the bootstrap test of phylogeny with 1000 iterations.

### *In silico* identification and visualization of seed-specific *cis*-regulatory elements in rice SSP encoding gene promoters

For promoter analysis in 65 rice *SSPs*, upstream 2 kb promoter sequences from the translation start site were retrieved from MSU-RGAP 7 and NCBI (National Centre for Biotechnology Information). The promoter sequences were analyzed for *cis*-regulatory elements using the PLACE database (Higo et al., 1998). The identified *cis*-elements were arranged and compared based on their frequencies, order of occurrences, and their presence on positive and negative strands across all the *SSPs*. To understand the distribution pattern of *cis*-elements, the 2 kb promoter region of each *SSP* was divided into 100 bp fragments, creating 20 bins, and all *cis*-elements were marked for their presence in every bin. Seed-specific *cis-*elements were arranged on the 2 kb upstream promoter region for all the *SSPs* using R-studio.

### Plant material

Rice plants (*indica* var. IR64) were grown under field conditions at NIPGR, New Delhi. Panicles were tagged from the day after pollination (DAP) till 29 days, for the collection of seeds. Seeds were harvested at different seed developmental stages, categorized as S1 (0 to 2 DAP), S2 (3 to 4 DAP), S3 (5 to 10 DAP), S4 (11 to 20 DAP) and S5 (21 to 29 DAP) (Agarwal et al., 2011). Seeds were frozen immediately in liquid nitrogen and kept at -80°C for further analysis. Flag leaves used as controls were collected and frozen in liquid nitrogen and stored at -80 °C.

### Expression analysis by microarray and quantitative RT-PCR

Unique Affymetrix probe set IDs were retrieved from ROAD (www.ricearray.org/expression/meta_analysis.shtml) for microarray analysis of the identified *SSP* genes. The expression was examined in above mentioned five seed development stages and young leaves, from our previously published microarray data (Agarwal et al., 2007; Sharma et al., 2012; Agarwal et al., 2016). qPCR was performed for genes that either did not have or had ambiguous probe sets. For this, total RNA was extracted from 100 mg seeds in three biological replicates of each seed developmental stage (S1 to S5) according to previously reported seed-specific protocol (Singh et al., 2003) with few modifications, followed by DNase treatment to purify the RNA samples by using RNeasy^®^ MinElute Cleanup Kit (Qiagen) according to manufacturer’s protocol. RNA was isolated from flag leaf tissue using TRI Reagent^®^ Solution from Invitrogen. The quality and quantity of RNA samples were checked by NANODROP 2000 Spectrophotometer (Thermo Scientific) and integrity was analyzed by running 1 μl RNA sample in Bioanalyzer (Agilent 2100). RNA samples with RIN (RNA integrity number) value above 8.0 were used for first strand cDNA synthesis using RNA High Capacity cDNA Reverse Transcription Kit (Applied Biosystems). cDNA was made in biological triplicates for each stage and flag leaf. qPCR assay was carried out with KAPA SYBR FAST qPCR Kit Master Mix (2X) Universal (KAPA BIOSYSTEMS) with 1µl cDNA template (1:10 Dilution) as per manufacturer’s protocol using Applied Biosystems 7500 Fast Real-Time Systems and 7500 software v2.0.1. The expression level of each gene was calculated by the ΔΔCt method (Rao et al., 2013), where *Actin* (*ACTI*) was used as an internal control. To assess microarray and qPCR data together, log2fold change was calculated for each gene and the seed development stages (S1 to S5) with respect to the flag leaf. The data was visualized as a heat map in MEV.

### *In silico* secondary and tertiary structure prediction of rice SSPs

Secondary structures of all SSPs were predicted by PsiPred (McGuffin et al., 2000) using default parameters (http://bioinf.cs.ucl.ac.uk/psipred/). The 3D structures of albumins (except ALB8, 9, 10 and 15), globulins, glutelins (except GLU6, 8, 10, 17, and 20), and prolamins (except PRO5, and PRO20) were extracted from the AlphaFold Structure Database (Varadi et al., 2022) (https://alphafold.com/). The 3D structures of the proteins were downloaded in PDB format and visualized using PyMol (https://pymol.org/2/). For 10 proteins with unavailable 3D structures, the abinitio method using Alphafold2 Colabfold platform was used to predict their structures (https://colab.research.google.com/github/sokrypton/ColabFold/blob/main/AlphaFold2.ipynb). This structure prediction method generated five models per protein with varying pLDDT scores. The best structure for each protein was selected based on the highest average pLDDT score, indicating the model’s reliability.

## RESULTS

### *In silico* identification of putative SSP encoding genes in rice

Genome-wide HMMER search identified a total of 65 SSP encoding genes in the rice genome, including 19 newly discovered SSPs. This study resulted in the identification of 16 albumin proteins in rice, which have been named as ALB1-16 based on their chromosomal positions (Figure 1, Table S2). Out of these, only six (ALB1, 2, 3, 8, 9, and 16) have been previously reported (Wu et al., 1998; Xu and Messing, 2009; Zhou et al., 2017) and the rest have been newly identified in this analysis. All albumins contained the identifying functional 2S albumin domain (Figure 2). ALB1, 2, 3, and 16 are reported as albumins in MSU-RGAP 7. In this analysis, LOC_Os04g13540 designated as a putative albumin, was excluded because of the absence of any relevant domain. ALB8 and ALB9 are reported as RAL3 and RAL4 seed-allergenic proteins, respectively in MSU-RGAP 7. Most of the identified albumins were present on chromosome 7, while the remaining albumins were distributed on chromosomes 3, 5, and 11. Screening of the albumin proteins for tandem duplication identified four tandemly duplicated clusters localized on chromosomes 3 and 7 (Figure 1, Table S3). Gene evolution following tandem duplication has been reported for rice *SSPs* (Xu and Messing, 2009).

**Figure 1.**
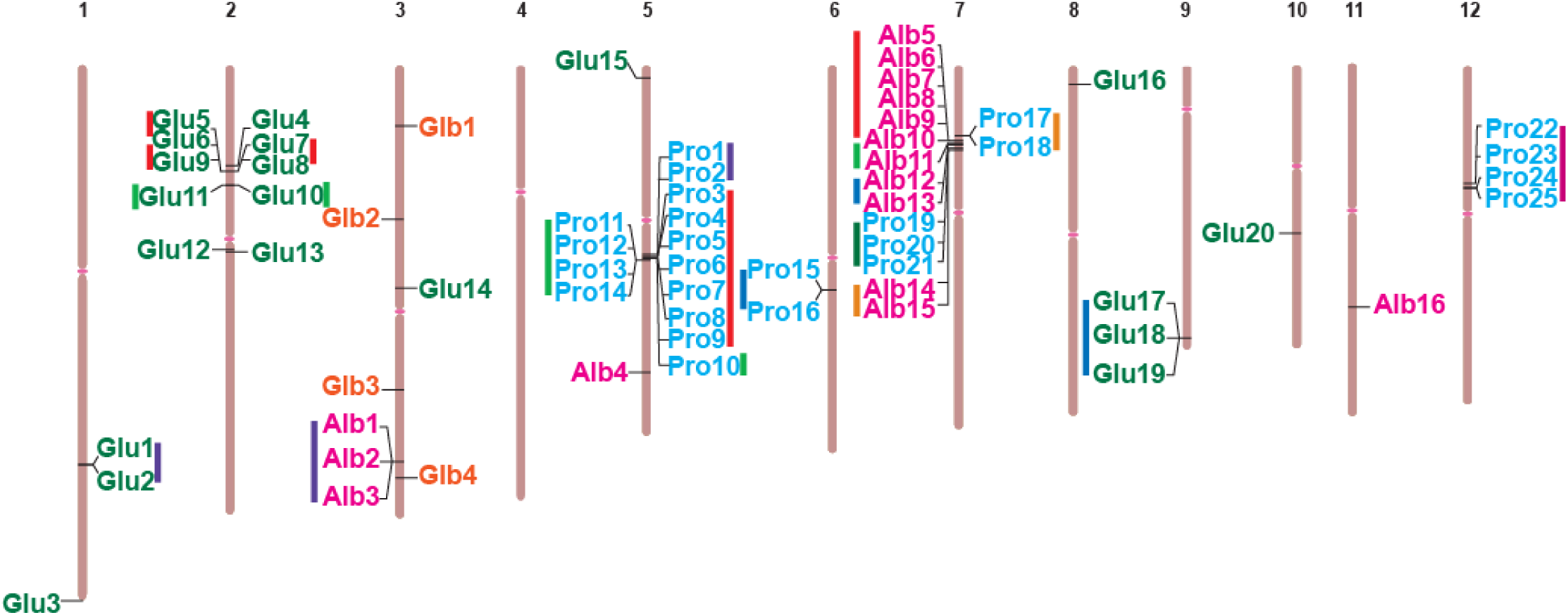
Chromosomal localization of seed storage protein (*SSPs*) genes in rice. Distribution of the 65 welve rice chromosomes. The chromosome numbers are indicated above each chromosome. The ach chromosome as well as the position of the genes has been made according to the coordinates SU-RGAP 7. The genes are marked according to their occurrence on the chromosome with ORFs osite orientation marked on either side. The genes falling within the tandemly duplicated groups pecified by different colored lines.

**Figure 2.**
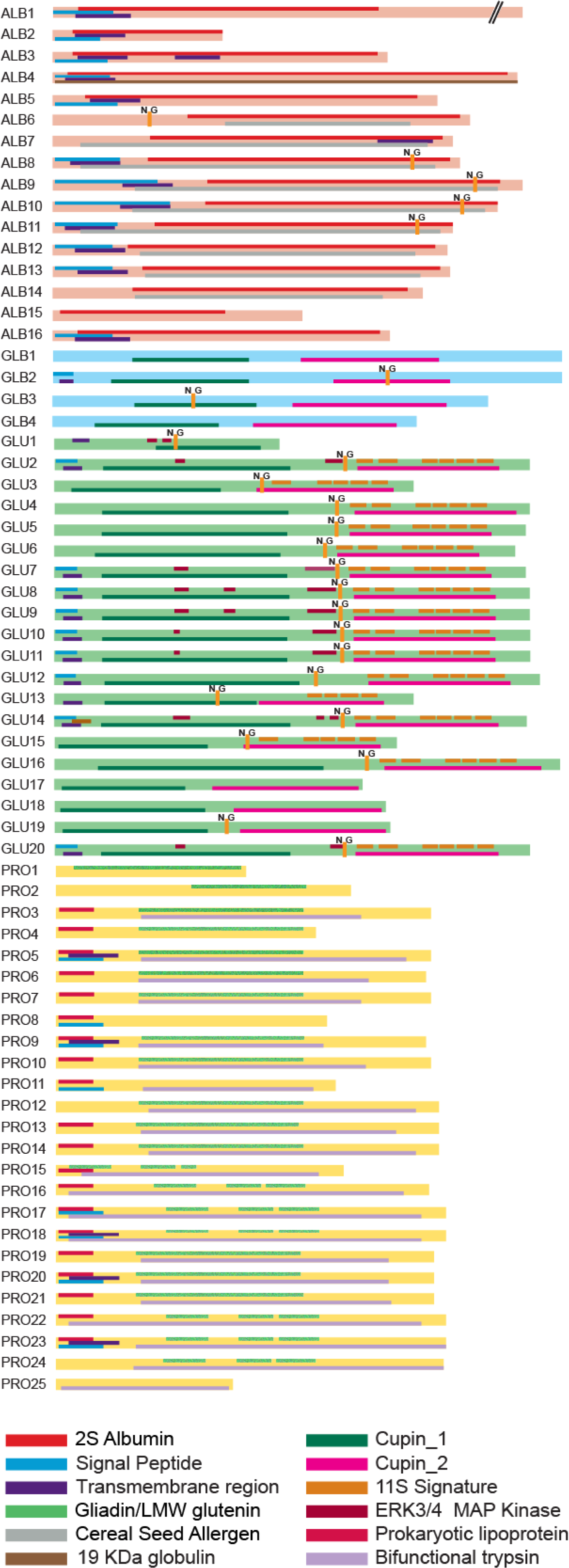
Domain analysis in various classes of SSPs. The four different classes of SSPs can be distinguished by the presence of certain domains, which are marked with specific color as mentioned at the lower end of the figure. Proteolytic cleavage site has been shown by an insertion mark with N_G written in the flanking position. The diagram is drawn to scale.

Only four rice loci were considered as coding for globulins as these showed the presence of specific cupin domains and conserved pattern in motif arrangement. All these newly identified globulins were named as GLB1-4. Three proteins reported as globulins in MSU-RGAP 7 and LOC_Os05g41970 reported as globulin in a previous publication (Wu et al., 1998) lacked the specific domain, motif, and tree clustering pattern and were omitted from further analysis. LOC_Os05g41970 has been identified as coding for albumin in our study. All the globulin proteins were localized onto the third chromosome (Figure 1), though they were not tandemly duplicated. For the identification of glutelin proteins, the analyses resulted in the identification of 20 unique loci, which contained the signature domains and were named as GLU1-20. Glutelins included five new proteins (GLU3, 15, 17, 18, and 19) along with fifteen reported glutelin proteins in MSU-RGAP 7 (Table S2). Thirteen glutelin proteins clustered into four tandemly duplicated groups (Figure 1, Table S3). Out of these thirteen, eight glutelins belonging to tandem duplicated clusters were localized on the second chromosome whereas two were localized on chromosome one and three on chromosome nine. Additionally, GLU3 (LOC_Os01g74480) and GLU15 (LOC_Os05g02520) resulted from a segmental duplication event.

Analysis for prolamins resulted in 25 sequences containing the prolamin-specific domain. Since, there were discrepancies in the naming of prolamins in MSU-RGAP 7, all 25 prolamins were renamed as PRO1-25 based on their chromosome positions. Prolamin proteins satisfying the tandem duplication criteria were classified into 8 groups. Fourteen prolamins, belonging to four tandem duplicated clusters, were present on chromosome 5, while the remaining 11 prolamins were localized on chromosomes 6, 7, and 12 as four tandem duplicated clusters (Figure 1, Tables S2, S3).

### Distribution of motifs and domains in rice SSPs

The structure and function of different proteins are attributed by the presence of characteristic functional domains in them. Rice SSPs showed the presence of distinct domains categorizing them into various subfamilies (Figure 2). Albumins had the signature 2S albumin domain encompassing the majority of the protein length. Additionally, many of these proteins had transmembrane and a signal peptide region predicted at their N-terminal end. Some of these proteins (ALB6, 7, 8, 9, 10, 11, 12, 13, and 14) contained the cereal seed allergen domain. Out of these, ALB6, 7, 8, 9 10, and 11 have been reported as seed allergenic proteins in MSU RGAP. MEME analysis (Figure S1) showed that ALB1, 2, 3, and 16 had a motif (motif 9) at the extreme N-terminal end that was shared with 20 other prolamins. All the albumins except ALB2 had motif 18, which was present within the 2S albumin domain and contained the CCxQL motif (Figure S2). This motif was also present in PRO15 and 16, between the first and second gliadin stretch. Motif 16, present in ALB12 and 13 was shared with 10 glutelin proteins.

As in the case of other reported globulin proteins, rice globulins also had two cupin domains. Globulins belong to the cupin superfamily of proteins and are characterized by their conserved β-barrel fold formed from the two cupin domains (Kesari et al., 2017). Out of the four globulins, GLB2 and GLB3 had a conserved APE cleavage site (Figure 2). Globulin precursor proteins have also been shown to undergo different post-translational modifications including proteolytic cleavage and glycosylation. All the rice globulin proteins showed the presence of motif 24, which was also shared by three glutelins (GLU17, 18, and 19), present within the cupin 2 domain (Figure S1).

All the glutelin proteins except GLU1 were bicupin proteins like the other reported globulin proteins (Figure 2). The rice glutelins are said to be essentially globulins, showing a high homology with the latter (Zhao et al., 1983; Kawakatsu et al., 2008). Hence, many of these proteins excluding GLU1, 17, 18 and 19 contained the signature 11S globulin domain found to overlap with the second cupin domain. Most of these proteins also contained an asparaginyl cleavage site, absent only in Glu17 and 18 (Figure 2). Most of these proteins except GLU17, 18 and 19 contained motif 3, overlapping with the third 11S globulin stretch (Figure S1). All the glutelin proteins, other than GLU1 contained motifs 6 and 4, present in the first and second cupin domain, respectively. Motifs 5 and 12, within the first cupin domain, and motif 11 outside the cupin domain, were present in many glutelin proteins. Some motifs like motif 22, 23 and 25 were present only in GLU3, 15, 17, 18 and 19.

MEME analysis showed high motif conservation and sequence homology amongst prolamins as well (Figure S1, S2). Motif 2 containing the CCxQL sequence and motif 8 were present in most of these proteins. MEME and domain analysis showed similarity between albumin and prolamin proteins as well as between globulin and glutelin proteins on rice (Figure 2, Figure S1). Apart from the Gliadin/LMW glutenin signature domains, the prolamin class of proteins contained several other domains like the bifunctional trypsin/alpha amylase inhibitor helical domain, signal peptide, transmembrane region and prokaryotic lipoprotein domain as shown in Figure 2.

### Phylogenetic analysis of rice and grass family SSPs

Phylogenetic analysis of rice SSPs has been carried out using the Maximum Likelihood method and also done by taking into account all the SSPs reported in different Poaceae species. From the unrooted phylogenetic analysis of 65 SSPs, it has been found that 65 rice SSPs were divided into two main clades, one comprised of albumins and prolamins and the second of glutelins and globulins (Figure 3). Mostly, tandemly and segmentally duplicated proteins arose from the same branch. Albumin-prolamin clade branched out further in six sub-clades. The first subclade was constituted by the proteins previously reported as allergens ALB7-11 and ALB12-15 in MSU-RGAP7. ALB1, 2, 3, and 16, which have been reported previously as 10 kDa prolamins in rice (Xu and Messing, 2009), formed the second subclade. ALB4 branched out from the second albumin subclade and ALB5 appeared as an outgroup to the first and second albumin subclades (Figure 3). The phylogenetic tree clearly showed the clustering of all the prolamins into a single group within the albumin-prolamin clade. The prolamin subclade was further divisible into three sub-groups. Out of these, the first two were constituted by PRO22, 23, 24, and 25 located on the 12th and PRO17 and 18 on the 7th chromosomes, respectively. These prolamins have been described as the 16 kDa prolamins in rice (Kawakatsu et al., 2008; Xu and Messing, 2009). All the remaining prolamins clustered together and formed the third sub-group. It was constituted by seventeen b-type 13 kDa proteins (Figure 3). Thus, the strong conservation in the protein sequences as observed in domain and motif analysis was further confirmed by the close phylogeny of these proteins (Figure 2, 3 and Figure S1).

**Figure 3.**
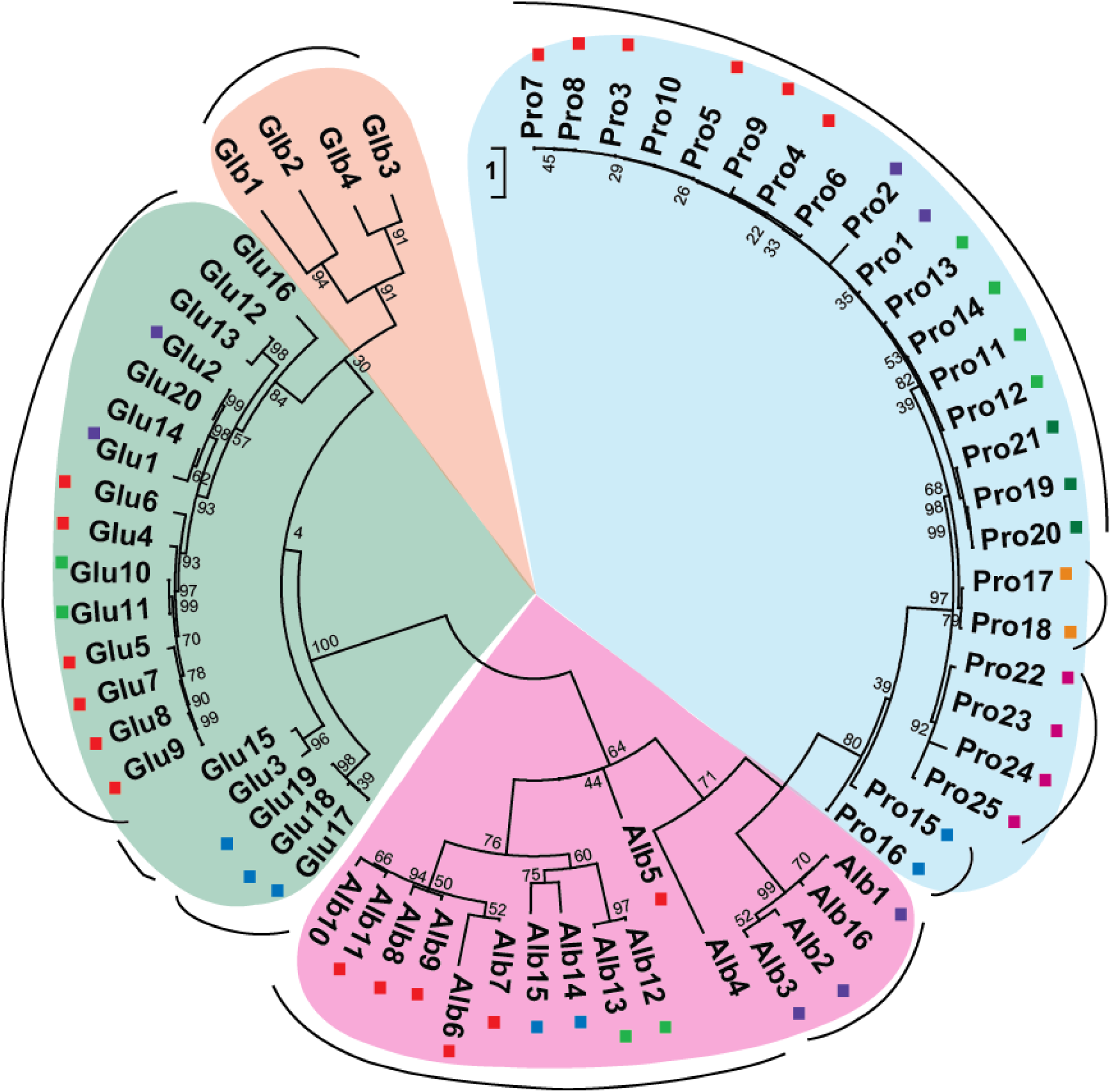
Phylogenetic analysis of rice SSPs. The unrooted tree of rice SSPs has been constructed by using maximum likelihood (ML) method based on the JTT matrix based model. The tree has been drawn to scale with branch length denoting the rate of amino acid substitution. Bootstrap values are shown at the branch points. SSPs belonging to the four major classes are boxed in different colors; with subclades represented by an arc. The tandemly duplicated genes within each class are highlighted using identically colored boxes.

The second major clade in the rice SSP tree was formed by the globulin and glutelin proteins which formed four distinct sub-groups. The first sub-group was formed by the tandem duplicated genes, GLU17, 18, and 19 present on chromosome 9 (Figure 1 and 3). The segmentally duplicated pair GLU3 and 15 formed the second sub-group. MEME patterning has also shown the sharing of different motifs (motifs 22, 23, and 25) amongst these proteins (Figure 3 and Figure S1). All the remaining glutelins grouped together to form the third subclade in the globulin-glutelin major clade. The four globulin proteins formed the fourth subclade (Figure 3). The branching of the globulin and the glutelin proteins from the same node agreed well with the domain and MEME analysis of these proteins. All these results suggested a common ancestral origin for these two types of SSPs.

A similar analysis of rice SSPs with the SSPs belonging to different subfamilies of Poaceae, further corroborated the clustering pattern amongst rice SSPs. Various proteins used for the phylogenetic analysis have been listed in Table S1. Briefly, two proteins belonging to each category within a tribe have been used for the tree construction. The maximum likelihood tree showed a clear distinction into the albumin-prolamin clade and globulin-glutelin clade (Figure 4). The albumin-prolamin clade showed distinction four sub-groups, named as group I-IV. Group I was constituted by four rice albumin proteins, reported previously as 10 kDa prolamin proteins. The clustering of four rice albumins (ALB1, 2, 3 and 16) together with α and δ-prolamins, reported in other plant species suggest a comparatively recent evolution of these proteins. All the rice prolamin proteins grouped with the β-prolamins were categorized as group II proteins. Other proteins as LMW prolamins, γ and ω prolamins, sharing high homology with the β-prolamins formed an entirely distinct cluster categorized as group III in the phylogenetic tree, with no rice protein grouped here. The fourth group within the albumin-prolamin clade was formed by the albumin proteins showing their grouping with the HMW prolamin and the 2S globulin proteins. Amongst prolamins, HMW proteins are the oldest, initially originated from the α-globulins (Xu and Messing, 2009). Thus, the unrooted phylogenetic tree comprising the Poaceae SSPs showed the origin of prolamins and albumins from a common branch point (Figure 3 and 4). This also suggested a common ancestor for these two families of proteins in rice.

**Figure 4.**
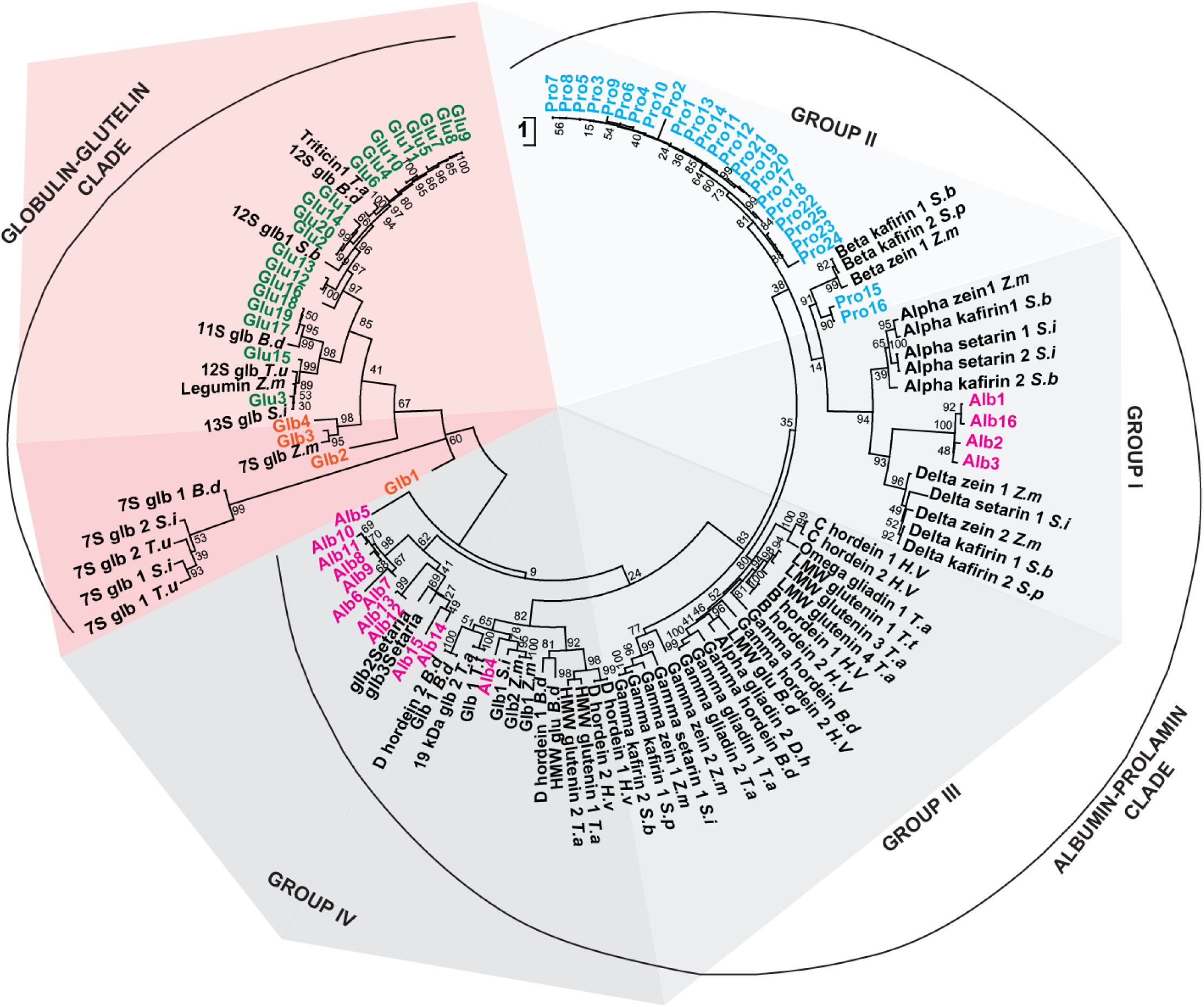
Evolutionary analysis of grass SSPs. The unrooted tree of grass SSPs has been constructed by taking into account the various classes of SSPs reported in different plant species within the grass family including 65 rice SSPs, using maximum likelihood (ML) method. Rice SSPs identified in the present study are uniquely color coded for each class. The tree has been drawn to scale with bootstrap test of phylogeny with 1000 iterations. Branch length is proportional to the rate of amino acid substitution as calculated by JTT model. Members of two major clades are shown by multiple shades of pink and grey color.

The second clade in the phylogenetic tree was constituted by the globulin and glutelin proteins. The globulin-glutelin clade formed an entirely separate clade in the tree and could be clearly divided into the 11S-12S clade and the 7S-9S globulin clade. All rice glutelins belonged to the 11S-12S group, whereas the four rice globulins belonged to the 7S-9S group (Figure 4). Evolutionary studies revealed the sequence homology of 11S legumin and 7S vicilin proteins. Both these proteins have a common single-domain ancestor (Barasa, 2022). Both 7S and 11S globulin proteins are known to have an equivalent N-terminal and C-terminal region which forms β-barrel shaped cupin domain, despite their relatively lesser sequence homology (Costa and Mafra, 2023; Hu et al., 2023).

### Identification and analysis of *cis*-elements in the promoter region of rice *SSPs*

To investigate the role of *cis*-elements in the regulation of *SSP*s, the 2 kb upstream promoter regions of 65 seed storage proteins (SSPs) were analyzed. A total of 67 seed-specific *cis*-elements were identified using the PLACE database (Figure 5; Table S4). These elements, ranging from 4 to 19 base pairs in length (with the majority between 6-7 bp), have been reported to play essential roles in seed development across various plant species. Many *cis*-elements contained ambiguous bases, leading to multiple sequence variations. As a result, some elements shared identical sequences; for example, 2SSEEDPROTBANAPA and CANBNNAPA were found to be sequence-identical (Table S5). The expansion of ambiguous bases resulted in a totality of 255 seed-specific elements present of the promoters of *SSP*s. The analysis of these *cis*-elements across the four major SSP categories revealed significant variations in their numbers and distribution (Table S6). Certain elements, such as CAATBOX, DOFCOREZM, EBOXBNNAPA, NTBBF1ARROLB, POLASIG1, PYRIMIDINEBOXOSRAMY1A, RYREPEATBNNAPA, SEF4MOTIFGM7S, TATABOX5, WBOXHVISO1 and WRKY71OS were among the most frequently occurring across all *SSP* promoters. Meanwhile, some *cis*-elements were unique to specific SSP groups. This included CEREGLUBOX3PSLEGA, PROXBBNNAPA, and CEREGLUBOX1PSLEGA which were exclusive to *ALB* promoters, while ACGTSEED3 and ARE1 were specific to *GLB* promoters. Similarly, AACAOSGLUB1, GLUTAACAOS, GLUTEBOX1OSGT2, GLUTEBOX1OSGT3, GLUTEBOX2OSGT2, GLUTEBOX2OSGT3, GLUTEBP1OS, GLUTEBP2OS, GLUTECOREOS and SPHCOREZMC1, and were exclusively present on *GLU* promoters while TATABOX1, CEREGLUBOX2PSLEGA and OPAQUE2ZMB32 were unique to *PRO* promoters.

**Figure 5.**
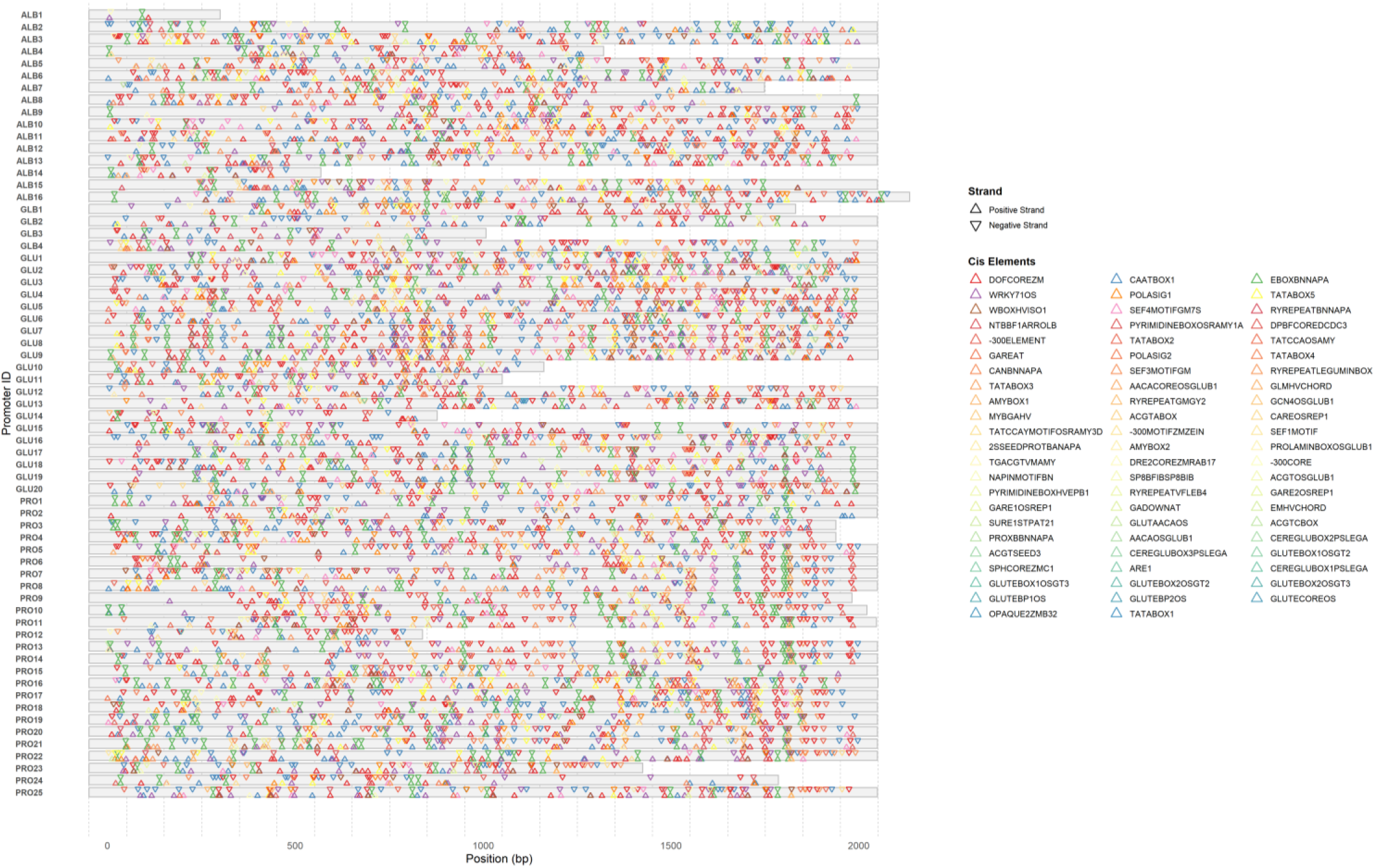
Analysis of seed-specific *cis*-elements in the promoter sequence of *SSPs*. The seed-specific *cis-*elements present on the 2 kb upstream promoter sequence of 65 *SSPs* are shown. Each element is denoted by a colored triangle as per the legend on the right.

Furthermore, some elements, such as EMHVCHORD and RYREPEATVFLEB4 were entirely absent in *ALB* promoters, while 2SSEEDPROTBANAPA, -300CORE, 300MOTIFZMZEIN, AMYBOX2, CEREGLUBOX1PSLEGA, EMHVCHORD GARE1OSREP1, GARE2OSREP1, GLMHVCHORD, GLN40SGLUB1 and SURE1STPAT21, were absent in *GLB* promoters. To further analyze the spatial distribution of these *cis*-elements, the promoter sequences of each SSP were divided into 20 bins of 100 bp each (Table S7). While most *cis*-elements were dispersed throughout the promoter regions, certain elements such as, 300MOTIFZMZEIN, AACAOSGLUB1, GCN4OSGLUB1, GLMHVCHORD, GLUTEBOX1OSGT2, GLUTEBOXOSGT3, GLUTEBOX2OSGT2, GLUTEBOX2OSGT3, GLUTEBP1OS, GLUTCOREOS and TATABOX1 were predominantly concentrated near the 3’ end of the promoter sequence. This suggests a potential regulatory role of these elements in fine-tuning SSP expression, likely influencing transcriptional activation at key developmental stages. Overall, this study highlights the diversity and distribution patterns of seed-specific *cis*-elements in *SSP* promoters. These insights pave the way for further functional validation of *cis*-elements in seed development and storage protein biosynthesis.

### Expression analysis of different *SSPs* in rice seed developmental stages

To analyze the expression of 65 *SSP* genes, we used our previously published microarray data (Agarwal et al., 2007; Sharma et al., 2012). Affymetrix probe sets were obtained for all genes from ROAD and out of 65 *SSP*s, unique probe sets were found for 36 of them including 15 *ALBUMINS*, 3 *GLOBULINS*, 14 *GLUTELINS*, and 4 *PROLAMINS*. Normalized signal values are presented in Table S8. qRT-PCR was performed for 25 remaining genes *i.e. ALB3*, *GLU7-11*, *PRO1-5*, *7*, *8*, *10*, *11-14*, *19-25* (primers in table S9). Due to high homology, the expression of *GLU8-9* and *10-11* was analyzed together. Unique qPCR primers were designed for 13 prolamins genes, while *PRO 5-7-10* and *PRO19-20-21* had common primers. qPCR could not be performed for GLB2, *GLU17*, *PRO6* and *PRO9*. The expression data revealed that most of the genes were expressed at high levels during seed development stages of rice (Figure 6). Out of 61 genes, analyzed for the expression analysis, 36 genes were more than 2 folds up-regulated in the S2-S5 stages of seed development, whereas *ALB3*, *GLU15*, *16*, *19* were up-regulated throughout seed development (Figure 6, Table S8 and 10). The genes *ALB9*, *ALB15*, *GLU3,* and *GLU18* did not show any expression. *PROs* showed a variable expression pattern, *PRO1*, *13* were highly up-regulated in S5 stage, *PRO4*, *5, 7, 10*, *8*, *11*, *12*, *14*, *19, 20, 21*, 22, 23, 24, 25 were up-regulated in S3-S5 stages while four *PROs* (*15*, *16*, *17*, *18*) expressed in S2-S5 stages of seed development. Most *ALB*, *GLU*, and *GLB* were up-regulated in the S2-S5 stages while *PROs* were up-regulated in the S3-S5 stages of seed development. Thus, this analysis indicates that *SSP* transcripts start accumulating early during seed development.

**Figure 6.**
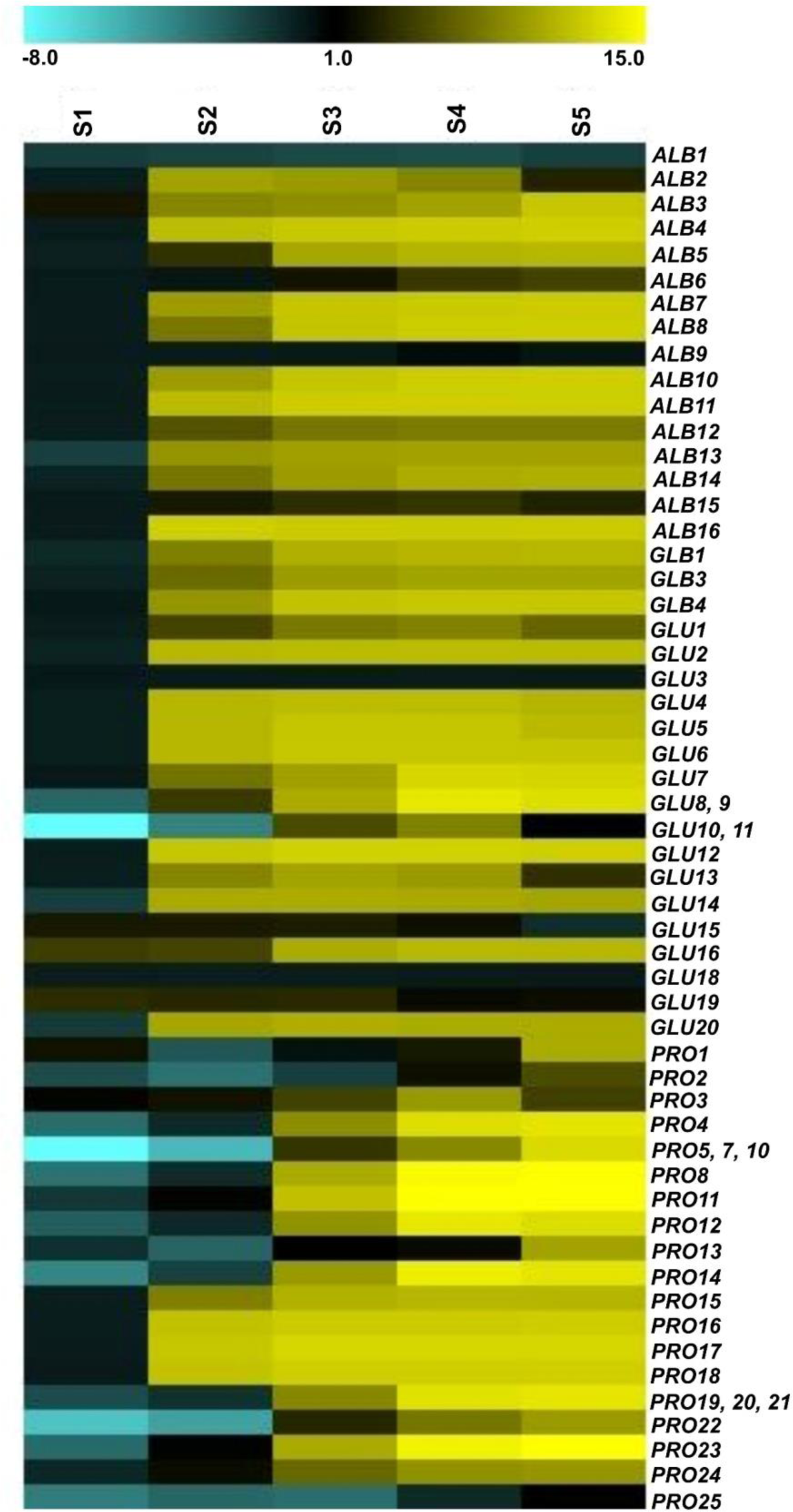
Expression profile of different *SSPs* in rice. Heat map representing the relative expression pattern of 65 *SSPs* in different rice seed developmental stages (S1-S5). Log_2_ fold change values of the expression signals have been used for generating the heat map in MeV Version 4.9.0. As per the color coding given in the legend, up and down regulated genes are indicated by yellow and cyan blue colors, respectively.

### Secondary and tertiary structural characterization of SSPs

The biological function of a protein is determined by its three-dimensional structure. To understand the functional divergence among the various SSPs, their higher-order structures have been predicted. Total amino acid length in albumins ranged from 68-118 residues, where ALB1 was exceptionally long with 340 amino acids. SVM-prot analysis predicted these proteins as belonging to lipid-binding or metal/ion-binding protein families (Table S11). The predicted secondary structure for rice albumins consisted mainly of helices and coils. Few albumins, like ALB4 and 9 had β-strand structure towards the N-terminal, mostly outside of the helical region (Figure S3a). Rice globulins are larger proteins ranging in size from 470-565 amino acid residues. Similar to albumins, all these four proteins belonged to lipid binding and metal-ion binding families (Table S11). Major secondary structural elements present in globulins were coils and β-strands (Figure S3b). Multiple small helical stretches were also distributed over the protein. The two cupin domains were organized into multiple β-strands connected by coils. Rice glutelins were also predicted to be lipid-binding and/or metal/ion-binding proteins (Table S11). They formed large proteins with 325-500 amino acids in length. Just like globulins, the secondary structure of glutelins consisted majorly of strands and coils, with few interspersed helices (Figure S3c). Cupin domains were organized as a β-stranded structures separated by coils. Prolamins were also predicted as metal/ion and lipid-binding family proteins (Table S11). They formed smaller proteins ranging between 76-156 amino acid residues in length. All prolamins were predicted to have a secondary structure predominated by 4-5 helical structures connected by coils (Figure S3d). PRO2 and 9 had a short stretch of a β-strands.

For 3D protein structure analysis of rice SSPs, PDB structures were extracted for most of albumins (AlB1-7, ALB11-16), all globulins (GLB1-4), most of glutelins (GLU1-5, GLU7, GLU9-16, GLU18-19), and most of prolamins (PRO1-4, PRO6-19, PRO21-25) from the AlphaFold Database (Figure 7, Table S13). Protein structures were assessed on the basis of per-residue confidence scores (average pLDDT values). Each protein with their UniProt ID, protein symbol, sequence length and confidence scores are summarised in Table S13. pLDDT scores above 90, between 70-90, between 50-70 and below 50 are considered as very high, high, low and very low, respectively. Among the 55 SSPs with retrieved structures, some albumins (ALB1, 7-15), all globulins, and most of glutelins (GLU1-5, 7, GLU9-16, GLU18-19) have high pLDDT scores ranging from 70-90, indicating the high reliability structures of the protein. Some of the albumins (ALB2-5, ALB16) and most of prolamins (PRO1-4, PRO6-9, PRO14-16, PRO18, PRO22-23, PRO25) have low pLDDT scores ranging from 50-70, indicating moderate reliability of these structures (Figure 7a, d, Table S13). One albumin protein, ALB6 and some of prolamins, PRO10-13, PRO17, PRO19-21, and PRO24 has very low pLDDT score which ranges between 40-50, indicating the presence of more disordered regions.

**Figure 7.**
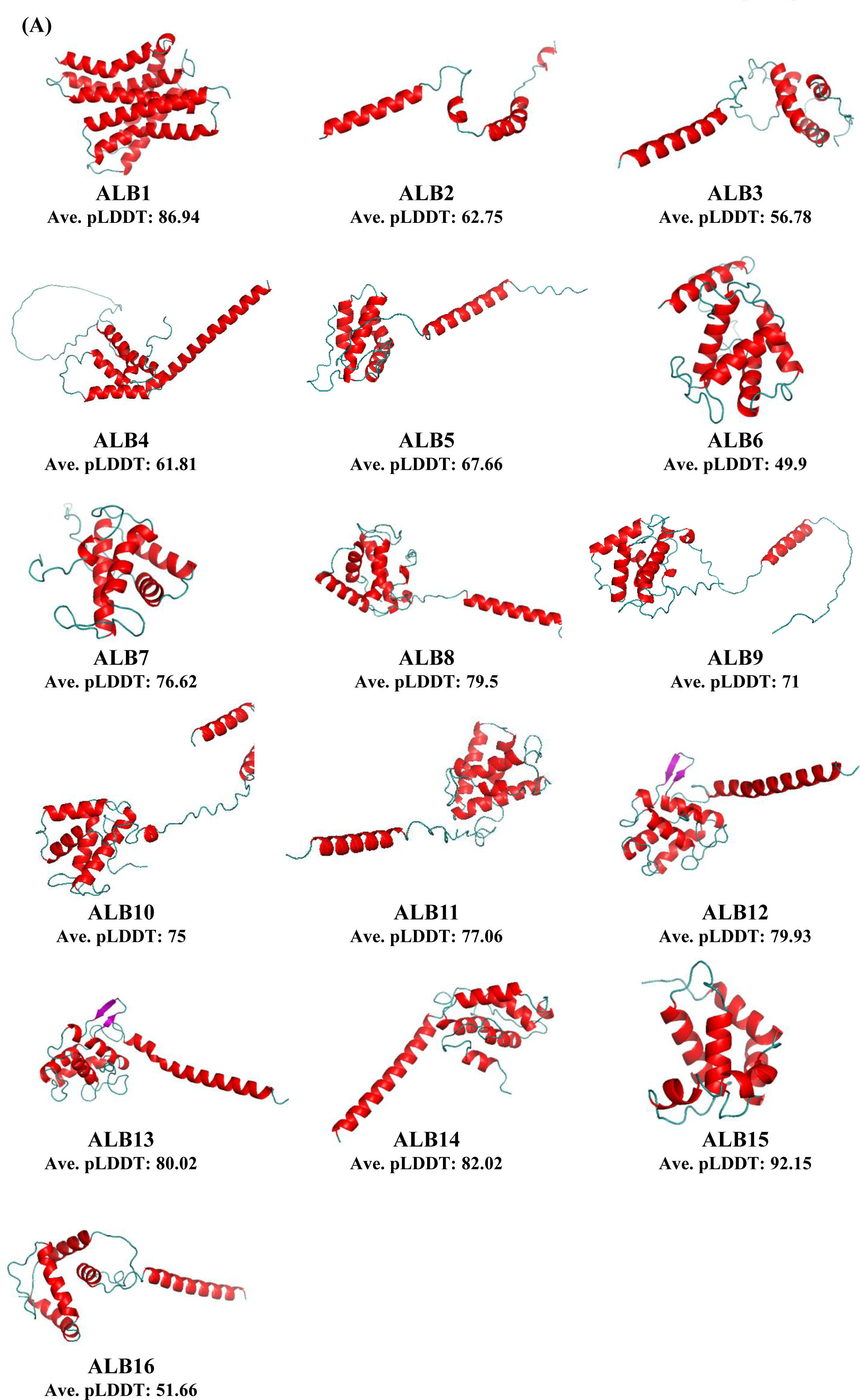
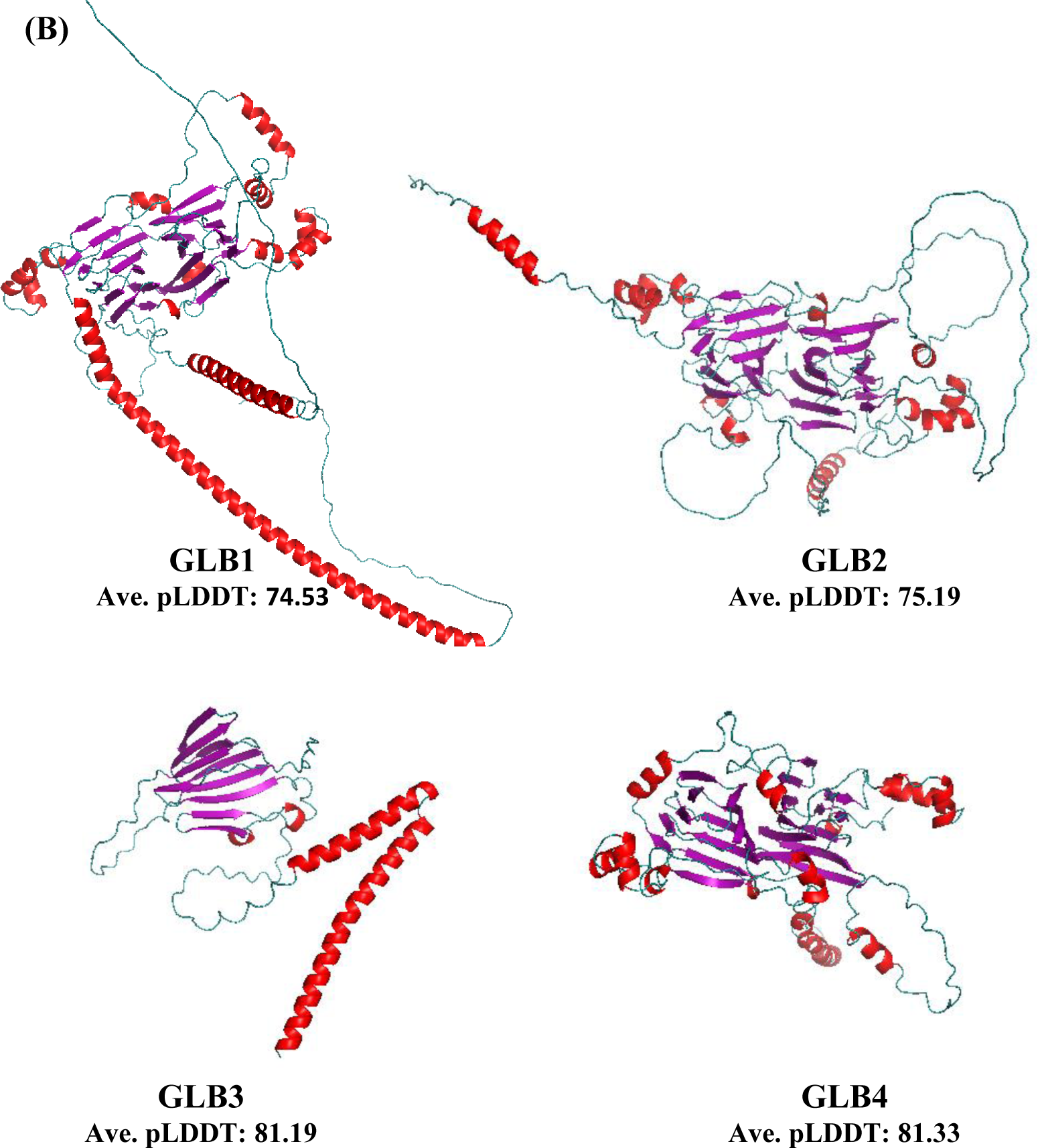
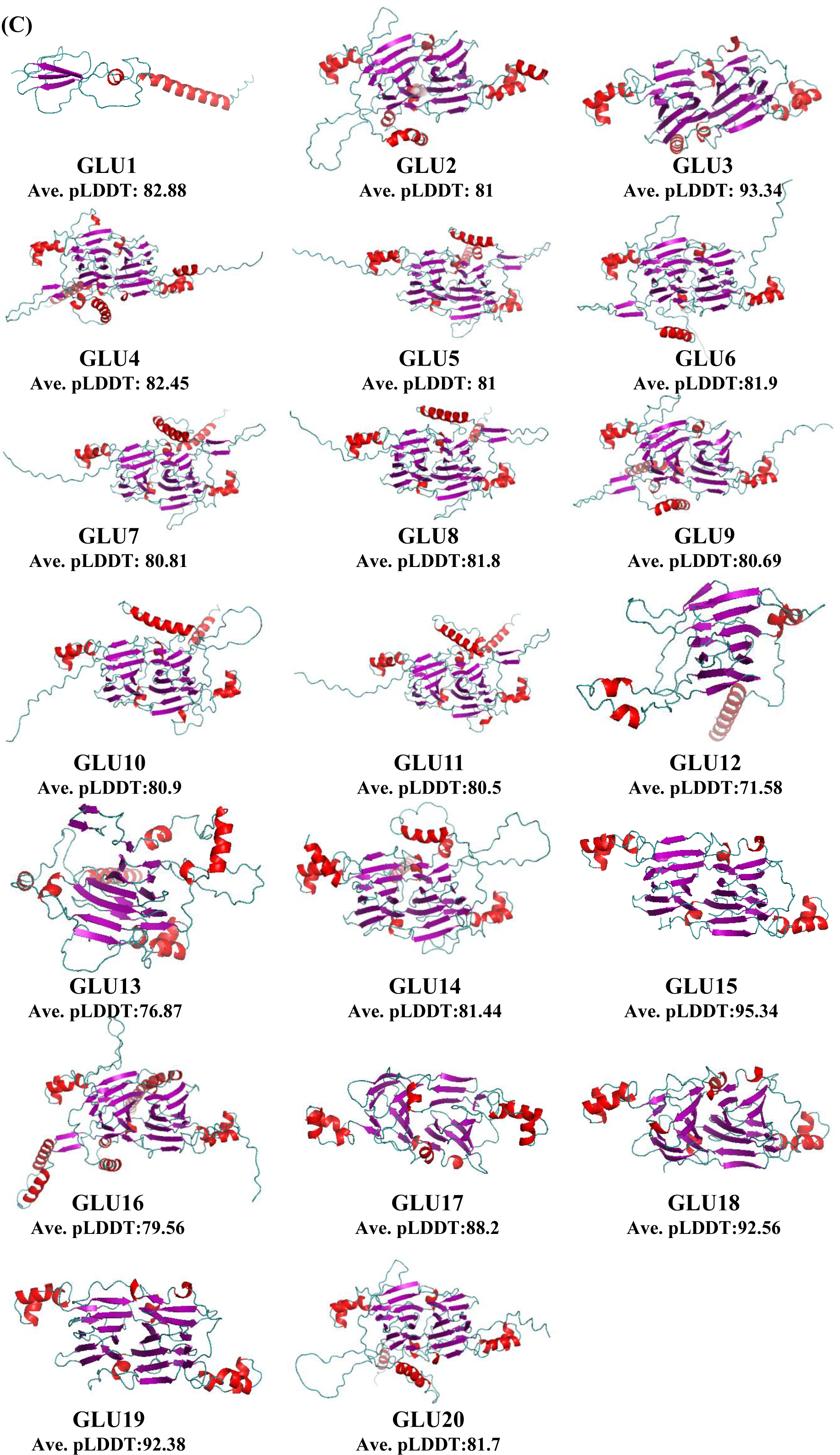
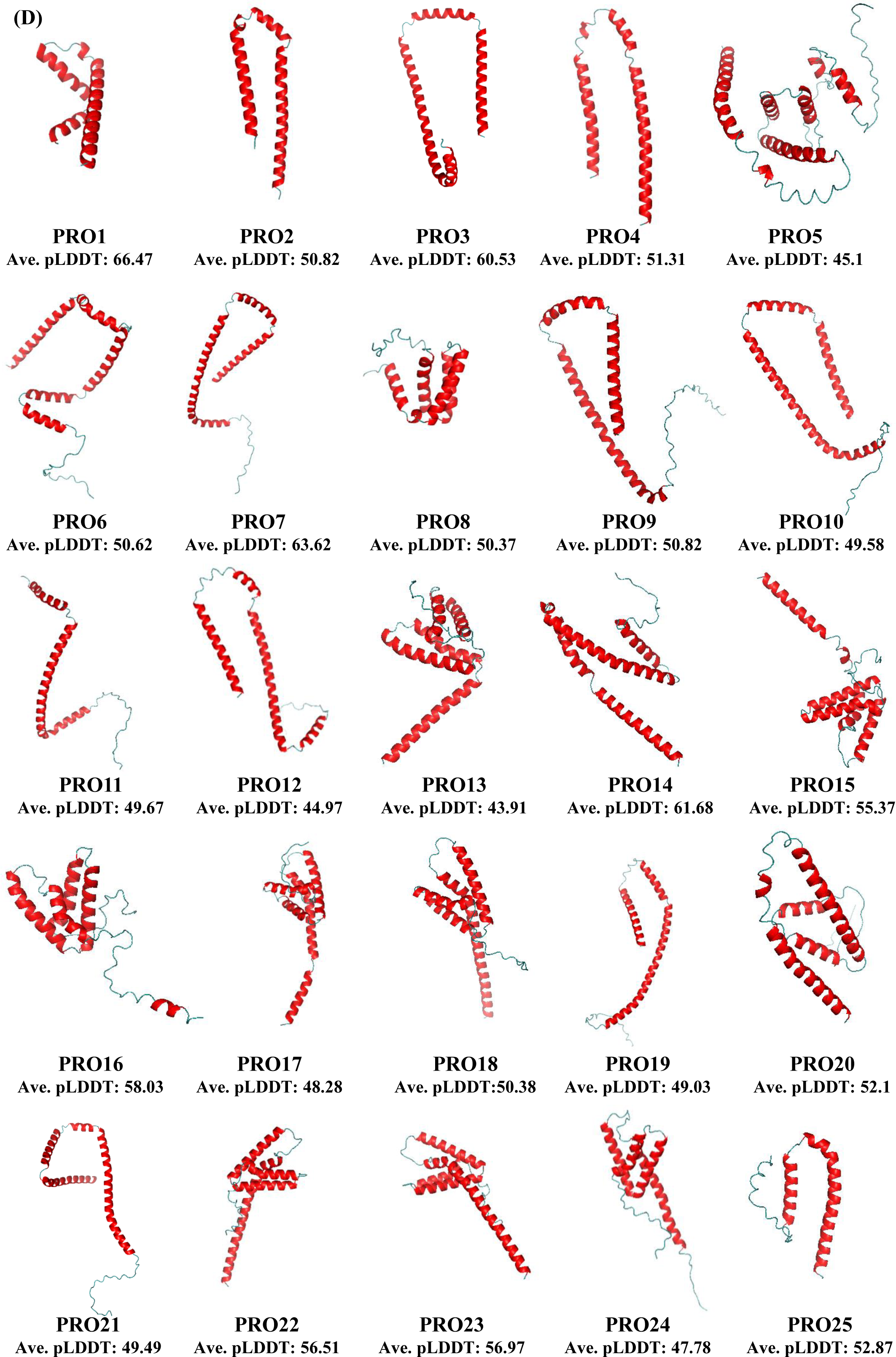
Tertiary structure models of different SSPs. The retrieved and predicted 3D structures of (A) albumins, (B) globulins, (C) glutelins and (D) prolamins are shown with their confidence scores. The helix region is shown in red and β-sheet is in purple. Here, ALBs: albumins, GLBs: globulins, GLUs: glutelins, PROs: prolamins.

For the proteins which did not have a structure available on AlphaFold Structure Database, 3D structures were predicted using abinitio method through AlphaFold Colab. Structures were predicted for three albumins (ALB8-10), five glutelins (GLU6, GLU8, GLU10, GLU17, and GLU20) and two prolamins (PRO5, PRO20). Five structures were generated for each protein with varying pLDDT values (Table S14). The optimal structure was chosen on the basis of highest pLDDT values. ALB8-10, GLU6, GLU8, GLU10, GLU17, and GLU20 have pLDDT values in the high range, which indicate the closeness of predicted structure with the experimentally determined structure. PRO20 has a low pLDDT score in range of 50-55, and PRO5 has very low pLDDT score of 45.1, which indicates less reliable structure due to more disordered regions in the protein.

3D structural analysis showed that each ALB belonged to the alpha group and contained the bifunctional inhibitor/seed storage helical domain (Figure 7a, Table S12). This domain is present in moderate positions of different albumins. It consisted of 4-6 conserved motifs, each composed of alpha helices, separated by a less conserved region composed of another two helixes with an intervening variable loop. Rice globulins revealed a highly conserved 3-D structure and belonged to alpha-beta mixed category of proteins. GLBs had two structurally similar units, each of which is formed of multiple β-sheets (Figure 7b, Table S12). All four globulins had a region showing a high resemblance to the β-barrel, as reported for other globulins. The β-barrel structure was formed by the cupin and RmlC domains, which overlapped with each other. The cupin domains were ∼115 residues long and RmlC domains were ∼200 residues long. This family of cupin domains represent the conserved barrel domain of the ’Cupin’ superfamily (’cupa’ is the Latin term for a small barrel). The cupin domain contained a six-stranded β barrel structure, which contains two conserved motifs, each corresponding to two beta strands separated by a less conserved region composed of another two beta strands with an intervening variable loop. At the higher structural level, rice glutelins shared close resemblance with globulins (Figure 7c, Table S12). Both are bicupin proteins with a core β-barrel confirmation. The only exception was GLU1 with a single cupin domain that showed the presence of a single β-barrel structure. The overlapping RmlC domain was formed from two beta-sheets arranged in a beta-sandwich. The length of Cupin 1, RmlC and 11s seed storage domains were ∼100, ∼200 and ∼24 residues, respectively (Table S12). Apart from β-strands, all GLU structural domains consisted of at least four helices with a folded leaf topology, forming a right-handed superhelix (Figure 7c, Table S12). As seen for the primary and secondary structures, the tertiary structure of prolamins showed homology to rice albumins (Figure 7d). Each PRO had repeated alpha helices separated by loop structures. Prolamins showed the presence of bifunctional trypsin and gliadin domain and both were present on overlapping positions contributing to the helices.

## DISCUSSION

The nutrient quality of cereals is largely dependent on starch and SSPs that are reserved as sources of carbohydrates and nitrogen, respectively (Cao et al., 2024). Research on SSPs in cereals is important since their composition and content affect the overall nutritional quality of seeds (He et al., 2021). Understanding the physical, functional, and biological properties of SSPs, which determine the biosynthesis, trafficking, and deposition of SSPs, will assist future attempts to boost grain quality through genetic engineering (Cao et al., 2024). In this study, we performed *in silico* analyses for the identification, characterization, phylogeny, expression and structure prediction, and of rice SSPs. A HMMER-based search in the rice genome identified a total of 65 SSPs, which included 16 albumins, four globulins, 20 glutelins, and 25 prolamins. Out of these, 10 albumins, 4 globulins, 5 glutelins were newly identified (Figure 1, Table S2). All four major categories of SSPs except globulins, undergo tandem duplication as shown by their close localization on the chromosomes (Table S2). Like in the case of prolamins, all 25 proteins were localized on four chromosomes i.e. 5, 6, 7, and 12 which suggested the involvement of inter-chromosomal duplication events that might have happened during the course of evolution. The expansion of the SSP gene family in rice and higher similarity among them is due to the occurrence of several duplication events that are also supported by a study in chickpea (Verma and Bhatia, 2019).

Domain analysis of the 16 rice albumins revealed their higher similarity to seed allergenic proteins and as well as to the lipid transfer proteins/LTPs, all these are considered as similar proteins (Mills et al., 2002; Jain, 2023). 2S albumins are a major source of seed storage proteins and are known to possess both nutritional and clinical properties. Reports suggest that transferring 2S albumin genes from different sources to legume seeds by genetic engineering, provides great nutritional value and also plays a protective role against fungal attack (Moreno and Clemente, 2008; Platani et al., 2023). Albumins and prolamins both shared the 2S albumin domain, motifs 9, 18 as well as CCxQL motif. CCxQL consensus motifs are known to form disulfide bonds with dicysteine residues and play an important role in targeting proteins into PBs (Kawagoe et al., 2005; Masumura et al., 2015). Domain and MEME analysis of glutelins and globulins revealed the presence of two cupin domains and motif 6, 24 in common. The bicupin proteins are also known to provide pathogenicity as well as nutritional value to seeds (El Hadrami et al., 2015). The sharing of motifs between albumin-prolamin and glutelin-globulin classes of SSPs suggested the similarity among the protein sequences of these classes in rice, although they have undergone significant diversification over time. (Figure 2, Figure S1). Most of the SSPs belonging to albumin, globulin, and glutelin classes had a proteolytic (N-G) cleavage site which suggests their synthesis as precursors initially. These precursor SSPs undergo proteolytic cleavage by asparaginyl endopeptidase (APE)/vacuolar processing enzymes (VPE)/legumin to form functional proteins (Figure 2), which has also been observed previously (Koziol et al., 2012; Nonis et al., 2021).

Further, phylogenetic analysis among rice SSPs and with SSPs from other species of *Poaceae* family showed that albumin and prolamin as well as glutelin and globulin arose from common ancestors (Figure 3, 4). Thus, strong conservation in the protein sequences as observed in domain and motif analysis was confirmed by the close phylogeny of these proteins. Phylogenetic studies of rice SSPs suggested that albumin proteins (ALB4-15) are the oldest (Figure 3 and Figure 4). Four albumins (ALB1, 2 3, and 16), showed more similarity with the prolamins, suggesting their evolution possibly from prolamins. The globulin and glutelin proteins formed an entirely separate clade in the tree and could be clearly divided into the 11S-12S clade and the 7S-9S globulin clade. All rice glutelins belonged to the 11S-12S group, whereas the four rice globulins belonged to the 7S-9S group. Evolutionary studies revealed sequence homology amongst 11S legumin and 7S vicilin proteins. Both these proteins have a common single-domain ancestor (Gibbs et al., 1989; Shutov et al., 1995). Both 7S and 11S globulin proteins are known to have an equivalent N-terminal and C-terminal region which forms β-barrel shaped cupin domain, despite their relatively lesser sequence homology (Mills et al., 2002; Jain, 2023).

Analyzing *cis*-regulatory elements within gene promoters is essential for understanding gene function and for engineering promoters with tissue-specific activity (Xu et al., 2024). Seed-specific promoters, commonly derived from genes active during seed development, have diverse applications, including targeted expression of industrial and pharmaceutical products, as well as improving seed nutritional and processing qualities (Hernandez-Garcia et al., 2014). Promoters from cereal storage protein genes like *HORDEIN* and *GLUTENIN* have been successfully used for producing recombinant proteins in seeds (Kawakatsu et al., 2010). Since seeds of crops like rice, wheat, corn, and legumes are major global food sources, enhancing their value using specific promoters is particularly valuable. In monocots, storage reserves accumulate in the endosperm, making endosperm-specific promoters highly useful. These promoters have been used to express proteins such as PRO-INSULIN, LACCASE, Β-CAROTENE, and viral antigens in seeds (Chen et al., 2007; Sharma et al., 2009). *In silico* analysis of the upstream 2 kb promoter region of each SSP showed the presence of different seed-specific *cis*-regulatory elements like 2SSEEDPROTBANAPA, CAATBOX1, different GCN motifs, DOFCOREZM, TATABOX, EBOXBNNAPA, RYREPEATBNNAPA, SEF4MOTIFGM7S, SKN-1-LIKE etc. (Figure 5, Table S4). Similar results have been observed in a recent study in which the promoters of barley and rice orthologs of *TaMFT-3A* contain seed-specific *cis*-elements such as the RY motif, CCAAT box, A-box, and G-box, which drive predominant expression in the scutellum (Utsugi et al., 2025). Similarly, the sesame *FAD2* promoter harbors multiple seed- and endosperm-specific motifs, including GCN4, ACGT, AACA, E-boxes, Dof cores, RY repeats, ABRE, and G-box elements, supporting its role in seed-specific expression (Kim et al., 2006). Sequence difference among the promoter regions of *SSP*s led to the differential abundance of *cis*-elements that might have contributed to their differential expression among the seed developmental stages (Table S4-7). These elements have previously been suggested to bind different transcription factors and establish their role in seed development. For instance, in maize, coordinated activation of *ZEIN* genes involves the interaction of the prolamin box (P-box) with endosperm-specific transcription factors like zinc finger and bZIP proteins (Vicente-Carbajosa et al., 1997), while the ESP element contributes to strong endosperm-specific activity in globulin promoters (Vickers et al., 2006), CAATBOX and TATABOX are considered as important regulatory elements of the core-promoter region affecting the transcriptional efficiency (Marand et al., 2023). DOFCOREZM elements were present in multiple copies in various *SSPs*, these DOF TFs specifically bind to DOF core elements and regulate their tissue-specific expression (Kawakatsu et al., 2010; Yang et al., 2018). The soybean TFs, GmDOF4 and GmDOF11 bind to the DOF core element and increase the fatty acid and lipid content in transgenic *Arabidopsis* seeds (Wang et al., 2007; Sun et al., 2018). Specific binding of RPBF activator to DOF core elements results in increased protein accumulation in rice endosperm (Kawakatsu and Takaiwa, 2010). E-box elements present in SPATULA (SPT) and ALCATRAZ (ALC) are involved in flower and fruit development (Cheng et al., 2022). RY element is enriched in many endosperm-specific genes in rice and also enhances the expression of globulin proteins in soyabean seeds (Fujiwara and Beachy, 1994; Nie et al., 2013). They also confer a seed-specific expression of *PvALF/Phaseolus vulgaris ABI3-LIKE FACTOR* (Nie et al., 2013). The SEF4MOTIFGM7S is shown to be the binding site for SOYBEAN EMBRYO FACTOR/SEF and regulates the expression of α-subunit of β-conglycinin *SSP* (De Silva et al., 2022). SKN-1-LIKE elements also play an important role in guiding the endosperm-specific expression of seed storage proteins as of rice glutelin gene *GluB-1* (Jin et al., 2019). Promoter analysis revealed the biased positioning of *cis*-elements, as they were mostly present on the transcribed positive strand or sense but some were specific to the antisense strands of DNA. A similar biased *cis*-regulatory elements (CRE) distribution was also observed in rice sperm cells where the number of CREs was more on the antisense strand as compared to the sense strand (Sharma et al., 2011).

The expression profiling of 65 *SSP* genes at five seed developmental stages suggested their important role in rice seed development. However, the expression pattern of *SSP*s varied within the seed developmental stages. Higher level of *SSP* transcripts in S2-S5 stages of seed development correlates with its considerable accumulations in the total protein in rice seeds and also indicates its role in seed germination and as a storage reserve of nitrogen, sulphur, and carbon for the growing seedlings (Figure 6) (Mondal et al., 2022; Kawakastu et al., 2010)). In rice, higher expression levels of prolamins in later stages of seed development focus its involvement in grain filling and seed maturation. Similar to this, the major *Arabidopsis SSPs* 12S globulins and 2S albumins accumulate during the mid- to late-stages of embryogenesis and are involved in seed maturation (Fujiwara et al., 2002; Yang et al., 2023). A similar difference in the pattern and level of expression among the *SSPs* was also observed in *Arabidopsis* where a hybrid promoter construct between the genes *at2S1* and *at2S2* resulted in the tissue-specific expression of 2S albumin (Da Silva Conceição and Krebbers, 1994). Hence, *SSPs* expression is tissue-specific and is transcriptionally regulated, and varies during the different seed developmental stages.

The 3D structure prediction of rice SSPs, through AlphaFold, provides valuable insights into structural reliability and protein stability. The high pLDDT values of rice SSPs structures, particularly in globulins and glutelins suggest these proteins possess reliable structures with stable conformations. The high confidence scores imply these proteins may play crucial roles in seed development, especially in nutrient storage. Some proteins possessed lower pLDDT values, which dictate the presence of disordered segments, particularly in albumins and prolamins. The presence of more flexible regions in albumins and prolamins indicates the functional adaptability of these proteins. This indicates the involvement of ligand-protein interactions under various physiological conditions. Proteins with very low confidence scores as observed for ALB6 and some prolamins like PRO10-13, PRO17, PRO19-21, PRO24, could undergo conformational changes in response to cellular environment.

Various albumin structures reported from different species like *Ricinus communis* (RicC3 protein) (Pantoja-Uceda et al., 2003), *Arachis hypogea* (Arh 2 and Arh6) (Lehmann et al., 2006; Mueller et al., 2011), sunflower (SESA2-1, SESA2-2 etc.) (Franke et al., 2016), albumin domain of sunflower preproalbumin (PawS1 and PawS2) (Franke et al., 2016), soyabean (Soy Al 1 and Soy Al 3) (Han et al., 2016) and secondary structure of Cor a 14 from hazelnut (Han et al., 2016), all showed 4-6 α helix domains arranged in a right handed superhelix with leaf folded topology, 4 disulphide bonds and some disordered loops. The present study showed similar results with the presence of bifunctional inhibitor/seed storage helical domain in all albumins showing 4-6 α helical conserved motifs, separated by two less conserved helical regions with an intervening loop (Figure 7a). The 3D structure of rice ALBs and from these different species also exhibited several structural variations with respect to number, position and length of their helix, interhelical angles and folding of loops, which indicate that rice ALBs belong to different classes. These structural differences may contribute to differences in the functions they perform.

All globulins showed the presence of two distinct domains, cupin 1 and RmlC domain. Cupin domain was composed of six stranded β barrel structures. The other domain with a double-stranded beta-helix jelly roll fold similar to that found in RmlC was also found in globulins (Figure 7b). The previous reports coincide with our findings, where different globulin protein stuctures like, structure of 8S alpha globulin of mung bean (*Vignaradiata* (L.)) (Itoh et al., 2006), β viginin of cowpea (Rocha et al., 2018), Bg7S from soyabean (Yoshizawa et al., 2011; Magni et al., 2018), *Wrightia tinctoria* 11S globulin (Kumar et al., 2017), SM80.1 protein from *Solanum melongena* (Jain et al., 2016) and 7S vicilins from olive (Jain et al., 2016), also showed the presence of two structurally similar units each made up of a core β-barrel and an extended α-helix domain. In case of globulins, the hypervariable loops mainly contribute to their structural and functional divergence in different species. The 3D structures which well characterize these hypervariable loops may give deeper insights into functional characteristics of different globulins.

Domain analysis showed the presence of two cupin 1 domain, RmlC domain and an 11S conserved site in each glutelin (Figure 2 and Supplemental table S12). Our study revealed the existence of helix, strands and coil region in secondary structure of glutelins that is in accordance with the previous studies which predict that secondary structure of glutelins is a combination of variable forms like helices, sheets, random coils and turns (Wang et al., 2023) (Figure S3d). Some studies have also shown the occurrence of beta turns or sheets and alpha helices in HMW and LMW glutelins respectively (Ortolan et al., 2022). Present study showed that all glutelin structural domains consisted of 4 helices with a folded leaf topology, and forming a right-handed superhelix, just like the one shown in wheat gluten proteins (Figure 7c), (Zhu et al., 2021).

Prolamins exhibited bifunctional trypsin and Gliadin domain (Figure 2, Table S12) mainly having 4-6 helix regions in their secondary structure (Figure S3d), similar to the structure reported for zeins (Zhang et al., 2018). Previous study in wheat has also shown the presence of equal number of α-helix, linear β structure, β turns and unordered structure in prolamins (Rani et al., 2021), whereas some observed the unordered prolamin structure with only small proportions of alpha helix and beta conformation (Hajjari and Sharif, 2024). Hence, structural differences along with some similarities of rice SSPs from different species put them into different classes and contribute to their functional divergence that still needs to be explored. As the available knowledge about the SSP encoding genes is inadequate, the present study reported the genome-wide identification and characterization of 65 SSPs in rice. This analysis has provided the comprehensive information of rice SSPs including gene expression and protein’s structural features, that can be further utilized for functional genomic studies and rice nutritional improvement.

## ACKNOWLEDGEMENTS

The authors thank the NBT e-library consortium for providing online access to research articles. IM, AY, AP acknowledges University Grants Commission for JRF and SRF fellowships. P.J. is thankful to Council of Scientific and Industrial Research, for JRF and SRF fellowships.

## AUTHORS CONTRIBUTIONS

P.A. arranged for funding, designed and supervised the analyses and experiments. AY, PJ, IEM and AP performed various analyses. All authors analyzed the data, prepared the figures, and wrote the article. All authors have read and approved the article.

