## Supplementary Figures for "Decoding Rice Seed Storage Proteins: From Gene Identification to Structural Prediction"

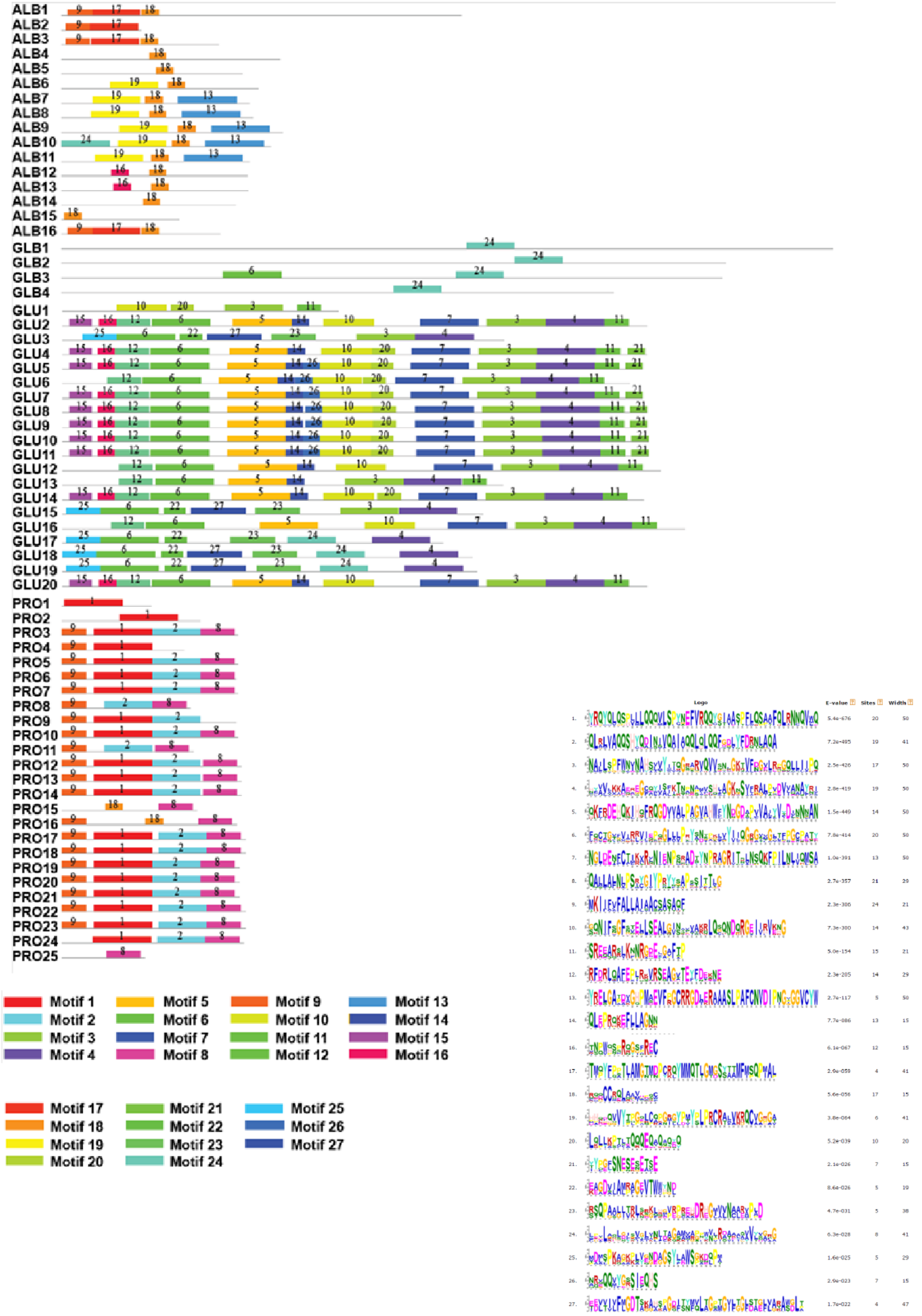


**Figure S1. Distribution of conserved motifs on SSPs by MEME analysis.** Different motifs highlighted in colored boxes numbered 1-27 are conserved among rice SSPs. The conserved sequence of each motif is shown on the right side. Gray line represents the non-conserved sequences present in the protein. The color legend for each motif is shown below the figure.

| | | | |
|---|---|---|---|
| ALB5 | GGDPPGVRAR | CCHQL | REVAARCRCD |
| ALB11 | AAAAEQVRRD | CCRQL | AAVDDSWCRC |
| ALB10 | GTVDEQVRRG | CCRQL | AAIDSSWCRC |
| ALB9 | GAVDEQLRQD | CCRQL | AAVDDSWCRC |
| ALB8 | SAADEQVWQD | CCRQL | AAVDDGWCRC |
| ALB7 | GAVDEQVRQD | CCRQL | AAIDDSFCRC |
| ALB6 | GNVDEQVWRD | CCRQL | ATINNNLCRC |
| ALB4 | RPGALGLRMQ | CCQQL | QDVSRECRCA |
| PRO24 | LRNCQVMQQQ | CCQQL | RMIAQQSHCQ |
| PRO23 | LRNCQVMQQQ | CCQQL | RMIAQQSHCQ |
| PRO22 | LRNCQVMQQQ | CCQQL | RMIAQQSHCQ |
| PRO18 | LINNQVMQQQ | CCQQL | RLVAQQSHYQ |
| PRO17 | LINNQVMQQQ | CCQQL | RLVAQQSHYQ |
| ALB16 | SQPMAPLQQQ | CCMQL | QGMMPQCNCD |
| ALB3 | SQPIALLQQQ | CCMQL | QGMIPQCHCG |
| ALB1 | SQPMALLQQQ | CCMQL | QGMMPQCHCG |
| ALB15 | MAKAR | CCREL | AAVQPRCRCE |
| ALB14 | RLYDWSLKER | CCQEL | AAVPAYCRCA |
| ALB13 | KQPPRQLLEP | CCREL | AAVPMQCRCD |
| ALB12 | KQPPREFLEP | CCREL | AAVPMQCRCD |
| PRO16 | LSSCQVMRQQ | CCQQM | RLMAQQYHCQ |
| PRO20 | HYQDINVVQA | IAHQL | HLQQFGNLYI |
| PRO19 | HYQDINVVQA | IAHQL | HLQQFGNLYI |
| PRO21 | HYQDINVVQA | IAQQL | HLQQFGDLYI |
| PRO14 | HYQDINIVQA | IAQQL | QLQQFDDLYF |
| PRO13 | HYEDINIFQV | IAQQL | QLQQFDDLYF |
| PRO12 | HYQDINIVQA | IAQQL | QLQQFDDLYF |
| PRO11 | HYQGINIVQA | IAQQL | QLQQFDDLYF |
| PRO9 | HYQDINIVQA | IAQQL | QLQQFGDLYF |
| PRO8 | HYQDINIVQA | IAQQL | QLQQFGDLYF |
| PRO7 | HYQDINIVQA | IAQQL | QLQQFGDLYF |
| PRO6 | HYQDINIVQA | IAQQL | QLQQFGDLYF |
| PRO5 | HYQDINIVQA | IAQQL | QLQQFGDLYF |
| PRO4 | HYQDINIVQA | IAQQL | QLQ |
| PRO3 | HYQDINIVQA | IAQQL | QLQQFGDLYF |
| PRO1 | HYQDINIVQA | IAQQL | |
| PRO25 | HCQAISSVLA | IVQQL | QLQQFAGVYF |
| PRO10 | HYQDINIVQA | IVQQL | QLQQFGDLYF |
| PRO15 | LSSCQVMRQQ | CCRRM | RLMAQQYRCQ |
| ALB2 | PTIAMGNMDP | CRQYM | MQTTGTDSYA |

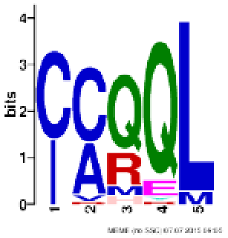


**Figure S2. Consensus motif sequence in albumins and prolamins by MEME analysis.** Presence of consensus motif CCxQL in most albumin and prolamin proteins is shown.

(A)

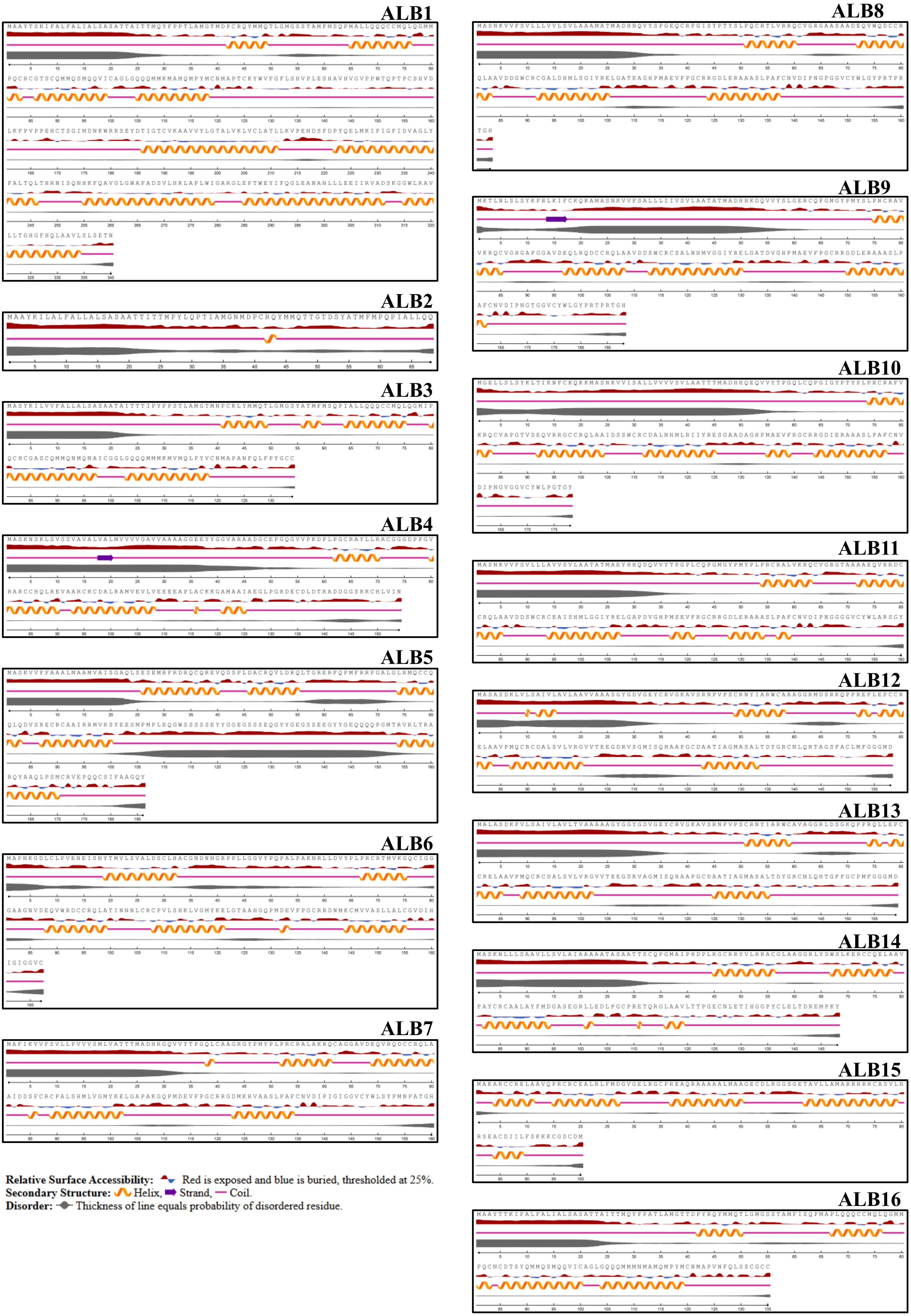
ALB1
ALB2
ALB3
ALB4
ALB5
ALB6
ALB7
ALB8
ALB9
ALB10
ALB11
ALB12
ALB13
ALB14
ALB15
ALB16
Relative Surface Accessibility: Red is exposed and blue is buried, thresholded at 25%.
Secondary Structure: Helix, Strand, Coil.
Disorder: Thickness of line equals probability of disordered residue.

(B)

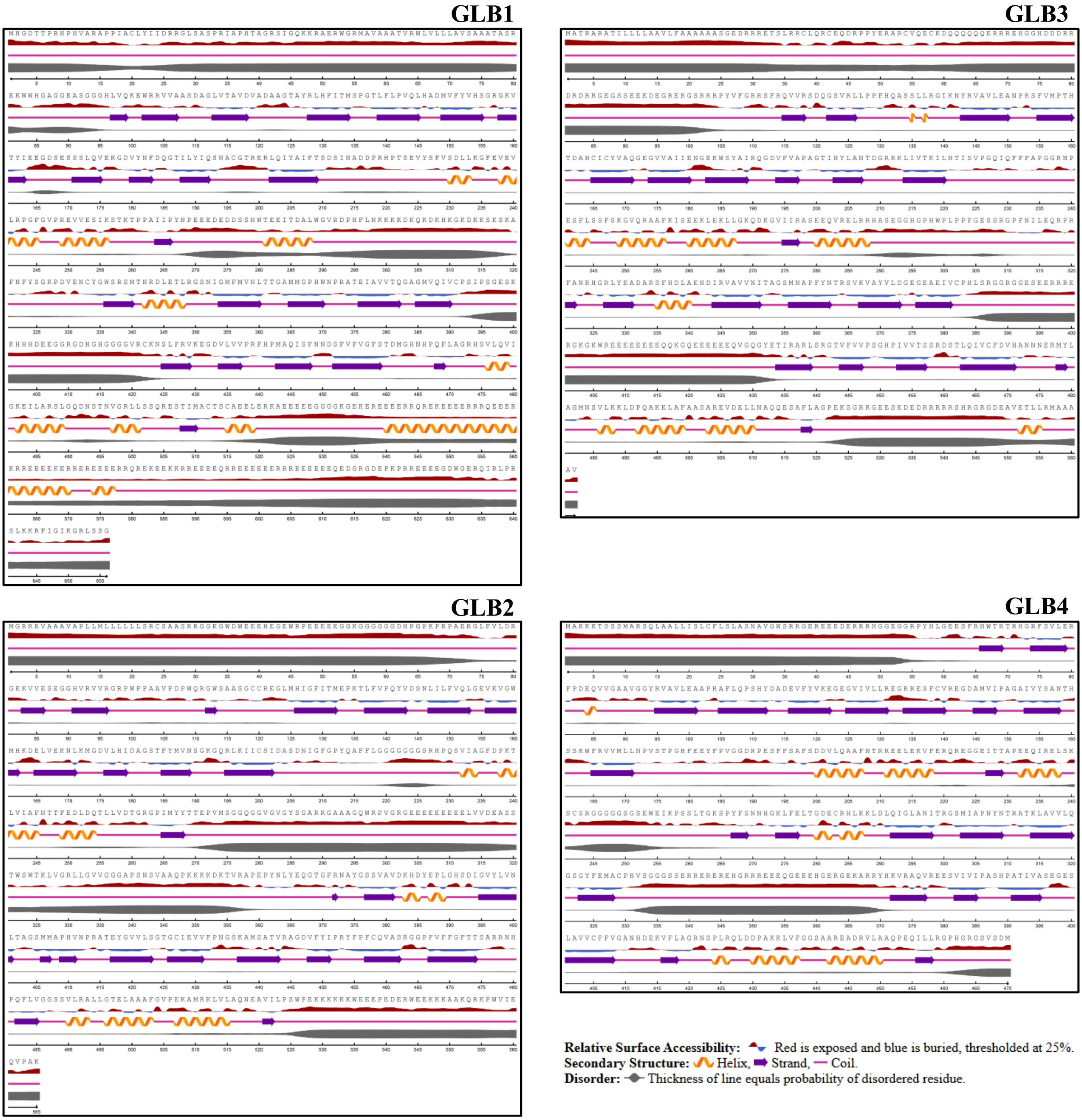

GLB1
GLB3
GLB2
GLB4
Relative Surface Accessibility: Red is exposed and blue is buried, thresholded at 25%.
Secondary Structure: Helix, Strand, Coil.
Disorder: Thickness of line equals probability of disordered residue.

(C)

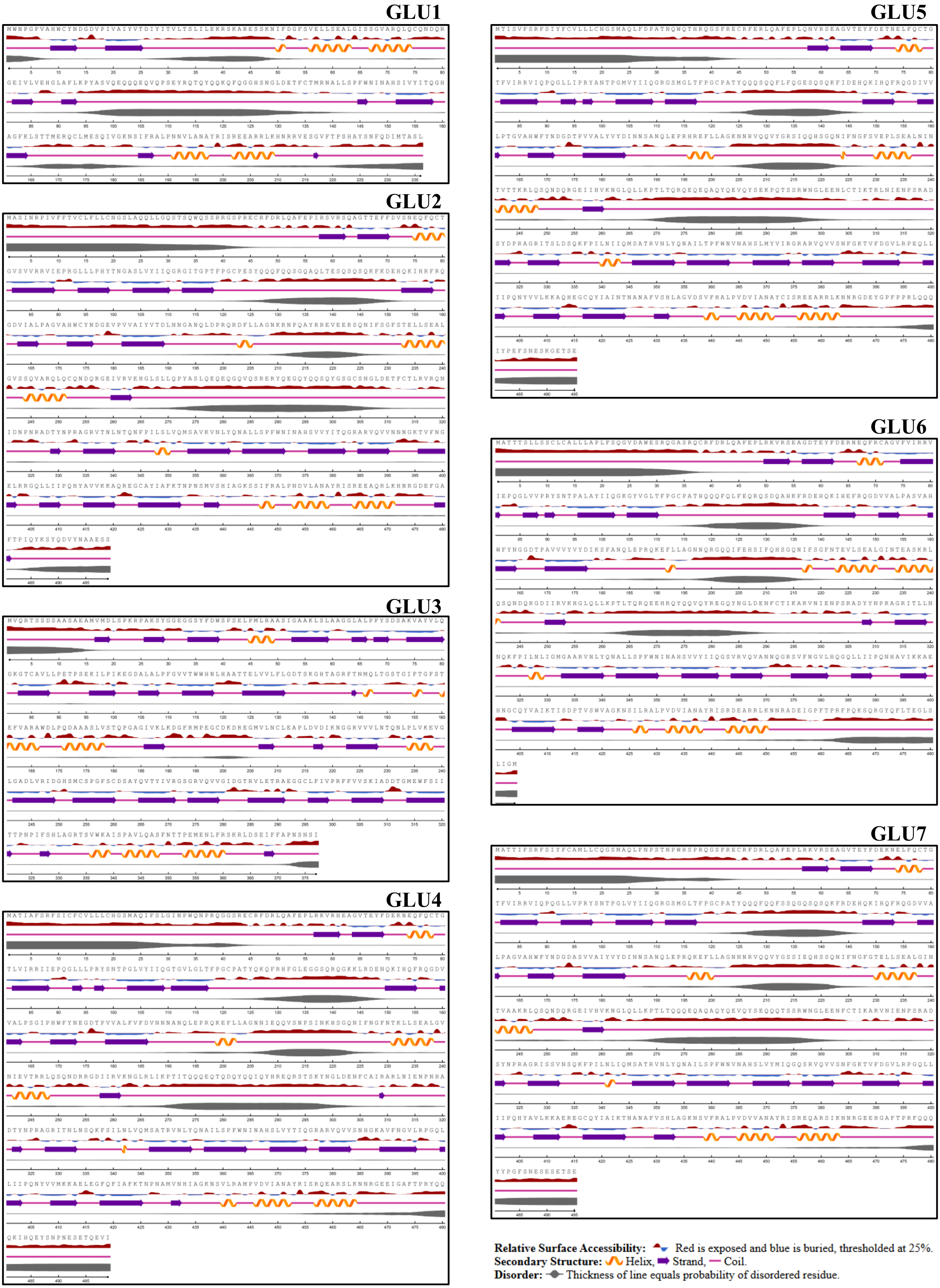


**Relative Surface Accessibility:** Red is exposed and blue is buried, thresholded at 25%.
**Secondary Structure:** Helix, Strand, Coil.
**Disorder:** Thickness of line equals probability of disordered residue.

**GLU8**

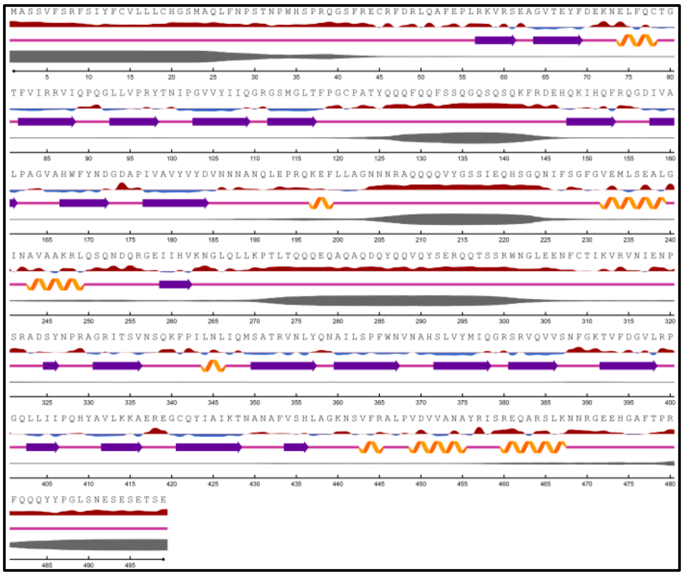

**GLU11**

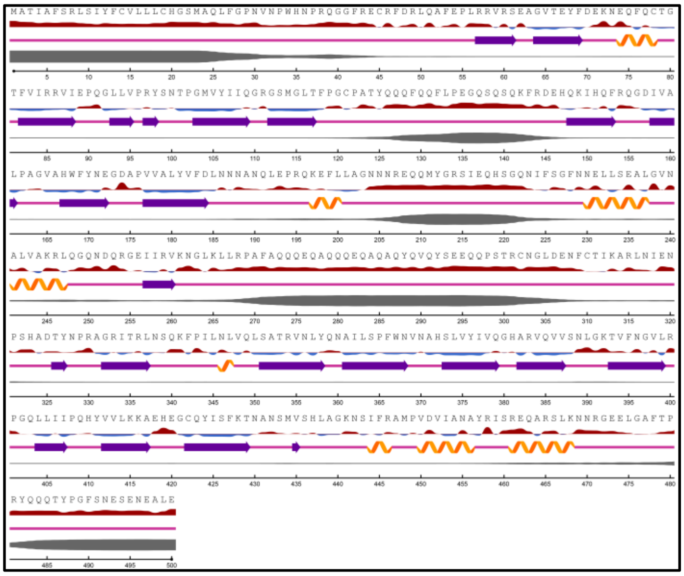

**GLU9**

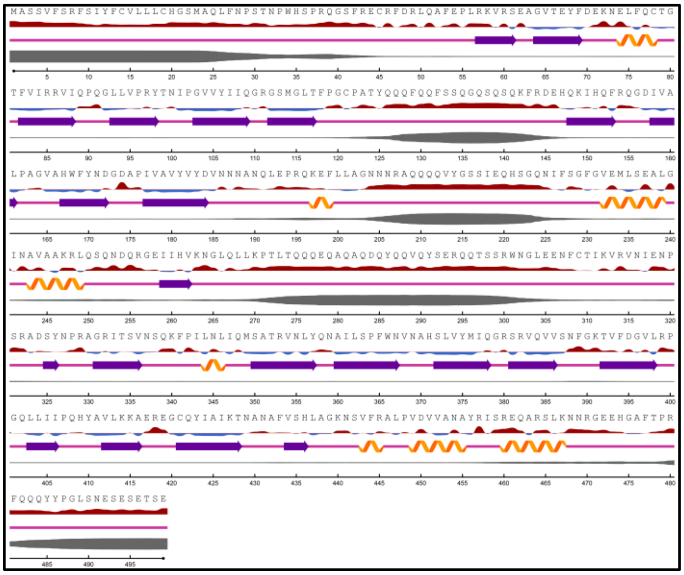

**GLU12**

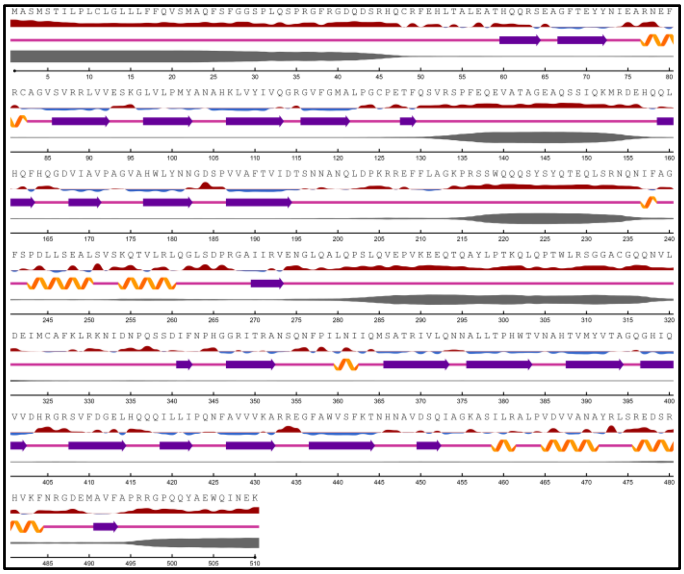

**GLU10**

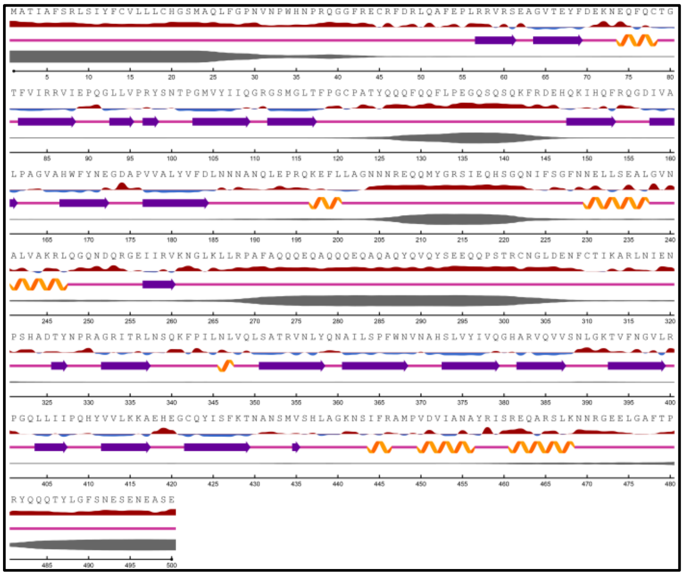

**GLU13**

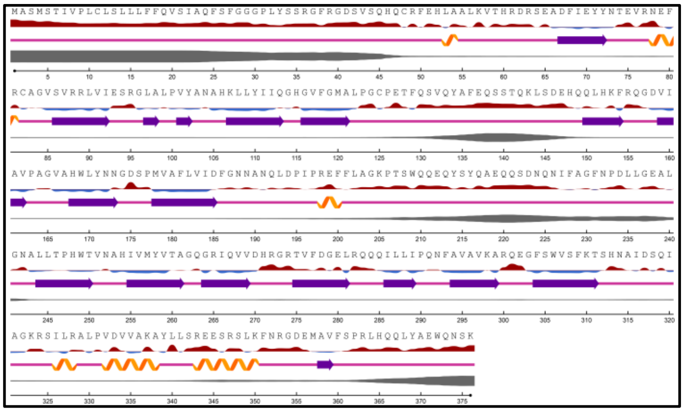

**Relative Surface Accessibility:** Red is exposed and blue is buried, thresholded at 25%.
**Secondary Structure:** Helix, Strand, Coil.
**Disorder:** Thickness of line equals probability of disordered residue.

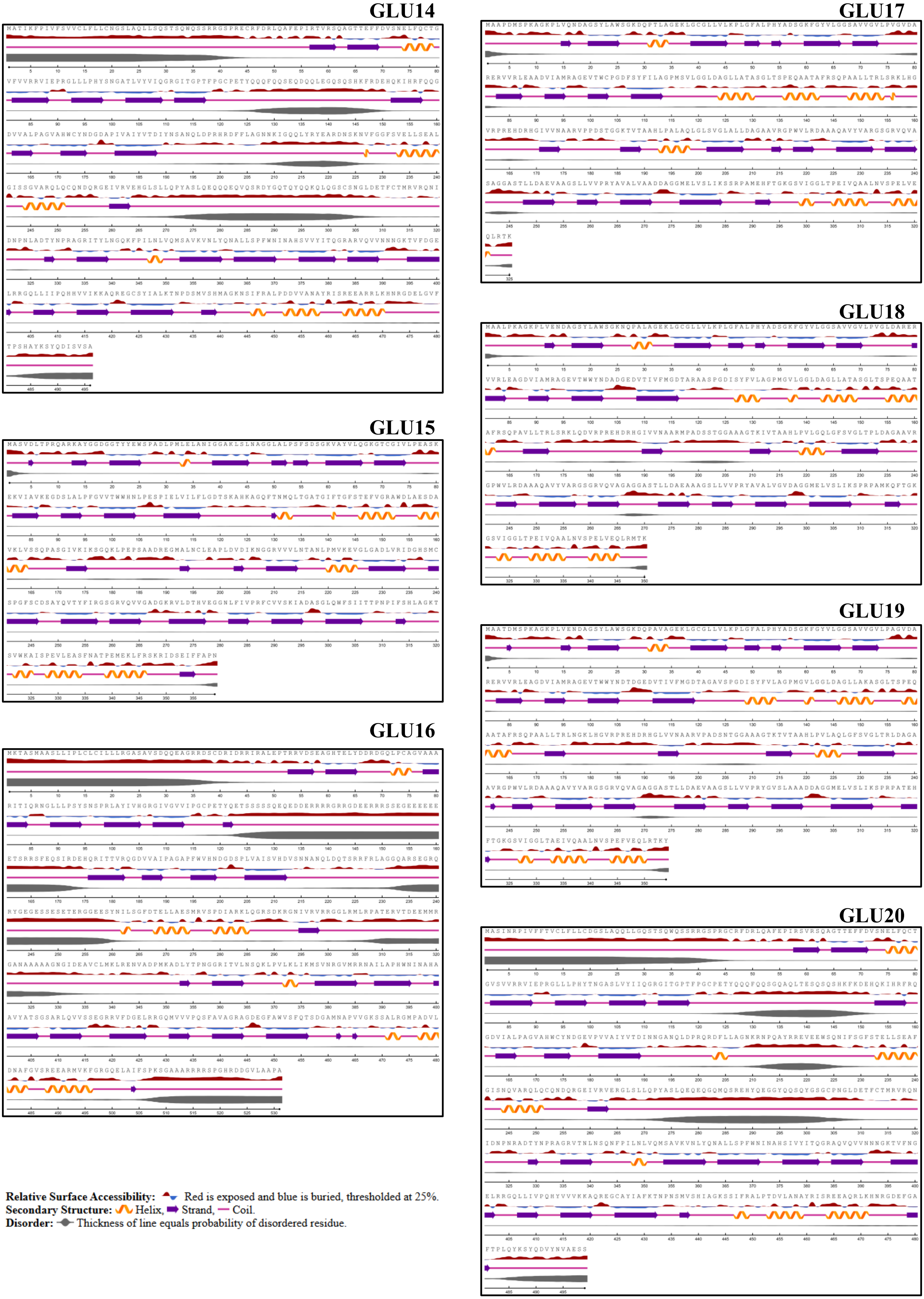
GLU14
GLU15
GLU16
GLU17
GLU18
GLU19
GLU20
Relative Surface Accessibility: Red is exposed and blue is buried, thresholded at 25%.
Secondary Structure: Helix, Strand, Coil.
Disorder: Thickness of line equals probability of disordered residue.

(D)

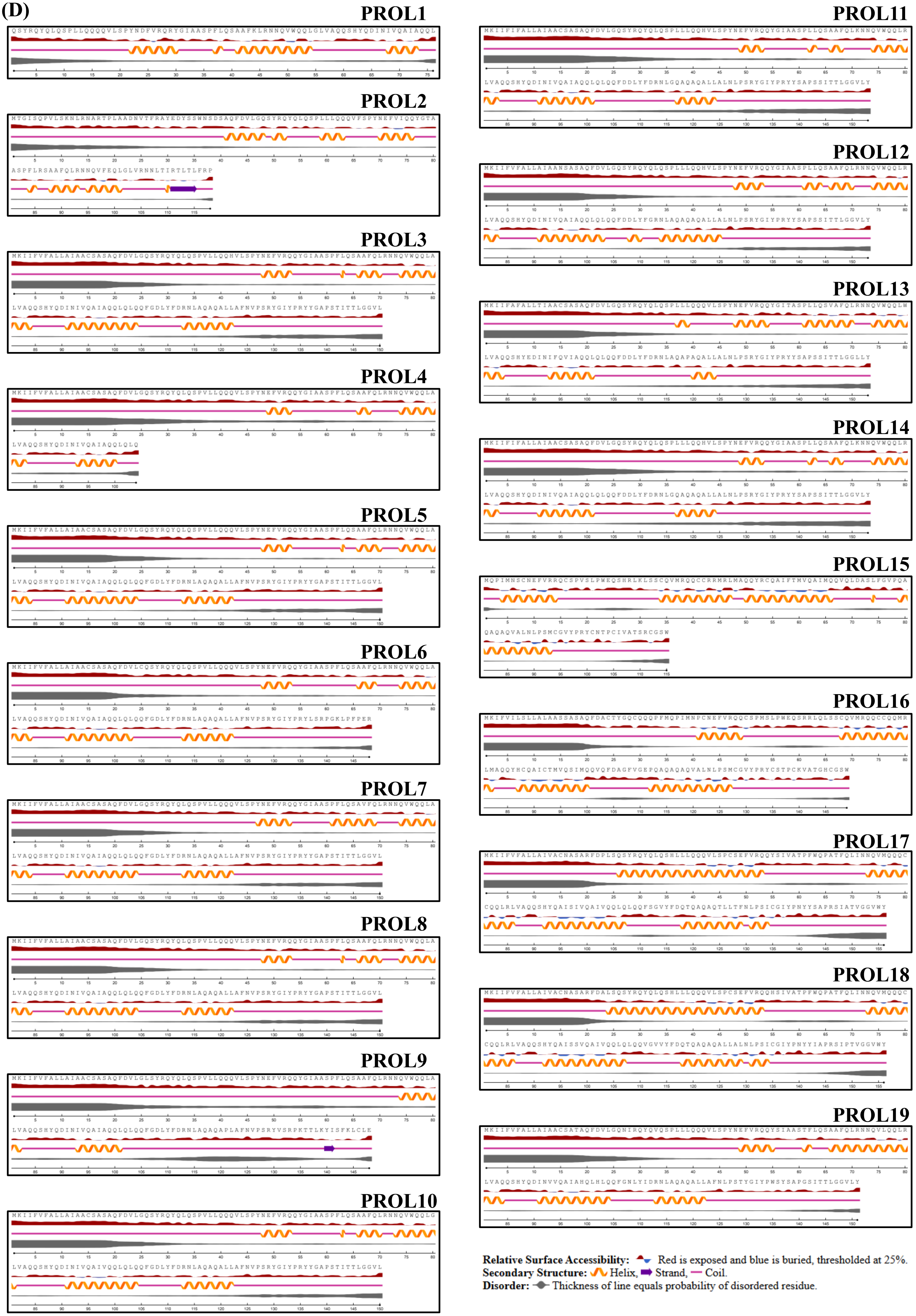
PROL1
PROL2
PROL3
PROL4
PROL5
PROL6
PROL7
PROL8
PROL9
PROL10
PROL11
PROL12
PROL13
PROL14
PROL15
PROL16
PROL17
PROL18
PROL19
Relative Surface Accessibility: Red is exposed and blue is buried, thresholded at 25%.
Secondary Structure: Helix, Strand, Coil.
Disorder: Thickness of line equals probability of disordered residue.

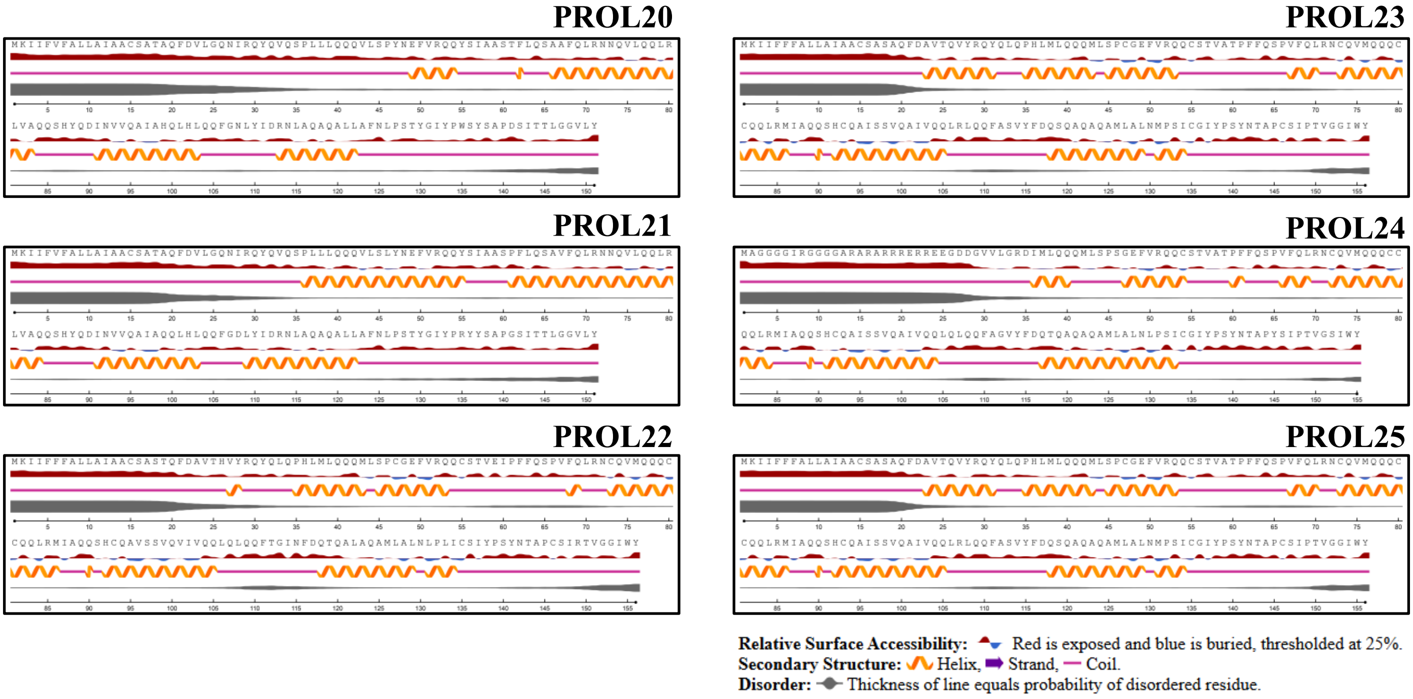


**Figure S3. Predicted secondary structures of rice SSPs.** The predicted secondary structures of (A) albumins, (B) globulins, (C) glutelins, and (D) prolamins are depicted. Helix is shown in orange, strand is shown as a purple arrow and pink line represents coil. Due to the varying lengths of the proteins, the structures are displayed across multiple horizontally aligned segments, with each segment including a scale bar at the bottom indicating the amino acid positions. At the top of each image, the amino acid sequence is shown, accompanied by red and blue peaks beneath it representing exposed and buried residues, respectively. Below the structural features, a grey line indicates the predicted disorder probability, with line thickness correlating to the likelihood of disorder, thicker regions suggest higher probability of disordered residues.
