## Supplementary Tables for "Decoding Rice Seed Storage Proteins: From Gene Identification to Structural Prediction"

**Table S1. List of accessions used for phylogenetic analysis within the grass family.**

| Sl. No. | Name given in the tree | Protein group | Acession ID | Species | Phylogenetic Group | Protein name as given in database |
|---|---|---|---|---|---|---|
| 1 | Alpha zein1 Z.m | Prolamins | NP_001105747.1 | *Zea mays* | Group I | 22kD alpha zein |
| 2 | Beta zein1 Z.m | Prolamins | NP_001106004.1 | *Zea mays* | Group II | 15kD beta zein |
| 3 | Gamma zein1 Z.m | Prolamins | ABD63258.1 | *Zea mays* | Group III | 16kD gamma zein |
| 4 | Gamma zein2 Z.m | Prolamins | AAL16979.1 | *Zea mays* | Group III | 50kD gamma zein |
| 5 | Delta zein1 Z.m | Prolamins | AAL16982.1 | *Zea mays* | Group I | 10kD delta zein |
| 6 | Delta zein2 Z.m | Prolamins | AAL16981.1 | *Zea mays* | Group I | 18kD delta zein |
| 7 | Alpha kafirin1 S.b | Prolamins | ACA28705.1 | *Sorghum bicolor* | Group I | alpha kafirin |
| 8 | Alpha kafirin2 S.b | Prolamins | ABP64793.1 | *Sorghum bicolor* | Group I | 19kD alpha kafirin B1 |
| 9 | Beta kafirin1 S.b | Prolamins | CAG30668.1 | *Sorghum bicolor* | Group II | beta kafirin |
| 10 | Beta kafirin2 S.p | Prolamins | ADD98897.1 | *Sorghum propinquum* | Group II | beta kafirin |
| 11 | Gamma kafirin1 S.p | Prolamins | ADD98901.1 | *Sorghum propinquum* | Group III | gamma kafirin |
| 12 | Gamma kafirin2 S.b | Prolamins | AAS73289.1 | *Sorghum bicolor* | Group III | gamma kafirin protein |
| 13 | Delta kafirin1 S.b | Prolamins | AAW32936.1 | *Sorghum bicolor* | Group I | delta kafirin |
| 14 | Delta kafirin2 S.p | Prolamins | ADD98903.1 | *Sorghum propinquum* | Group I | delta kafirin |
| 15 | Alpha setarin1 S.i | Prolamins | Si028022m | *Setaria italica* | Group I | alpha zein |
| 16 | Alpha setarin2 S.i | Prolamins | Si027308m | *Setaria italica* | Group I | alpha zein |
| 17 | Gamma setarin1 S.i | Prolamins | Si032320m | *Setaria italica* | Group III | gamma zein |
| 18 | Gamma hordein1 B.d | Prolamins | Bradi2g38530.1 | *Brachypodium distachyon* | Group III | |
| 19 | Gamma hordein2 B.d | Prolamins | Bradi1g50290.1 | *Brachypodium distachyon* | Group III | GAMMA HORD |
| 20 | Delta setarin1 S.i | Prolamins | Si026997m | *Setaria italica* | Group I | delta zein |
| 21 | LMW Glu B.d | Prolamins | Bradi2g33280.1 | *Brachypodium distachyon* | Group III | LMW Glut |
| 22 | HMWglu B.d | Prolamins | Bradi2g20910.1 | *Brachypodium distachyon* | Group IV | HMW glut |
| 23 | D hordein1 B.d | Prolamins | Bradi2g20870.1 | *Brachypodium distachyon* | Group IV | D HORD |
| 24 | D hordein2 B.d | Prolamins | Bradi2g20860.1 | *Brachypodium distachyon* | Group IV | D HORD |
| 25 | Gamma gliadin1 T.a | Prolamins | ABO37961.1 | *Triticum aestivum* | Group III | gamma gliadin2 |
| 26 | Omega gliadin1 T.a | Prolamins | BAE20328.1 | *Triticum aestivum* | Group III | omega-5 gliadin |
| 27 | B hordein1 H.v | Prolamins | CAA37729.1 | *Hordeum vulgare subsp. Vulgare* | Group III | B hordein precursor |
| 28 | C hordein1 H.v | Prolamins | AAA92333.1 | *Hordeum vulgare subsp. Vulgare* | Group III | C hordein |
| 29 | D hordein1 H.v | Prolamins | AAP31051.1 | *Hordeum vulgare* | Group IV | D-Hordein |
| 30 | HMW glutenin1 T.a | Prolamins | ABG68035.1 | *Triticum aestivum* | Group IV | Y-type HMW glutenin |
| 31 | LMW glutenin1 T.t | Prolamins | ABO37957.1 | *Triticum turgidum* | Group III | low-molecular-weight glutenin GLU-B3 |
| 32 | Alpha gliadin2 D.h | Prolamins | ABY48881.1 | *Dasypyrum hordeaceum* | Group III | alpha-gliadin |
| 33 | Gamma gliadin2 T.a | Prolamins | AAA34272.1 | *Triticum aestivum* | Group III | gamma gliadin precursor |
| 34 | Gamma hordein2 H.v | Prolamins | AFM77739.1 | *Hordeum vulgare* | Group III | gamma 3 hordein |
| 35 | B hordein2 H.v | Prolamins | ABB82614.1 | *Hordeum vulgare subsp. vulgare* | Group III | B hordein |
| 36 | C hordein2 H.v | Prolamins | AFM77749.1 | *Hordeum vulgare subsp. vulgare* | Group III | C hordein |
| 37 | D hordein2 H.v | Prolamins | BAA11642.1 | *Hordeum vulgare subsp. vulgare* | Group IV | D hordein |
| 38 | HMW glutenin2 T.a | Prolamins | AAA62315.1 | *Triticum aestivum* | Group IV | high-molecular-weight glutenin |
| 39 | LMW glutenin 3 T.a | Prolamins | AAP44989.1 | *Triticum aestivum* | Group III | low molecular weight glutenin |
| 40 | LMW glutenin4 T.a | Prolamins | AAS10193.1 | *Triticum aestivum* | Group III | low molecular weight glutenin |
| 41 | Glb1 Z.m | Protease inhibitor/seed storage protein/L | AAL16993.1 | *Zea mays* | Group IV | alpha globulin |
| 42 | Glb2 Z.m | Protease inhibitor/seed storage protein/L | NP_001105060.1 | *Zea mays* | Group IV | globulin3 |
| 43 | 7S glb Z.m | 7S globulin | AAA33467.1 | *Zea mays* | Glutelins and Globulins | globulin precursor |
| 44 | Legumin Z.m | 11S globulin | AAO63625.1 | *Zea mays* | Glutelins and Globulins | legumin-like protein |
| 45 | 7S glb1 S.i | 7S globulin | XP_004971203.1 | *Setaria italica* | Glutelins and Globulins | basic 7S globulin2 |
| 46 | 7S glb2 S.i | 7S globulin | XP_004971200.1 | *Setaria italica2* | Glutelins and Globulins | basic7S globulin2-like |
| 47 | 13S glb S.i | 11S globulin | XP_004980234.1 | *Setaria italica* | Glutelins and Globulins | 13S globulin |
| 48 | 7S glb1 B.d | 7S globulin | XP_003568479. | *Brachypodium distachyon* | Glutelins and Globulins | basic 7S globulin-like |
| 49 | 12S glb B.d | 11S globulin | XP_003576425.1 | *Brachypodium distachyon* | Glutelins and Globulins | 12S seed storage globulin1-like |
| 50 | 11S glb B.d | 11S globulin | XP_003571555.1 | *Brachypodium distachyon* | Glutelins and Globulins | 11S globulin seed storage protein |
| 51 | 19kDa glb2 T.a | Alpha amylase inhibitor/ seed storage pro | AAP80642.1 | *Triticum aestivum* | Group IV | 19kDa globulin |
| 52 | Glb1T.t | Alpha amylase inhibitor/ seed storage pro | AAR95703.1 | *Triticum turgidum* | Group IV | globulin |
| 53 | 7S glb1 T.u | 7S globulin | EMS63957.1 | *Triticum urartu* | Glutelins and Globulins | Basic 7S globulin2 |
| 54 | 7S glb2 T.u | 7S globulin | EMS50909.1 | *Triticum urartu* | Glutelins and Globulins | Basic 7S globulin2 |
| 55 | Triticin1 T.a | 11S globulin | AAB27108.2 | *Triticum aestivum* | Glutelins and Globulins | triticin precursor |
| 56 | 12S glb T.u | 11S globulin | EMS60030.1 | *Triticum urartu* | Glutelins and Globulins | 12S seed storage globulin1 |
| 57 | Glb1 B.d | Protease inhibitor/seed storage protein/L | Bradi2g20860.1 | *Brachypodium distachyon* | Group IV | |
| 58 | Glb1 S.i | Protease inhibitor/seed storage protein/L | Si023413m | *Setaria italica* | Group IV | |
| 59 | Glb2 S.i | Protease inhibitor/seed storage protein/L | Si032232m | *Setaria italica* | Group IV | |
| 60 | Glb3 S.i | Alpha amylase inhibitor/ seed storage pro | Si031402m | *Setaria italica* | Group IV | |
| 61 | 12S glb1 S.b | 11S globulin | Sobic.009G007100 | *Sorghum bicolor* | Glutelins and Globulins | 12S Globulin |

Based on the availability of sequences in the public databases, two proteins belonging to each category within a tribe have been analyzed

**Table S2. Details of SSP encoding genes with MSU locus IDs in rice. The ones which have been identified in previous studies have been mentioned.**

| Sl. No. | Gene name | TIGR Locus ID | Previous Name | Reference |
|---|---|---|---|---|
| 1 | *ALB1* | LOC_Os03g55730 | *Ory10.3* | Xu and Messing, 2009 |
| 2 | *ALB2* | LOC_Os03g55734 | *Ory10.2* | Xu and Messing, 2009 |
| 3 | *ALB3* | LOC_Os03g55740 | *Ory10.1* | Xu and Messing, 2009 |
| 4 | *ALB4* | LOC_Os05g41970 | | - |
| 5 | *ALB5* | LOC_Os07g11310 | | - |
| 6 | *ALB6* | LOC_Os07g11320 | | - |
| 7 | *ALB7* | LOC_Os07g11330 | | - |
| 8 | *ALB8* | LOC_Os07g11360 | *RAG1* | Wu et al, 1998 |
| 9 | *ALB9* | LOC_Os07g11380 | *RAG2* | Zhou et al, 2017 |
| 10 | *ALB10* | LOC_Os07g11410 | | - |
| 11 | *ALB11* | LOC_Os07g11510 | | - |
| 12 | *ALB12* | LOC_Os07g11630 | | - |
| 13 | *ALB13* | LOC_Os07g11650 | | - |
| 14 | *ALB14* | LOC_Os07g12080 | | - |
| 15 | *ALB15* | LOC_Os07g12090 | | - |
| 16 | *ALB16* | LOC_Os11g33000 | *Ory10.4* | Xu and Messing, 2009 |
| 17 | *GLB1* | LOC_Os03g10110 | | - |
| 18 | *GLB2* | LOC_Os03g21790 | | - |
| 19 | *GLB3* | LOC_Os03g46100 | | - |
| 20 | *GLB4* | LOC_Os03g57960 | | - |
| 21 | *GLU1* | LOC_Os01g55630 | | - |
| 22 | *GLU2* | LOC_Os01g55690 | *GluA-1* | Kawakatsu et al., 2008 |
| 23 | *GLU3* | LOC_Os01g74480 | | - |
| 24 | *GLU4* | LOC_Os02g14600 | *GluB-7* | Kawakatsu et al., 2008 |
| 25 | *GLU5* | LOC_Os02g15070 | *GluB-6* | Kawakatsu et al., 2008 |
| 26 | *GLU6* | LOC_Os02g15090 | *GluD-1* | Kawakatsu et al., 2008 |
| 27 | *GLU7* | LOC_Os02g15150 | *GluB-2* | Kawakatsu et al., 2008 |
| 28 | *GLU8* | LOC_Os02g15169 | *GluB-1b* | Kawakatsu et al., 2008 |
| 29 | *GLU9* | LOC_Os02g15178 | *GluB-1a* | Kawakatsu et al., 2008 |
| 30 | *GLU10* | LOC_Os02g16820 | *GluB-5* | Kawakatsu et al., 2008 |
| 31 | *GLU11* | LOC_Os02g16830 | *GluB-4* | Kawakatsu et al., 2008 |
| 32 | *GLU12* | LOC_Os02g25640 | *GluC-1* | Kawakatsu et al., 2008 |
| 33 | *GLU13* | LOC_Os02g25860 | | - |
| 34 | *GLU14* | LOC_Os03g31360 | *GluA-3* | Kawakatsu et al., 2008 |
| 35 | *GLU15* | LOC_Os05g02520 | | - |
| 36 | *GLU16* | LOC_Os08g03410 | | - |
| 37 | *GLU17* | LOC_Os09g37958 | | - |
| 38 | *GLU18* | LOC_Os09g37967 | | - |
| 39 | *GLU19* | LOC_Os09g37976 | | - |
| 40 | *GLU20* | LOC_Os10g26060 | *GluA-2* | Kawakatsu et al., 2008 |
| 41 | *PRO1* | LOC_Os05g26240 | *Ory13b1* | Xu and Messing, 2009 |
| 42 | *PRO2* | LOC_Os05g26400 | *Ory13b3/ Ory13b2/ Ory13b2* | Xu and Messing, 2009 |
| 43 | *PRO3* | LOC_Os05g26350 | *Ory13b4* | Xu and Messing, 2009 |
| 44 | *PRO4* | LOC_Os05g26368 | *Ory13b5* | Xu and Messing, 2009 |
| 45 | *PRO5* | LOC_Os05g26377 | *Ory13b7/ Ory13b6* | Xu and Messing, 2009 |
| 46 | *PRO6* | LOC_Os05g26440 | *Ory13b10* | Xu and Messing, 2009 |
| 47 | *PRO7* | LOC_Os05g26460 | *Ory13b11* | Xu and Messing, 2009 |
| 48 | *PRO8* | LOC_Os05g26480 | *Ory13b12* | Xu and Messing, 2009 |
| 49 | *PRO9* | LOC_Os05g26490 | *Ory13b13* | Xu and Messing, 2009 |
| 50 | *PRO10* | LOC_Os05g26620 | *Ory13b14* | Xu and Messing, 2009 |
| 51 | *PRO11* | LOC_Os05g26690 | *Ory13b15* | Xu and Messing, 2009 |
| 52 | *PRO12* | LOC_Os05g26720 | *Ory13b16* | Xu and Messing, 2009 |
| 53 | *PRO13* | LOC_Os05g26750 | *Ory13b17* | Xu and Messing, 2009 |
| 54 | *PRO14* | LOC_Os05g26770 | *Ory13b18* | Xu and Messing, 2009 |
| 55 | *PRO15* | LOC_Os06g31060 | *Ory13a1* | Xu and Messing, 2009 |
| 56 | *PRO16* | LOC_Os06g31070 | *Ory13a2* | Xu and Messing, 2009 |
| 57 | *PRO17* | LOC_Os07g10570 | *Ory16.1* | Xu and Messing, 2009 |
| 58 | *PRO18* | LOC_Os07g10580 | *Ory16.2* | Xu and Messing, 2009 |
| 59 | *PRO19* | LOC_Os07g11900 | *Ory13b19* | Xu and Messing, 2009 |
| 60 | *PRO20* | LOC_Os07g11910 | *Ory13b20* | Xu and Messing, 2009 |
| 61 | *PRO21* | LOC_Os07g11920 | *Ory13b21/ Ory13b22* | Xu and Messing, 2009 |
| 62 | *PRO22* | LOC_Os12g16880 | *Ory16.3* | Xu and Messing, 2009 |
| 63 | *PRO23* | LOC_Os12g16890 | *Ory16.4* | Xu and Messing, 2009 |
| 64 | *PRO24* | LOC_Os12g17010 | *Ory16.5* | Xu and Messing, 2009 |
| 65 | *PRO25* | LOC_Os12g17030 | *Ory16.6* | Xu and Messing, 2009 |

**Table S3. Tandemly duplicated rice *SSP*.**

| Group No. | Duplicated genes | Chromosome Number | Number of genes between the duplicated genes |
|---|---|---|---|
| I | *ALB1-ALB2* | 3 | 0 |
| I | *ALB2-ALB3* | 3 | 0 |
| II | *ALB5-ALB6* | 3 | 0 |
| II | *ALB6-ALB7* | 7 | 0 |
| II | *ALB7-ALB8* | 7 | 2 |
| II | *ALB8-ALB9* | 7 | 1 |
| II | *ALB9-ALB10* | 7 | 2 |
| III | *ALB12-ALB13* | 7 | 1 |
| IV | *ALB14-ALB15* | 7 | 0 |
| I | *GLU1-GLU2* | 1 | 4 |
| II | *GLU5-GLU6* | 2 | 1 |
| II | *GLU6-GLU7* | 2 | 5 |
| II | *GLU7-GLU8* | 2 | 2 |
| II | *GLU8-GLU9* | 2 | 0 |
| III | *GLU10-GLU11* | 2 | 0 |
| IV | *GLU17-GLU18* | 9 | 0 |
| IV | *GLU18-GLU19* | 9 | 0 |
| I | *PRO1-PRO2* | 5 | 0 |
| II | *PRO3-PRO4* | 5 | 0 |
| II | *PRO4-PRO5* | 5 | 0 |
| II | *PRO5-PRO6* | 5 | 3 |
| II | *PRO6-PRO7* | 5 | 1 |
| II | *PRO7-PRO8* | 5 | 1 |
| II | *PRO8-PRO9* | 5 | 0 |
| III | *PRO10-PRO11* | 5 | 5 |
| III | *PRO11-PRO12* | 5 | 2 |
| III | *PRO12-PRO13* | 5 | 2 |
| III | *PRO13-PRO14* | 5 | 1 |
| IV | *PRO15-PRO16* | 6 | 0 |
| V | *PRO17-PRO18* | 7 | 0 |
| VI | *PRO19-PRO20* | 7 | 0 |
| VI | *PRO20-PRO21* | 7 | 0 |
| VII | *PRO22-PRO23* | 12 | 0 |
| VIII | *PRO24-PRO25* | 12 | 1 |

**Table S4. Sequences of seed-specific cis-elements in rice SSP promoter identified from PLACE database.**

| Seed Specific Element | Sequence |
|---|---|
| 2SSEEDPROTBANAPA | CAAACAC |
| -300CORE | TGTAAAG |
| -300ELEMENT | TGHAAARK |
| -300MOTIFZMZEIN | RTGAGTCAT |
| AACACOREOSGLUB1 | AACAAAC |
| AACAOSGLUB1 | CAACAAACTATATC |
| ACGTABOX | TACGTA |
| ACGTOSGLUB1 | GTACGTG |
| ACGTSEED3 | GTACGTGGCG |
| AMYBOX1 | TAACARA |
| AMYBOX2 | TATCCAT |
| ARE1 | RGTGACNNNGC |
| CAATBOX1 | CAAT |
| CANBNNAPA | CNAACAC |
| CAREOSREP1 | CAACTC |
| CEREGLUBOX1PSLEGA | TGTTAAAGT |
| CEREGLUBOX2PSLEGA | TGAAAACT |
| CEREGLUBOX3PSLEGA | TGTAAAAGT |
| DOFCOREZM | AAAG |
| DPBFCOREDCDC3 | ACACNNG |
| DRE2COREZMRAB17 | ACCGAC |
| EBOXBNNAPA | CANNTG |
| EMHVCHORD | TGTAAAGT |
| GADOWNAT | ACGTGTC |
| GARE1OSREP1 | TAACAGA |
| GARE2OSREP1 | TAACGTA |
| GAREAT | TAACAAR |
| GCN4OSGLUB1 | TGAGTCA |
| GLMHVCHORD | RTGASTCAT |
| GLUTAACAOS | AACAAACTCTAT |
| GLUTEBOX1OSGT2 | ATATCATGAGTCACTTCA |
| GLUTEBOX1OSGT3 | TATCTAGTGAGTCACTTCA |
| GLUTEBOX2OSGT2 | TCCGTGTACCA |
| GLUTEBOX2OSGT3 | CTTTTGTGTACCTTA |
| GLUTEBP1OS | AAGCAACACACAAC |
| GLUTEBP2OS | ATGCTCAATAGATATAAGT |
| GLUTECOREOS | CTTTCGTGTAC |
| MYBGAHV | TAACAAA |
| NAPINMOTIFBN | TACACAT |
| NTBBF1ARROLB | ACTTTA |
| OPAQUE2ZMB32 | GATGAYRTGG |
| POLASIG1 | AATAAA |
| POLASIG2 | AATTAAA |
| PROLAMINBOXOSGLUB1 | TGCAAAG |
| PROXBBNNAPA | CAAACACC |
| PYRIMIDINEBOXHVEPB1 | TTTTTTCC |
| PYRIMIDINEBOXOSRAMY1A | CCTTTT |
| RYREPEATBNNAPA | CATGCA |
| RYREPEATGMGY2 | CATGCAT |
| RYREPEATLEGUMINBOX | CATGCAY |
| RYREPEATVFLEB4 | CATGCATG |
| SEF1MOTIF | ATATTTAWW |
| SEF3MOTIFGM | AACCCA |
| SEF4MOTIFGM7S | RTTTTTR |
| SP8BFIBSP8BIB | TACTATT |
| SPHCOREZMC1 | TCCATGCAT |
| SURE1STPAT21 | AATAGAAAA |
| TATABOX1 | CTATAAATAC |
| TATABOX2 | TATAAAT |
| TATABOX3 | TATTAT |
| TATABOX4 | TATATAA |
| TATABOX5 | TTATTT |
| TATCCAOSAMY | TATCCA |
| TATCCAYMOTIFOSRAMY3D | TATCCAY |
| TGACGTVMAMY | TGACGT |
| WBOXHVISO1 | TGACT |
| WRKY71OS | TGAC |

**Table S5. *Cis*-elements of SSP promoters with all the sequences.**

| Element | Sequence |
|---|---|
| 2SSEEDPROTBANAPA | CAAACAC |
| -300ELEMENT | GAAAAAG |
| -300ELEMENT | GAAAAAT |
| -300ELEMENT | GAAAAGG |
| -300ELEMENT | GAAAAGT |
| -300ELEMENT | GCAAAAG |
| -300ELEMENT | GCAAAAT |
| -300ELEMENT | GCAAAGG |
| -300ELEMENT | GCAAAGT |
| -300ELEMENT | GTAAAAG |
| -300ELEMENT | GTAAAAT |
| -300ELEMENT | GTAAAGG |
| -300ELEMENT | GTAAAGT |
| -300MOTIFZMZEIN | ATGAGTCAT |
| -300MOTIFZMZEIN | GTGAGTCAT |
| AACACOREOSGLUB1 | AACAAAC |
| ACGTABOX | TACGTA |
| ACGTOSGLUB1 | GTACGTG |
| CAATBOX1 | CAAT |
| CANBNNAPA | CAAACAC |
| CANBNNAPA | CTAACAC |
| CANBNNAPA | CCAACAC |
| CANBNNAPA | CGAACAC |
| CEREGLUBOX3PSLEGA | TGTAAAAGT |
| DOFCOREZM | AAAG |
| DPBFCOREDCDC3 | ACACAAAG |
| DPBFCOREDCDC3 | ACACAATG |
| DPBFCOREDCDC3 | ACACAACG |
| DPBFCOREDCDC3 | ACACAAGG |
| DPBFCOREDCDC3 | ACACTAAG |
| DPBFCOREDCDC3 | ACACTATG |
| DPBFCOREDCDC3 | ACACTACG |
| DPBFCOREDCDC3 | ACACTAGG |
| DPBFCOREDCDC3 | ACACCAAG |
| DPBFCOREDCDC3 | ACACCATG |
| DPBFCOREDCDC3 | ACACCACG |
| DPBFCOREDCDC3 | ACACCAGG |
| DPBFCOREDCDC3 | ACACGAAG |
| DPBFCOREDCDC3 | ACACGATG |
| DPBFCOREDCDC3 | ACACGACG |
| DPBFCOREDCDC3 | ACACGAGG |
| EBOXBNNAPA | CAAATG |
| EBOXBNNAPA | CAACTG |
| EBOXBNNAPA | CAAGTG |
| EBOXBNNAPA | CAATTG |
| EBOXBNNAPA | CATAATG |
| EBOXBNNAPA | CATACTG |
| EBOXBNNAPA | CATAGTG |
| EBOXBNNAPA | CATATTG |
| EBOXBNNAPA | CACAATG |
| EBOXBNNAPA | CACACTG |
| EBOXBNNAPA | CACAGTG |
| EBOXBNNAPA | CACATTG |
| EBOXBNNAPA | CAGAATG |
| EBOXBNNAPA | CAGACTG |
| EBOXBNNAPA | CAGAGTG |
| EBOXBNNAPA | CAGATTG |
| GCN4OSGLUB1 | TGAGTCA |
| GLUTAACAOS | AACAAACTCTAT |
| GLUTEBOX1OSGT2 | ATATCATGAGTCACTTCA |
| GLUTEBOX1OSGT3 | TATCTAGTGAGTCACTTCA |
| GLUTEBOX2OSGT2 | TCCGTGTACCA |
| GLUTEBOX2OSGT3 | CTTTTGTGTACCTTA |
| GLUTEBP1OS | AAGCAACACACAAC |
| GLUTEBP2OS | ATGCTCAATAGATATAAGT |
| GLUTECOREOS | CTTTCGTGTAC |
| NAPINMOTIFBN | TACACAT |
| PROLAMINBOXOSGLUB1 | TGCAAAG |
| RYREPEATBNNAPA | CATGCA |
| SEF4MOTIFGM7S | ATTTTTA |
| SEF4MOTIFGM7S | ATTTTTG |
| SEF4MOTIFGM7S | GTTTTTA |
| SEF4MOTIFGM7S | GTTTTTG |
| SP8BFIBSP8BIB | TACTATT |
| TATABOX1 | CTATAAATAC |
| TATABOX2 | TATAAAT |
| TATABOX3 | TATTAT |
| TATABOX4 | TATATAA |
| TATABOX5 | TTATTT |
| WRKY71OS | TGAC |
| DRE2COREZMRAB17 | ACCGAC |
| POLASIG1 | AATAAA |
| NTBBF1ARROLB | ACTTTA |
| WBOXHVISO1 | TGACT |
| CAREOSREP1 | CAACTC |

| | |
|---|---|
| RYREPEATLEGUMINBOX | CATGCAC |
| RYREPEATLEGUMINBOX | CATGCAT |
| RYREPEATGMGY2 | CATGCAT |
| SURE1STPAT21 | AATAGAAAA |
| TATCCAOSAMY | TATCCA |
| | TATCCAC |
| TATCCAYMOTIFOSRAMY3D | TATCCAT |
| PYRIMIDINEBOXOSRAMY1A | CCTTTT |
| POLASIG2 | AATTAAA |
| GADOWNAT | ACGTGTC |
| SEF3MOTIFGM | AACCCA |
| AMYBOX1 | TAACAAA |
| AMYBOX1 | TAACAGG |
| MYBGAHV | TAACAAA |
| GAREAT | TAACAAA |
| GAREAT | TAACAAG |
| PYRIMIDINEBOXHVEPB1 | TTTTTTCC |
| GARE2OSREP1 | TAACGTA |
| PROXBBNNAPA | CAAACACC |
| GLMHVCHORD | RTGASTCAT |
| GLMHVCHORD | ATGACTCST |
| GLMHVCHORD | ATGAGTCST |
| GLMHVCHORD | GTGACTCST |
| GLMHVCHORD | GTGAGTCST |
| TGACGTVMAMY | TGACGT |
| AMYBOX2 | TATCCAT |
| GARE1OSREP1 | TAACAGA |
| SEF1MOTIF | ATATTTAAA |
| SEF1MOTIF | ATATTTAAT |
| SEF1MOTIF | ATATTTATA |
| SEF1MOTIF | ATATTTATT |
| -300CORE | TGTAAAG |
| CEREGLUBOX1PSLEGA | TGTTAAAGT |
| RYREPEATVFLEB4 | CATGCATG |
| EMHVCHORD | TGTAAAGT |
| SPHCOREZMC1 | TCCATGCAT |
| AACAOSGLUB1 | CAACAAACTATATC |
| OPAQUE2ZMB32 | GATGACGTGG |
| OPAQUE2ZMB32 | GATGACATGG |
| OPAQUE2ZMB32 | GATGATGTGG |
| OPAQUE2ZMB32 | GATGATATGG |
| CEREGLUBOX2PSLEGA | TGAAAACT |
| ACGTSEED3 | GTACGTGGCG |
| ARE1 | AGTGACAAAGC |
| ARE1 | AGTGACAATGC |
| ARE1 | AGTGACAAGGC |
| ARE1 | AGTGACAACGC |
| ARE1 | AGTGACATAGC |
| ARE1 | AGTGACATTGC |
| ARE1 | AGTGACATGGC |
| ARE1 | AGTGACATCGC |
| ARE1 | AGTGACAGAGC |
| ARE1 | AGTGACAGTGC |
| ARE1 | AGTGACAGGGC |
| ARE1 | AGTGACAGCGC |
| ARE1 | AGTGACACAGC |
| ARE1 | AGTGACACTGC |
| ARE1 | AGTGACACGGC |
| ARE1 | AGTGACACCGC |
| ARE1 | AGTGACTAAGC |
| ARE1 | AGTGACTATGC |
| ARE1 | AGTGACTAGGC |
| ARE1 | AGTGACTACGC |
| ARE1 | AGTGACTTAGC |
| ARE1 | AGTGACTTTGC |
| ARE1 | AGTGACTTGGC |
| ARE1 | AGTGACTTCGC |
| ARE1 | AGTGACTGAGC |
| ARE1 | AGTGACTGTGC |
| ARE1 | AGTGACTGGGC |
| ARE1 | AGTGACTGCGC |
| ARE1 | AGTGACTCAGC |
| ARE1 | AGTGACTCTGC |
| ARE1 | AGTGACTCGGC |
| ARE1 | AGTGACTCCGC |
| ARE1 | AGTGACGAAGC |
| ARE1 | AGTGACGATGC |
| ARE1 | AGTGACGAGGC |
| ARE1 | AGTGACGACGC |
| ARE1 | AGTGACGTAGC |
| ARE1 | AGTGACGTTGC |
| ARE1 | AGTGACGTGGC |
| ARE1 | AGTGACGTCGC |
| ARE1 | AGTGACGGAGC |
| ARE1 | AGTGACGGTGC |

| | |
|---|---|
| ARE1 | AGTGACGGGGC |
| ARE1 | AGTGACGGCGC |
| ARE1 | AGTGACGCAGC |
| ARE1 | AGTGACGCTGC |
| ARE1 | AGTGACGCGGC |
| ARE1 | AGTGACGCCGC |
| ARE1 | AGTGACCAAGC |
| ARE1 | AGTGACCATGC |
| ARE1 | AGTGACCAGGC |
| ARE1 | AGTGACCACGC |
| ARE1 | AGTGACCTAGC |
| ARE1 | AGTGACCTTGC |
| ARE1 | AGTGACCTGGC |
| ARE1 | AGTGACCTCGC |
| ARE1 | AGTGACCGAGC |
| ARE1 | AGTGACCGTGC |
| ARE1 | AGTGACCGGGC |
| ARE1 | AGTGACCGCGC |
| ARE1 | AGTGACCCAGC |
| ARE1 | AGTGACCCTGC |
| ARE1 | AGTGACCCGGC |
| ARE1 | AGTGACCCCGC |
| ARE1 | GGTGACAAAGC |
| ARE1 | GGTGACAATGC |
| ARE1 | GGTGACAAGGC |
| ARE1 | GGTGACAACGC |
| ARE1 | GGTGACATAGC |
| ARE1 | GGTGACATTGC |
| ARE1 | GGTGACATGGC |
| ARE1 | GGTGACATCGC |
| ARE1 | GGTGACAGAGC |
| ARE1 | GGTGACAGTGC |
| ARE1 | GGTGACAGGGC |
| ARE1 | GGTGACAGCGC |
| ARE1 | GGTGACACAGC |
| ARE1 | GGTGACACTGC |
| ARE1 | GGTGACACGGC |
| ARE1 | GGTGACACCGC |
| ARE1 | GGTGACTAAGC |
| ARE1 | GGTGACTATGC |
| ARE1 | GGTGACTAGGC |
| ARE1 | GGTGACTACGC |
| ARE1 | GGTGACTTAGC |
| ARE1 | GGTGACTTTGC |
| ARE1 | GGTGACTTGGC |
| ARE1 | GGTGACTTCGC |
| ARE1 | GGTGACTGAGC |
| ARE1 | GGTGACTGTGC |
| ARE1 | GGTGACTGGGC |
| ARE1 | GGTGACTGCGC |
| ARE1 | GGTGACTCAGC |
| ARE1 | GGTGACTCTGC |
| ARE1 | GGTGACTCGGC |
| ARE1 | GGTGACTCCGC |
| ARE1 | GGTGACGAAGC |
| ARE1 | GGTGACGATGC |
| ARE1 | GGTGACGAGGC |
| ARE1 | GGTGACGACGC |
| ARE1 | GGTGACGTAGC |
| ARE1 | GGTGACGTTGC |
| ARE1 | GGTGACGTGGC |
| ARE1 | GGTGACGTCGC |
| ARE1 | GGTGACGGAGC |
| ARE1 | GGTGACGGTGC |
| ARE1 | GGTGACGGGGC |
| ARE1 | GGTGACGGCGC |
| ARE1 | GGTGACGCAGC |
| ARE1 | GGTGACGCTGC |
| ARE1 | GGTGACGCGGC |
| ARE1 | GGTGACGCCGC |
| ARE1 | GGTGACCAAGC |
| ARE1 | GGTGACCATGC |
| ARE1 | GGTGACCAGGC |
| ARE1 | GGTGACCACGC |
| ARE1 | GGTGACCTAGC |
| ARE1 | GGTGACCTTGC |
| ARE1 | GGTGACCTGGC |
| ARE1 | GGTGACCTCGC |
| ARE1 | GGTGACCGAGC |
| ARE1 | GGTGACCGTGC |
| ARE1 | GGTGACCGGGC |
| ARE1 | GGTGACCGCGC |
| ARE1 | GGTGACCCAGC |
| ARE1 | GGTGACCCTGC |
| ARE1 | GGTGACCCGGC |
| ARE1 | GGTGACCCCGC |

**Table S6. Total numbers of each seed-specific *cis* -element identified in each *SSP* promoter sequence.**

| Gene | 2SSEEDPROTBANAPA | -300ELEMENT | -300MOTIFZMZEIN | AACACOREOSGLUB1 | ACGTABOX | ACGTOSGLUB1 | CAATBOX1 | CANBNNAPA | CEREGLUBOX3PSLEGA | DOFCOREZM | DPBFCOREDCDC3 | EBOXBNNAPA | GCN4OSGLUB1 | GLUTAACAOS | GLUTEBOX1OSGT2 | GLUTEBOX1OSGT3 | GLUTEBOX2OSGT2 | GLUTEBOX2OSGT3 | GLUTEBP1OS | GLUTEBP2OS |
|---|---|---|---|---|---|---|---|---|---|---|---|---|---|---|---|---|---|---|---|---|
| *ALB1* | 0 | 0 | 0 | 0 | 0 | 0 | 0 | 0 | 0 | 1 | 0 | 2 | 0 | 0 | 0 | 0 | 0 | 0 | 0 | 0 |
| *ALB2* | 0 | 0 | 0 | 1 | 0 | 0 | 30 | 0 | 0 | 5 | 5 | 30 | 0 | 0 | 0 | 0 | 0 | 0 | 0 | 0 |
| *ALB3* | 1 | 2 | 1 | 0 | 4 | 1 | 31 | 2 | 1 | 29 | 2 | 14 | 1 | 0 | 0 | 0 | 0 | 0 | 0 | 0 |
| *ALB4* | 0 | 1 | 0 | 1 | 0 | 0 | 10 | 1 | 0 | 15 | 0 | 10 | 0 | 0 | 0 | 0 | 0 | 0 | 0 | 0 |
| *ALB5* | 1 | 2 | 0 | 0 | 4 | 0 | 23 | 1 | 1 | 27 | 0 | 6 | 0 | 0 | 0 | 0 | 0 | 0 | 0 | 0 |
| *ALB6* | 1 | 1 | 0 | 0 | 2 | 1 | 26 | 2 | 0 | 31 | 3 | 16 | 0 | 0 | 0 | 0 | 0 | 0 | 0 | 0 |
| *ALB7* | 1 | 1 | 0 | 0 | 0 | 1 | 21 | 1 | 0 | 25 | 1 | 12 | 0 | 0 | 0 | 0 | 0 | 0 | 0 | 0 |
| *ALB8* | 0 | 1 | 0 | 0 | 0 | 0 | 15 | 0 | 0 | 24 | 1 | 8 | 0 | 0 | 0 | 0 | 0 | 0 | 0 | 0 |
| *ALB9* | 0 | 1 | 0 | 0 | 0 | 0 | 15 | 0 | 0 | 24 | 1 | 8 | 0 | 0 | 0 | 0 | 0 | 0 | 0 | 0 |
| *ALB10* | 0 | 3 | 0 | 1 | 2 | 1 | 22 | 0 | 0 | 32 | 0 | 6 | 0 | 0 | 0 | 0 | 0 | 0 | 0 | 0 |
| *ALB11* | 2 | 0 | 0 | 1 | 2 | 0 | 24 | 2 | 0 | 37 | 2 | 8 | 1 | 0 | 0 | 0 | 0 | 0 | 0 | 0 |
| *ALB12* | 0 | 3 | 0 | 0 | 2 | 1 | 32 | 1 | 0 | 26 | 0 | 14 | 0 | 0 | 0 | 0 | 0 | 0 | 0 | 0 |
| *ALB13* | 2 | 0 | 0 | 1 | 0 | 0 | 27 | 4 | 0 | 31 | 2 | 20 | 0 | 0 | 0 | 0 | 0 | 0 | 0 | 0 |
| *ALB14* | 0 | 1 | 0 | 0 | 0 | 0 | 5 | 0 | 0 | 8 | 0 | 4 | 0 | 0 | 0 | 0 | 0 | 0 | 0 | 0 |
| *ALB15* | 1 | 2 | 0 | 2 | 2 | 0 | 13 | 2 | 0 | 15 | 4 | 14 | 0 | 0 | 0 | 0 | 0 | 0 | 0 | 0 |
| *ALB16* | 0 | 3 | 0 | 1 | 0 | 0 | 33 | 0 | 0 | 25 | 5 | 24 | 1 | 0 | 0 | 0 | 0 | 0 | 0 | 0 |
| *GLB1* | 0 | 1 | 0 | 0 | 0 | 0 | 8 | 0 | 0 | 32 | 2 | 6 | 0 | 0 | 0 | 0 | 0 | 0 | 0 | 0 |
| *GLB2* | 0 | 1 | 0 | 0 | 2 | 0 | 19 | 0 | 0 | 14 | 6 | 20 | 0 | 0 | 0 | 0 | 0 | 0 | 0 | 0 |
| *GLB3* | 0 | 1 | 0 | 0 | 0 | 1 | 8 | 0 | 0 | 7 | 2 | 6 | 0 | 0 | 0 | 0 | 0 | 0 | 0 | 0 |
| *GLB4* | 0 | 0 | 0 | 1 | 2 | 1 | 13 | 4 | 0 | 19 | 2 | 14 | 0 | 0 | 0 | 0 | 0 | 0 | 0 | 0 |
| *GLU1* | 1 | 2 | 1 | 2 | 0 | 0 | 18 | 1 | 0 | 23 | 2 | 18 | 1 | 0 | 0 | 0 | 0 | 0 | 0 | 0 |
| *GLU2* | 0 | 4 | 0 | 2 | 0 | 0 | 23 | 2 | 0 | 37 | 3 | 22 | 2 | 1 | 1 | 0 | 1 | 0 | 0 | 0 |
| *GLU3* | 1 | 0 | 0 | 2 | 0 | 0 | 21 | 1 | 0 | 26 | 0 | 12 | 0 | 0 | 0 | 0 | 0 | 0 | 0 | 0 |
| *GLU4* | 0 | 2 | 0 | 1 | 4 | 0 | 14 | 0 | 0 | 34 | 2 | 16 | 0 | 0 | 0 | 0 | 0 | 0 | 0 | 0 |
| *GLU5* | 2 | 5 | 0 | 3 | 0 | 0 | 30 | 3 | 0 | 27 | 1 | 16 | 1 | 0 | 0 | 0 | 0 | 0 | 0 | 0 |
| *GLU6* | 0 | 2 | 0 | 1 | 2 | 0 | 34 | 1 | 0 | 30 | 4 | 16 | 0 | 0 | 0 | 0 | 0 | 0 | 0 | 0 |
| *GLU7* | 0 | 3 | 0 | 4 | 0 | 1 | 22 | 0 | 0 | 29 | 2 | 20 | 1 | 0 | 0 | 0 | 0 | 0 | 0 | 0 |
| *GLU8* | 0 | 3 | 0 | 4 | 0 | 1 | 22 | 0 | 0 | 29 | 2 | 20 | 1 | 0 | 0 | 0 | 0 | 0 | 0 | 0 |
| *GLU9* | 0 | 3 | 0 | 4 | 0 | 1 | 22 | 0 | 0 | 29 | 2 | 20 | 1 | 0 | 0 | 0 | 0 | 0 | 0 | 0 |
| *GLU10* | 0 | 3 | 1 | 3 | 0 | 0 | 12 | 0 | 0 | 20 | 1 | 8 | 1 | 1 | 0 | 0 | 0 | 0 | 0 | 0 |
| *GLU11* | 0 | 2 | 1 | 3 | 0 | 0 | 11 | 0 | 0 | 18 | 1 | 6 | 1 | 1 | 0 | 0 | 0 | 0 | 0 | 0 |
| *GLU12* | 0 | 1 | 0 | 2 | 0 | 0 | 19 | 1 | 0 | 34 | 1 | 8 | 0 | 0 | 0 | 0 | 0 | 0 | 0 | 0 |
| *GLU13* | 0 | 1 | 0 | 2 | 0 | 1 | 17 | 0 | 0 | 33 | 1 | 10 | 1 | 1 | 0 | 1 | 0 | 1 | 1 | 1 |
| *GLU14* | 0 | 1 | 0 | 0 | 0 | 0 | 2 | 2 | 0 | 12 | 2 | 4 | 0 | 0 | 0 | 0 | 0 | 0 | 0 | 0 |
| *GLU15* | 0 | 6 | 0 | 0 | 2 | 0 | 19 | 1 | 0 | 18 | 1 | 14 | 1 | 0 | 0 | 0 | 0 | 0 | 0 | 0 |
| *GLU16* | 0 | 3 | 1 | 2 | 0 | 0 | 25 | 2 | 0 | 31 | 2 | 16 | 2 | 1 | 1 | 0 | 0 | 0 | 0 | 0 |
| *GLU17* | 0 | 1 | 0 | 1 | 0 | 0 | 13 | 0 | 0 | 15 | 5 | 26 | 0 | 0 | 0 | 0 | 0 | 0 | 0 | 0 |
| *GLU18* | 1 | 0 | 0 | 0 | 2 | 0 | 21 | 1 | 0 | 24 | 2 | 14 | 0 | 0 | 0 | 0 | 0 | 0 | 0 | 0 |
| *GLU19* | 0 | 1 | 0 | 1 | 0 | 0 | 13 | 0 | 0 | 15 | 5 | 26 | 0 | 0 | 0 | 0 | 0 | 0 | 0 | 0 |
| *GLU20* | 0 | 2 | 0 | 2 | 0 | 0 | 21 | 0 | 0 | 26 | 1 | 14 | 0 | 0 | 0 | 0 | 0 | 0 | 0 | 0 |
| *PRO1* | 0 | 0 | 1 | 0 | 0 | 0 | 21 | 2 | 0 | 23 | 2 | 16 | 1 | 0 | 0 | 0 | 0 | 0 | 0 | 0 |
| *PRO2* | 0 | 1 | 0 | 0 | 0 | 0 | 16 | 0 | 0 | 18 | 2 | 10 | 0 | 0 | 0 | 0 | 0 | 0 | 0 | 0 |
| *PRO3* | 2 | 0 | 2 | 2 | 0 | 0 | 25 | 3 | 0 | 24 | 1 | 18 | 2 | 0 | 0 | 0 | 0 | 0 | 0 | 0 |
| *PRO4* | 2 | 2 | 2 | 1 | 2 | 0 | 17 | 3 | 0 | 27 | 1 | 20 | 2 | 0 | 0 | 0 | 0 | 0 | 0 | 0 |
| *PRO5* | 1 | 0 | 2 | 2 | 0 | 0 | 20 | 2 | 0 | 27 | 1 | 12 | 2 | 0 | 0 | 0 | 0 | 0 | 0 | 0 |
| *PRO6* | 0 | 1 | 1 | 0 | 0 | 0 | 8 | 1 | 0 | 22 | 2 | 10 | 1 | 0 | 0 | 0 | 0 | 0 | 0 | 0 |
| *PRO7* | 2 | 2 | 2 | 1 | 2 | 0 | 19 | 3 | 0 | 27 | 1 | 20 | 2 | 0 | 0 | 0 | 0 | 0 | 0 | 0 |
| *PRO8* | 0 | 1 | 2 | 0 | 0 | 0 | 18 | 1 | 0 | 18 | 3 | 16 | 2 | 0 | 0 | 0 | 0 | 0 | 0 | 0 |
| *PRO9* | 2 | 1 | 1 | 0 | 2 | 0 | 9 | 3 | 0 | 24 | 1 | 16 | 1 | 0 | 0 | 0 | 0 | 0 | 0 | 0 |
| *PRO10* | 2 | 1 | 1 | 0 | 2 | 0 | 13 | 3 | 0 | 24 | 1 | 20 | 1 | 0 | 0 | 0 | 0 | 0 | 0 | 0 |
| *PRO11* | 1 | 2 | 1 | 0 | 0 | 0 | 13 | 2 | 0 | 28 | 2 | 18 | 1 | 0 | 0 | 0 | 0 | 0 | 0 | 0 |
| *PRO12* | 0 | 0 | 2 | 0 | 0 | 0 | 8 | 1 | 0 | 11 | 2 | 6 | 2 | 0 | 0 | 0 | 0 | 0 | 0 | 0 |
| *PRO13* | 1 | 1 | 2 | 0 | 0 | 1 | 14 | 3 | 0 | 24 | 3 | 20 | 2 | 0 | 0 | 0 | 0 | 0 | 0 | 0 |
| *PRO14* | 1 | 0 | 0 | 0 | 0 | 0 | 0 | 0 | 0 | 0 | 0 | 0 | 0 | 0 | 0 | 0 | 0 | 0 | 0 | 0 |
| *PRO15* | 0 | 0 | 0 | 0 | 0 | 1 | 18 | 1 | 0 | 17 | 1 | 22 | 1 | 0 | 0 | 0 | 0 | 0 | 0 | 0 |
| *PRO16* | 0 | 3 | 2 | 1 | 2 | 2 | 23 | 0 | 0 | 19 | 1 | 26 | 2 | 0 | 0 | 0 | 0 | 0 | 0 | 0 |
| *PRO17* | 0 | 4 | 2 | 0 | 0 | 0 | 17 | 1 | 0 | 31 | 1 | 18 | 2 | 0 | 0 | 0 | 0 | 0 | 0 | 0 |
| *PRO18* | 0 | 1 | 2 | 1 | 0 | 0 | 24 | 1 | 0 | 29 | 2 | 18 | 2 | 0 | 0 | 0 | 0 | 0 | 0 | 0 |
| *PRO19* | 2 | 0 | 2 | 2 | 0 | 0 | 25 | 3 | 0 | 21 | 1 | 22 | 2 | 0 | 0 | 0 | 0 | 0 | 0 | 0 |
| *PRO20* | 2 | 0 | 2 | 2 | 0 | 0 | 26 | 3 | 0 | 24 | 1 | 22 | 2 | 0 | 0 | 0 | 0 | 0 | 0 | 0 |
| *PRO21* | 2 | 1 | 2 | 3 | 4 | 2 | 20 | 3 | 0 | 18 | 3 | 26 | 2 | 0 | 0 | 0 | 0 | 0 | 0 | 0 |
| *PRO22* | 1 | 1 | 2 | 1 | 0 | 0 | 21 | 1 | 0 | 18 | 0 | 22 | 2 | 0 | 0 | 0 | 0 | 0 | 0 | 0 |
| *PRO23* | 1 | 4 | 1 | 2 | 0 | 0 | 16 | 1 | 0 | 22 | 0 | 12 | 1 | 0 | 0 | 0 | 0 | 0 | 0 | 0 |
| *PRO24* | 0 | 1 | 0 | 0 | 0 | 0 | 13 | 0 | 0 | 16 | 2 | 10 | 0 | 0 | 0 | 0 | 0 | 0 | 0 | 0 |
| *PRO25* | 1 | 3 | 1 | 2 | 0 | 0 | 23 | 1 | 0 | 25 | 1 | 14 | 1 | 0 | 0 | 0 | 0 | 0 | 0 | 0 |
| Total occu | 37 | 99 | 40 | 68 | 46 | 17 | 1181 | 77 | 2 | 1474 | 113 | 962 | 52 | 5 | 2 | 1 | 1 | 1 | 1 | 1 |

| Gene | GLUTECOREOS | NAPINMOTIFBN | PROLAMINBOXOSGLUB1 | RYREPEATBNNAPA | SEF4MOTIFGM7S | SP8BFIBSP8BIB | TATABOX1 | TATABOX2 | TATABOX3 | TATABOX4 | TATABOX5 | WRKY71OS | DRE2COREZMRAB17 | POLASIG1 | NTBBF1ARROLB | WBOXHVISO1 | CAREOSREP1 | RYREPEATLEGUMINBOX |
|---|---|---|---|---|---|---|---|---|---|---|---|---|---|---|---|---|---|---|
| *ALB1* | 0 | 0 | 0 | 0 | 0 | 0 | 0 | 0 | 0 | 0 | 0 | 1 | 1 | 0 | 0 | 0 | 0 | 0 |
| *ALB2* | 0 | 2 | 0 | 4 | 3 | 0 | 0 | 0 | 0 | 0 | 2 | 9 | 2 | 2 | 2 | 2 | 2 | 1 |
| *ALB3* | 0 | 0 | 1 | 0 | 6 | 0 | 0 | 5 | 0 | 5 | 11 | 7 | 0 | 9 | 1 | 4 | 0 | 0 |
| *ALB4* | 0 | 0 | 1 | 0 | 2 | 0 | 0 | 0 | 0 | 0 | 2 | 12 | 0 | 2 | 0 | 2 | 1 | 0 |
| *ALB5* | 0 | 0 | 0 | 4 | 5 | 1 | 0 | 1 | 0 | 2 | 2 | 7 | 1 | 6 | 0 | 1 | 1 | 2 |
| *ALB6* | 0 | 0 | 0 | 2 | 1 | 1 | 0 | 1 | 0 | 2 | 6 | 9 | 0 | 5 | 3 | 1 | 0 | 1 |
| *ALB7* | 0 | 0 | 0 | 0 | 1 | 0 | 0 | 1 | 0 | 2 | 5 | 2 | 0 | 4 | 2 | 0 | 0 | 0 |
| *ALB8* | 0 | 0 | 0 | 2 | 5 | 0 | 0 | 0 | 2 | 2 | 6 | 8 | 1 | 8 | 5 | 1 | 1 | 0 |
| *ALB9* | 0 | 0 | 0 | 2 | 5 | 0 | 0 | 0 | 2 | 2 | 6 | 8 | 1 | 8 | 5 | 1 | 1 | 0 |
| *ALB10* | 0 | 0 | 0 | 4 | 8 | 0 | 0 | 0 | 2 | 2 | 12 | 11 | 0 | 8 | 1 | 1 | 2 | 1 |
| *ALB11* | 0 | 0 | 0 | 1 | 3 | 1 | 0 | 5 | 3 | 4 | 5 | 8 | 0 | 5 | 2 | 3 | 0 | 0 |
| *ALB12* | 0 | 0 | 1 | 2 | 9 | 0 | 0 | 3 | 0 | 1 | 6 | 12 | 0 | 6 | 2 | 4 | 0 | 0 |
| *ALB13* | 0 | 1 | 0 | 1 | 5 | 1 | 0 | 0 | 1 | 1 | 1 | 9 | 1 | 3 | 1 | 3 | 1 | 1 |
| *ALB14* | 0 | 0 | 0 | 0 | 2 | 0 | 0 | 0 | 3 | 1 | 0 | 2 | 0 | 0 | 1 | 2 | 0 | 0 |
| *ALB15* | 0 | 1 | 0 | 4 | 3 | 2 | 0 | 4 | 3 | 1 | 5 | 17 | 0 | 8 | 0 | 7 | 1 | 0 |
| *ALB16* | 0 | 1 | 1 | 2 | 3 | 0 | 0 | 1 | 0 | 0 | 4 | 10 | 0 | 5 | 1 | 4 | 2 | 1 |
| *GLB1* | 0 | 0 | 0 | 1 | 4 | 0 | 0 | 2 | 1 | 6 | 4 | 10 | 1 | 8 | 2 | 6 | 3 | 1 |
| *GLB2* | 0 | 0 | 0 | 4 | 0 | 0 | 0 | 0 | 0 | 0 | 1 | 12 | 1 | 0 | 0 | 1 | 2 | 4 |
| *GLB3* | 0 | 0 | 0 | 1 | 1 | 0 | 0 | 1 | 0 | 0 | 0 | 6 | 0 | 0 | 0 | 1 | 1 | 0 |
| *GLB4* | 0 | 1 | 1 | 4 | 4 | 1 | 0 | 1 | 1 | 0 | 3 | 12 | 0 | 3 | 1 | 3 | 1 | 2 |
| *GLU1* | 0 | 1 | 0 | 3 | 7 | 0 | 0 | 1 | 1 | 0 | 8 | 26 | 0 | 7 | 2 | 10 | 0 | 3 |
| *GLU2* | 0 | 2 | 1 | 7 | 5 | 0 | 0 | 2 | 0 | 0 | 4 | 28 | 0 | 3 | 7 | 11 | 0 | 4 |
| *GLU3* | 0 | 1 | 1 | 2 | 7 | 0 | 0 | 1 | 3 | 0 | 13 | 6 | 1 | 15 | 1 | 3 | 0 | 1 |
| *GLU4* | 0 | 4 | 0 | 2 | 7 | 0 | 0 | 2 | 2 | 1 | 2 | 10 | 1 | 5 | 4 | 2 | 1 | 0 |
| *GLU5* | 0 | 2 | 0 | 0 | 4 | 1 | 0 | 3 | 3 | 4 | 9 | 17 | 0 | 7 | 3 | 5 | 3 | 0 |
| *GLU6* | 0 | 0 | 0 | 1 | 4 | 1 | 0 | 0 | 1 | 0 | 1 | 10 | 0 | 6 | 0 | 5 | 1 | 1 |
| *GLU7* | 0 | 0 | 1 | 1 | 4 | 1 | 0 | 4 | 2 | 5 | 8 | 19 | 0 | 9 | 1 | 6 | 1 | 0 |
| *GLU8* | 0 | 0 | 1 | 1 | 4 | 1 | 0 | 4 | 2 | 5 | 8 | 19 | 0 | 9 | 1 | 6 | 1 | 0 |
| *GLU9* | 0 | 0 | 1 | 1 | 4 | 1 | 0 | 4 | 2 | 5 | 7 | 19 | 0 | 9 | 1 | 6 | 1 | 0 |
| *GLU10* | 0 | 0 | 1 | 0 | 1 | 0 | 0 | 0 | 0 | 2 | 3 | 11 | 0 | 6 | 2 | 1 | 0 | 0 |
| *GLU11* | 0 | 0 | 1 | 0 | 1 | 0 | 0 | 0 | 0 | 1 | 2 | 11 | 0 | 5 | 2 | 1 | 0 | 0 |
| *GLU12* | 0 | 0 | 0 | 3 | 4 | 1 | 0 | 2 | 0 | 3 | 4 | 8 | 0 | 3 | 3 | 3 | 1 | 2 |
| *GLU13* | 0 | 0 | 0 | 5 | 2 | 0 | 0 | 0 | 0 | 2 | 4 | 13 | 0 | 3 | 4 | 3 | 1 | 3 |
| *GLU14* | 0 | 0 | 0 | 1 | 1 | 0 | 0 | 1 | 0 | 0 | 0 | 3 | 0 | 0 | 0 | 0 | 0 | 1 |
| *GLU15* | 0 | 0 | 0 | 4 | 7 | 1 | 0 | 2 | 0 | 2 | 2 | 15 | 0 | 3 | 0 | 4 | 0 | 3 |
| *GLU16* | 1 | 1 | 1 | 2 | 3 | 0 | 0 | 3 | 0 | 2 | 5 | 17 | 0 | 2 | 0 | 5 | 2 | 1 |
| *GLU17* | 0 | 0 | 0 | 0 | 5 | 1 | 0 | 3 | 0 | 2 | 2 | 15 | 1 | 1 | 0 | 5 | 0 | 0 |
| *GLU18* | 0 | 0 | 0 | 1 | 8 | 0 | 0 | 1 | 1 | 0 | 9 | 13 | 1 | 5 | 2 | 6 | 0 | 0 |
| *GLU19* | 0 | 0 | 0 | 0 | 5 | 1 | 0 | 3 | 0 | 2 | 2 | 15 | 1 | 1 | 0 | 5 | 0 | 0 |
| *GLU20* | 0 | 0 | 1 | 2 | 6 | 0 | 0 | 0 | 5 | 2 | 5 | 14 | 1 | 5 | 3 | 7 | 2 | 0 |
| *PRO1* | 0 | 0 | 0 | 3 | 3 | 0 | 0 | 1 | 1 | 0 | 1 | 14 | 0 | 4 | 1 | 2 | 0 | 1 |
| *PRO2* | 0 | 0 | 0 | 1 | 0 | 1 | 0 | 2 | 0 | 0 | 2 | 8 | 1 | 4 | 1 | 3 | 0 | 1 |
| *PRO3* | 0 | 0 | 0 | 6 | 1 | 0 | 0 | 3 | 1 | 0 | 2 | 15 | 0 | 7 | 1 | 3 | 1 | 1 |
| *PRO4* | 0 | 0 | 1 | 4 | 4 | 0 | 0 | 1 | 2 | 0 | 3 | 11 | 0 | 6 | 2 | 4 | 0 | 3 |
| *PRO5* | 0 | 0 | 2 | 8 | 5 | 0 | 0 | 2 | 1 | 0 | 4 | 13 | 0 | 8 | 1 | 3 | 1 | 2 |
| *PRO6* | 0 | 0 | 1 | 0 | 3 | 0 | 0 | 1 | 0 | 1 | 4 | 13 | 0 | 6 | 1 | 4 | 0 | 0 |
| *PRO7* | 0 | 0 | 1 | 4 | 5 | 0 | 0 | 1 | 2 | 0 | 3 | 13 | 0 | 7 | 2 | 5 | 0 | 3 |
| *PRO8* | 0 | 0 | 1 | 1 | 2 | 0 | 0 | 1 | 1 | 0 | 3 | 9 | 1 | 3 | 1 | 3 | 1 | 1 |
| *PRO9* | 0 | 0 | 0 | 2 | 6 | 0 | 0 | 2 | 0 | 0 | 4 | 7 | 0 | 4 | 2 | 2 | 0 | 2 |
| *PRO10* | 0 | 0 | 0 | 2 | 6 | 0 | 0 | 2 | 0 | 0 | 4 | 7 | 0 | 4 | 2 | 2 | 0 | 2 |
| *PRO11* | 0 | 0 | 1 | 5 | 3 | 1 | 0 | 2 | 3 | 0 | 2 | 16 | 1 | 4 | 2 | 6 | 0 | 4 |
| *PRO12* | 0 | 0 | 0 | 0 | 1 | 0 | 0 | 1 | 0 | 0 | 0 | 6 | 1 | 0 | 1 | 2 | 0 | 0 |
| *PRO13* | 0 | 0 | 1 | 3 | 1 | 0 | 0 | 1 | 1 | 0 | 1 | 15 | 1 | 2 | 0 | 3 | 1 | 2 |
| *PRO14* | 0 | 0 | 0 | 0 | 0 | 0 | 0 | 0 | 0 | 0 | 0 | 0 | 0 | 0 | 0 | 0 | 0 | 0 |
| *PRO15* | 0 | 0 | 0 | 0 | 5 | 0 | 0 | 0 | 6 | 2 | 5 | 13 | 0 | 4 | 3 | 6 | 0 | 0 |
| *PRO16* | 0 | 1 | 0 | 1 | 7 | 0 | 1 | 3 | 6 | 2 | 7 | 12 | 0 | 9 | 4 | 5 | 0 | 0 |
| *PRO17* | 0 | 1 | 0 | 0 | 3 | 0 | 0 | 1 | 0 | 2 | 8 | 12 | 1 | 4 | 4 | 3 | 1 | 0 |
| *PRO18* | 0 | 0 | 3 | 4 | 5 | 0 | 0 | 1 | 1 | 3 | 6 | 13 | 1 | 5 | 3 | 4 | 0 | 3 |
| *PRO19* | 0 | 0 | 0 | 6 | 1 | 0 | 0 | 2 | 1 | 0 | 3 | 14 | 0 | 7 | 1 | 3 | 1 | 1 |
| *PRO20* | 0 | 0 | 0 | 6 | 1 | 0 | 0 | 3 | 1 | 1 | 2 | 16 | 0 | 7 | 1 | 3 | 1 | 1 |
| *PRO21* | 0 | 0 | 0 | 12 | 1 | 0 | 0 | 2 | 2 | 0 | 4 | 15 | 0 | 5 | 1 | 5 | 1 | 7 |
| *PRO22* | 0 | 0 | 0 | 0 | 2 | 0 | 0 | 1 | 0 | 0 | 6 | 18 | 0 | 6 | 3 | 5 | 2 | 0 |
| *PRO23* | 0 | 1 | 0 | 1 | 2 | 1 | 0 | 2 | 0 | 0 | 1 | 15 | 0 | 3 | 6 | 5 | 1 | 1 |
| *PRO24* | 0 | 0 | 1 | 1 | 1 | 0 | 0 | 0 | 0 | 1 | 2 | 7 | 0 | 5 | 1 | 2 | 1 | 1 |
| *PRO25* | 0 | 1 | 0 | 4 | 5 | 1 | 0 | 5 | 0 | 0 | 2 | 25 | 1 | 5 | 10 | 7 | 1 | 3 |
| Total occu | 1 | 21 | 26 | 148 | 232 | 20 | 1 | 101 | 69 | 81 | 258 | 768 | 23 | 313 | 119 | 231 | 46 | 72 |

| Gene | 2SSEEDPROTBANAPA | -300CORE | -300ELEMENT | -300MOTIFZMZEIN | AACACOREOSGLUB1 | AACAOSGLUB1 | ACGTABOX | ACGTOSGLUB1 | ACGTSEED3 | AMYBOX1 | AMYBOX2 | ARE1 | CAATBOX1 | CANBNNAPA | CAREOSREP1 | CEREGLUBOX1PSLEGA | CEREGLUBOX2PSLEGA | CEREGLUBOX3PSLEGA | DOFCOREZM |
|---|---|---|---|---|---|---|---|---|---|---|---|---|---|---|---|---|---|---|---|
| *ALB1* | 0 | 0 | 0 | 0 | 0 | 0 | 0 | 0 | 0 | 0 | 0 | 0 | 0 | 0 | 0 | 0 | 0 | 0 | 1 |
| *ALB2* | 0 | 0 | 0 | 0 | 1 | 0 | 0 | 0 | 0 | 0 | 0 | 0 | 30 | 0 | 2 | 0 | 0 | 0 | 5 |
| *ALB3* | 1 | 0 | 2 | 1 | 0 | 0 | 4 | 1 | 0 | 1 | 0 | 0 | 31 | 2 | 0 | 0 | 0 | 1 | 29 |
| *ALB4* | 0 | 0 | 1 | 0 | 1 | 0 | 0 | 0 | 0 | 0 | 0 | 0 | 10 | 1 | 1 | 0 | 0 | 0 | 15 |
| *ALB5* | 1 | 1 | 2 | 0 | 0 | 0 | 4 | 0 | 0 | 1 | 1 | 0 | 23 | 1 | 1 | 0 | 0 | 1 | 27 |
| *ALB6* | 1 | 0 | 1 | 0 | 0 | 0 | 2 | 1 | 0 | 2 | 0 | 0 | 26 | 2 | 0 | 0 | 0 | 0 | 31 |
| *ALB7* | 1 | 0 | 1 | 0 | 0 | 0 | 0 | 1 | 0 | 2 | 0 | 0 | 21 | 1 | 0 | 0 | 0 | 0 | 25 |
| *ALB8* | 0 | 0 | 1 | 0 | 0 | 0 | 0 | 0 | 0 | 1 | 0 | 0 | 15 | 0 | 1 | 0 | 0 | 0 | 24 |
| *ALB9* | 0 | 0 | 1 | 0 | 0 | 0 | 0 | 0 | 0 | 1 | 0 | 0 | 15 | 0 | 1 | 0 | 0 | 0 | 24 |
| *ALB10* | 0 | 0 | 3 | 0 | 1 | 0 | 2 | 1 | 0 | 2 | 0 | 0 | 22 | 0 | 2 | 0 | 0 | 0 | 32 |
| *ALB11* | 2 | 0 | 0 | 0 | 1 | 0 | 2 | 0 | 0 | 2 | 2 | 0 | 24 | 2 | 0 | 1 | 0 | 0 | 37 |
| *ALB12* | 0 | 0 | 3 | 0 | 0 | 0 | 2 | 1 | 0 | 3 | 0 | 0 | 32 | 1 | 0 | 0 | 0 | 0 | 26 |
| *ALB13* | 2 | 0 | 0 | 0 | 1 | 0 | 0 | 0 | 0 | 0 | 0 | 0 | 27 | 4 | 1 | 0 | 0 | 0 | 31 |
| *ALB14* | 0 | 0 | 1 | 0 | 0 | 0 | 0 | 0 | 0 | 0 | 0 | 0 | 5 | 0 | 0 | 0 | 0 | 0 | 8 |
| *ALB15* | 1 | 1 | 2 | 0 | 2 | 0 | 2 | 0 | 0 | 0 | 0 | 0 | 13 | 2 | 1 | 0 | 0 | 0 | 15 |
| *ALB16* | 0 | 1 | 3 | 0 | 1 | 0 | 0 | 0 | 0 | 1 | 0 | 0 | 33 | 0 | 2 | 0 | 0 | 0 | 25 |
| *GLB1* | 0 | 0 | 1 | 0 | 0 | 0 | 0 | 0 | 0 | 0 | 0 | 0 | 8 | 0 | 3 | 0 | 0 | 0 | 32 |
| *GLB2* | 0 | 0 | 1 | 0 | 0 | 0 | 2 | 0 | 0 | 0 | 0 | 0 | 19 | 0 | 2 | 0 | 0 | 0 | 14 |
| *GLB3* | 0 | 0 | 1 | 0 | 0 | 0 | 0 | 1 | 1 | 0 | 0 | 0 | 8 | 0 | 1 | 0 | 0 | 0 | 7 |
| *GLB4* | 0 | 0 | 0 | 0 | 1 | 0 | 2 | 1 | 1 | 1 | 0 | 1 | 13 | 4 | 1 | 0 | 0 | 0 | 19 |
| *GLU1* | 1 | 0 | 2 | 1 | 2 | 0 | 0 | 0 | 0 | 1 | 0 | 0 | 18 | 1 | 0 | 0 | 0 | 0 | 23 |
| *GLU2* | 0 | 1 | 4 | 0 | 2 | 0 | 0 | 0 | 0 | 1 | 2 | 0 | 23 | 2 | 0 | 0 | 0 | 0 | 37 |
| *GLU3* | 1 | 0 | 0 | 0 | 2 | 0 | 0 | 0 | 0 | 0 | 0 | 0 | 21 | 1 | 0 | 0 | 0 | 0 | 26 |
| *GLU4* | 0 | 0 | 2 | 0 | 1 | 0 | 4 | 0 | 0 | 1 | 0 | 0 | 14 | 0 | 1 | 0 | 0 | 0 | 34 |
| *GLU5* | 2 | 1 | 5 | 0 | 3 | 0 | 0 | 0 | 0 | 2 | 3 | 0 | 30 | 3 | 3 | 0 | 0 | 0 | 27 |
| *GLU6* | 0 | 0 | 2 | 0 | 1 | 0 | 2 | 0 | 0 | 1 | 1 | 0 | 34 | 1 | 1 | 0 | 0 | 0 | 30 |
| *GLU7* | 0 | 0 | 3 | 0 | 4 | 1 | 0 | 1 | 0 | 2 | 1 | 0 | 22 | 0 | 1 | 0 | 0 | 0 | 29 |
| *GLU8* | 0 | 0 | 3 | 0 | 4 | 1 | 0 | 1 | 0 | 2 | 1 | 0 | 22 | 0 | 1 | 0 | 0 | 0 | 29 |
| *GLU9* | 0 | 0 | 3 | 0 | 4 | 1 | 0 | 1 | 0 | 2 | 1 | 0 | 22 | 0 | 1 | 0 | 0 | 0 | 29 |
| *GLU10* | 0 | 1 | 3 | 1 | 3 | 0 | 0 | 0 | 0 | 1 | 1 | 0 | 12 | 0 | 0 | 0 | 0 | 0 | 20 |
| *GLU11* | 0 | 1 | 2 | 1 | 3 | 0 | 0 | 0 | 0 | 1 | 1 | 0 | 11 | 0 | 0 | 0 | 0 | 0 | 18 |
| *GLU12* | 0 | 2 | 1 | 0 | 2 | 0 | 0 | 0 | 0 | 4 | 1 | 0 | 19 | 1 | 1 | 0 | 0 | 0 | 34 |
| *GLU13* | 0 | 0 | 1 | 0 | 2 | 0 | 0 | 1 | 0 | 2 | 0 | 0 | 17 | 0 | 1 | 0 | 0 | 0 | 33 |
| *GLU14* | 0 | 0 | 1 | 0 | 0 | 0 | 0 | 0 | 0 | 1 | 1 | 0 | 2 | 2 | 0 | 0 | 0 | 0 | 12 |
| *GLU15* | 0 | 0 | 6 | 0 | 0 | 0 | 2 | 0 | 0 | 0 | 0 | 0 | 19 | 1 | 0 | 0 | 0 | 0 | 18 |
| *GLU16* | 0 | 2 | 3 | 1 | 2 | 0 | 0 | 0 | 0 | 1 | 1 | 0 | 25 | 2 | 2 | 0 | 0 | 0 | 31 |
| *GLU17* | 0 | 0 | 1 | 0 | 1 | 0 | 0 | 0 | 0 | 0 | 0 | 0 | 13 | 0 | 0 | 0 | 0 | 0 | 15 |
| *GLU18* | 1 | 0 | 0 | 0 | 0 | 0 | 2 | 0 | 0 | 1 | 1 | 0 | 21 | 1 | 0 | 0 | 0 | 0 | 24 |
| *GLU19* | 0 | 0 | 1 | 0 | 1 | 0 | 0 | 0 | 0 | 0 | 0 | 0 | 13 | 0 | 0 | 0 | 0 | 0 | 15 |
| *GLU20* | 0 | 1 | 2 | 0 | 2 | 0 | 0 | 0 | 0 | 1 | 0 | 0 | 21 | 0 | 2 | 0 | 0 | 0 | 26 |
| *PRO1* | 0 | 0 | 0 | 1 | 0 | 0 | 0 | 0 | 0 | 4 | 1 | 0 | 21 | 2 | 0 | 0 | 0 | 0 | 23 |
| *PRO2* | 0 | 2 | 1 | 0 | 0 | 0 | 0 | 0 | 0 | 1 | 0 | 0 | 16 | 0 | 0 | 0 | 0 | 0 | 18 |
| *PRO3* | 2 | 0 | 0 | 2 | 2 | 0 | 0 | 0 | 0 | 0 | 1 | 0 | 25 | 3 | 1 | 0 | 0 | 0 | 24 |
| *PRO4* | 2 | 0 | 2 | 2 | 1 | 0 | 2 | 0 | 0 | 0 | 0 | 0 | 17 | 3 | 0 | 0 | 0 | 0 | 27 |
| *PRO5* | 1 | 0 | 0 | 2 | 2 | 0 | 0 | 0 | 0 | 1 | 1 | 0 | 20 | 2 | 1 | 0 | 0 | 0 | 27 |
| *PRO6* | 0 | 0 | 1 | 1 | 0 | 0 | 0 | 0 | 0 | 0 | 0 | 0 | 8 | 1 | 0 | 0 | 0 | 0 | 22 |
| *PRO7* | 2 | 0 | 2 | 2 | 1 | 0 | 2 | 0 | 0 | 0 | 0 | 0 | 19 | 3 | 0 | 0 | 0 | 0 | 27 |
| *PRO8* | 0 | 0 | 1 | 2 | 0 | 0 | 0 | 0 | 0 | 1 | 0 | 0 | 18 | 1 | 1 | 0 | 0 | 0 | 18 |
| *PRO9* | 2 | 0 | 1 | 1 | 0 | 0 | 2 | 0 | 0 | 2 | 0 | 0 | 9 | 3 | 0 | 0 | 0 | 0 | 24 |
| *PRO10* | 2 | 0 | 1 | 1 | 0 | 0 | 2 | 0 | 0 | 2 | 0 | 0 | 13 | 3 | 0 | 0 | 0 | 0 | 24 |
| *PRO11* | 1 | 0 | 2 | 1 | 0 | 0 | 0 | 0 | 0 | 0 | 3 | 0 | 13 | 2 | 0 | 0 | 0 | 0 | 28 |
| *PRO12* | 0 | 0 | 0 | 2 | 0 | 0 | 0 | 0 | 0 | 1 | 1 | 0 | 8 | 1 | 0 | 0 | 0 | 0 | 11 |
| *PRO13* | 1 | 0 | 1 | 2 | 0 | 0 | 0 | 1 | 0 | 1 | 1 | 0 | 14 | 3 | 1 | 0 | 0 | 0 | 24 |
| *PRO14* | 1 | 0 | 0 | 0 | 0 | 0 | 0 | 0 | 0 | 0 | 0 | 0 | 0 | 0 | 0 | 0 | 0 | 0 | 0 |
| *PRO15* | 0 | 0 | 0 | 0 | 0 | 0 | 0 | 1 | 0 | 0 | 1 | 0 | 18 | 1 | 0 | 0 | 0 | 0 | 17 |
| *PRO16* | 0 | 1 | 3 | 2 | 1 | 0 | 2 | 2 | 0 | 1 | 0 | 0 | 23 | 0 | 0 | 0 | 0 | 0 | 19 |
| *PRO17* | 0 | 1 | 4 | 2 | 0 | 0 | 0 | 0 | 0 | 1 | 0 | 0 | 17 | 1 | 1 | 0 | 0 | 0 | 31 |
| *PRO18* | 0 | 1 | 1 | 2 | 1 | 0 | 0 | 0 | 0 | 4 | 0 | 0 | 24 | 1 | 0 | 0 | 2 | 0 | 29 |
| *PRO19* | 2 | 0 | 0 | 2 | 2 | 0 | 0 | 0 | 0 | 0 | 1 | 0 | 25 | 3 | 1 | 0 | 1 | 0 | 21 |
| *PRO20* | 2 | 0 | 0 | 2 | 2 | 0 | 0 | 0 | 0 | 0 | 1 | 0 | 26 | 3 | 1 | 0 | 0 | 0 | 24 |
| *PRO21* | 2 | 0 | 1 | 2 | 3 | 0 | 4 | 2 | 0 | 1 | 1 | 0 | 20 | 3 | 1 | 0 | 0 | 0 | 18 |
| *PRO22* | 1 | 0 | 1 | 2 | 1 | 0 | 0 | 0 | 0 | 1 | 0 | 0 | 21 | 1 | 2 | 0 | 0 | 0 | 18 |
| *PRO23* | 1 | 3 | 4 | 1 | 2 | 0 | 0 | 0 | 0 | 1 | 0 | 0 | 16 | 1 | 1 | 0 | 0 | 0 | 22 |
| *PRO24* | 0 | 0 | 1 | 0 | 0 | 0 | 0 | 0 | 0 | 1 | 0 | 0 | 13 | 0 | 1 | 0 | 0 | 0 | 16 |
| *PRO25* | 1 | 2 | 3 | 1 | 2 | 0 | 0 | 0 | 0 | 0 | 0 | 0 | 23 | 1 | 1 | 0 | 0 | 0 | 25 |
| 0 | 37 | 22 | 99 | 40 | 68 | 3 | 46 | 17 | 2 | 64 | 30 | 1 | 1181 | 77 | 46 | 1 | 3 | 2 | 1474 |

| Gene | DPBFCOREDCDC3 | DRE2COREZMRAB17 | EBOXBNNAPA | EMHVCHORD | GADOWNAT | GARE1OSREP1 | GARE2OSREP1 | GAREAT | GCN4OSGLUB1 | GLMHVCHORD | GLUTAACAOS | GLUTEBOX1OSGT2 | GLUTEBOX1OSGT3 | GLUTEBOX2OSGT2 | GLUTEBOX2OSGT3 | GLUTEBP1OS |
|---|---|---|---|---|---|---|---|---|---|---|---|---|---|---|---|---|
| *ALB1* | 0 | 1 | 2 | 0 | 0 | 0 | 0 | 0 | 0 | 0 | 0 | 0 | 0 | 0 | 0 | 0 |
| *ALB2* | 5 | 2 | 30 | 0 | 0 | 0 | 0 | 0 | 0 | 0 | 0 | 0 | 0 | 0 | 0 | 0 |
| *ALB3* | 2 | 0 | 14 | 0 | 1 | 0 | 1 | 2 | 1 | 2 | 0 | 0 | 0 | 0 | 0 | 0 |
| *ALB4* | 0 | 0 | 10 | 0 | 0 | 0 | 0 | 0 | 0 | 0 | 0 | 0 | 0 | 0 | 0 | 0 |
| *ALB5* | 0 | 1 | 6 | 0 | 0 | 1 | 0 | 0 | 0 | 0 | 0 | 0 | 0 | 0 | 0 | 0 |
| *ALB6* | 3 | 0 | 16 | 0 | 0 | 2 | 0 | 1 | 0 | 0 | 0 | 0 | 0 | 0 | 0 | 0 |
| *ALB7* | 1 | 0 | 12 | 0 | 0 | 2 | 0 | 1 | 0 | 0 | 0 | 0 | 0 | 0 | 0 | 0 |
| *ALB8* | 1 | 1 | 8 | 0 | 0 | 0 | 0 | 3 | 0 | 0 | 0 | 0 | 0 | 0 | 0 | 0 |
| *ALB9* | 1 | 1 | 8 | 0 | 0 | 0 | 0 | 3 | 0 | 0 | 0 | 0 | 0 | 0 | 0 | 0 |
| *ALB10* | 0 | 0 | 6 | 0 | 0 | 0 | 0 | 5 | 0 | 0 | 0 | 0 | 0 | 0 | 0 | 0 |
| *ALB11* | 2 | 0 | 8 | 0 | 0 | 0 | 0 | 2 | 1 | 0 | 0 | 0 | 0 | 0 | 0 | 0 |
| *ALB12* | 0 | 0 | 14 | 0 | 0 | 0 | 0 | 3 | 0 | 0 | 0 | 0 | 0 | 0 | 0 | 0 |
| *ALB13* | 2 | 1 | 20 | 0 | 0 | 0 | 0 | 0 | 0 | 0 | 0 | 0 | 0 | 0 | 0 | 0 |
| *ALB14* | 0 | 0 | 4 | 0 | 0 | 0 | 1 | 0 | 0 | 0 | 0 | 0 | 0 | 0 | 0 | 0 |
| *ALB15* | 4 | 0 | 14 | 0 | 0 | 0 | 0 | 0 | 0 | 0 | 0 | 0 | 0 | 0 | 0 | 0 |
| *ALB16* | 5 | 0 | 24 | 0 | 0 | 0 | 0 | 1 | 1 | 0 | 0 | 0 | 0 | 0 | 0 | 0 |
| *GLB1* | 2 | 1 | 6 | 0 | 0 | 0 | 0 | 0 | 0 | 0 | 0 | 0 | 0 | 0 | 0 | 0 |
| *GLB2* | 6 | 1 | 20 | 0 | 1 | 0 | 0 | 0 | 0 | 0 | 0 | 0 | 0 | 0 | 0 | 0 |
| *GLB3* | 2 | 0 | 6 | 0 | 0 | 0 | 0 | 0 | 0 | 0 | 0 | 0 | 0 | 0 | 0 | 0 |
| *GLB4* | 2 | 0 | 14 | 0 | 0 | 0 | 0 | 1 | 0 | 0 | 0 | 0 | 0 | 0 | 0 | 0 |
| *GLU1* | 2 | 0 | 18 | 0 | 0 | 0 | 0 | 1 | 1 | 2 | 0 | 0 | 0 | 0 | 0 | 0 |
| *GLU2* | 3 | 0 | 22 | 1 | 0 | 0 | 0 | 1 | 2 | 1 | 1 | 1 | 0 | 1 | 0 | 0 |
| *GLU3* | 0 | 1 | 12 | 0 | 0 | 0 | 0 | 0 | 0 | 0 | 0 | 0 | 0 | 0 | 0 | 0 |
| *GLU4* | 2 | 1 | 16 | 0 | 0 | 0 | 0 | 2 | 0 | 0 | 0 | 0 | 0 | 0 | 0 | 0 |
| *GLU5* | 1 | 0 | 16 | 1 | 0 | 0 | 0 | 2 | 1 | 0 | 0 | 0 | 0 | 0 | 0 | 0 |
| *GLU6* | 4 | 0 | 16 | 0 | 0 | 0 | 0 | 1 | 0 | 0 | 0 | 0 | 0 | 0 | 0 | 0 |
| *GLU7* | 2 | 0 | 20 | 0 | 0 | 0 | 0 | 2 | 1 | 0 | 0 | 0 | 0 | 0 | 0 | 0 |
| *GLU8* | 2 | 0 | 20 | 0 | 0 | 0 | 0 | 2 | 1 | 0 | 0 | 0 | 0 | 0 | 0 | 0 |
| *GLU9* | 2 | 0 | 20 | 0 | 0 | 0 | 0 | 2 | 1 | 0 | 0 | 0 | 0 | 0 | 0 | 0 |
| *GLU10* | 1 | 0 | 8 | 1 | 0 | 0 | 0 | 1 | 1 | 2 | 1 | 0 | 0 | 0 | 0 | 0 |
| *GLU11* | 1 | 0 | 6 | 1 | 0 | 0 | 0 | 1 | 1 | 2 | 1 | 0 | 0 | 0 | 0 | 0 |
| *GLU12* | 1 | 0 | 8 | 0 | 0 | 0 | 0 | 5 | 0 | 0 | 0 | 0 | 0 | 0 | 0 | 0 |
| *GLU13* | 1 | 0 | 10 | 0 | 0 | 0 | 0 | 3 | 1 | 0 | 1 | 0 | 1 | 0 | 1 | 1 |
| *GLU14* | 2 | 0 | 4 | 0 | 0 | 1 | 0 | 0 | 0 | 0 | 0 | 0 | 0 | 0 | 0 | 0 |
| *GLU15* | 1 | 0 | 14 | 0 | 0 | 0 | 0 | 0 | 1 | 0 | 0 | 0 | 0 | 0 | 0 | 0 |
| *GLU16* | 2 | 0 | 16 | 0 | 0 | 0 | 1 | 1 | 2 | 2 | 1 | 1 | 0 | 0 | 0 | 0 |
| *GLU17* | 5 | 1 | 26 | 0 | 0 | 0 | 1 | 1 | 0 | 0 | 0 | 0 | 0 | 0 | 0 | 0 |
| *GLU18* | 2 | 1 | 14 | 0 | 2 | 0 | 0 | 1 | 0 | 0 | 0 | 0 | 0 | 0 | 0 | 0 |
| *GLU19* | 5 | 1 | 26 | 0 | 0 | 0 | 1 | 1 | 0 | 0 | 0 | 0 | 0 | 0 | 0 | 0 |
| *GLU20* | 1 | 1 | 14 | 1 | 0 | 0 | 0 | 1 | 0 | 0 | 0 | 0 | 0 | 0 | 0 | 0 |
| *PRO1* | 2 | 0 | 16 | 0 | 0 | 2 | 0 | 2 | 1 | 1 | 0 | 0 | 0 | 0 | 0 | 0 |
| *PRO2* | 2 | 1 | 10 | 1 | 0 | 1 | 0 | 0 | 0 | 0 | 0 | 0 | 0 | 0 | 0 | 0 |
| *PRO3* | 1 | 0 | 18 | 0 | 0 | 0 | 0 | 0 | 2 | 3 | 0 | 0 | 0 | 0 | 0 | 0 |
| *PRO4* | 1 | 0 | 20 | 0 | 2 | 0 | 1 | 2 | 2 | 3 | 0 | 0 | 0 | 0 | 0 | 0 |
| *PRO5* | 1 | 0 | 12 | 0 | 0 | 1 | 0 | 1 | 2 | 3 | 0 | 0 | 0 | 0 | 0 | 0 |
| *PRO6* | 2 | 0 | 10 | 0 | 0 | 0 | 0 | 1 | 1 | 1 | 0 | 0 | 0 | 0 | 0 | 0 |
| *PRO7* | 1 | 0 | 20 | 0 | 2 | 0 | 1 | 2 | 2 | 3 | 0 | 0 | 0 | 0 | 0 | 0 |
| *PRO8* | 3 | 1 | 16 | 0 | 0 | 0 | 1 | 2 | 2 | 3 | 0 | 0 | 0 | 0 | 0 | 0 |
| *PRO9* | 1 | 0 | 16 | 0 | 1 | 0 | 1 | 3 | 1 | 2 | 0 | 0 | 0 | 0 | 0 | 0 |
| *PRO10* | 1 | 0 | 20 | 0 | 1 | 0 | 1 | 3 | 1 | 2 | 0 | 0 | 0 | 0 | 0 | 0 |
| *PRO11* | 2 | 1 | 18 | 0 | 1 | 0 | 1 | 1 | 1 | 1 | 0 | 0 | 0 | 0 | 0 | 0 |
| *PRO12* | 2 | 1 | 6 | 0 | 0 | 0 | 1 | 1 | 2 | 3 | 0 | 0 | 0 | 0 | 0 | 0 |
| *PRO13* | 3 | 1 | 20 | 0 | 1 | 0 | 1 | 2 | 2 | 3 | 0 | 0 | 0 | 0 | 0 | 0 |
| *PRO14* | 0 | 0 | 0 | 0 | 0 | 0 | 0 | 0 | 0 | 0 | 0 | 0 | 0 | 0 | 0 | 0 |
| *PRO15* | 1 | 0 | 22 | 0 | 0 | 0 | 0 | 1 | 1 | 0 | 0 | 0 | 0 | 0 | 0 | 0 |
| *PRO16* | 1 | 0 | 26 | 1 | 0 | 0 | 0 | 2 | 2 | 4 | 0 | 0 | 0 | 0 | 0 | 0 |
| *PRO17* | 1 | 1 | 18 | 1 | 0 | 1 | 0 | 1 | 2 | 3 | 0 | 0 | 0 | 0 | 0 | 0 |
| *PRO18* | 2 | 1 | 18 | 0 | 0 | 2 | 0 | 2 | 2 | 3 | 0 | 0 | 0 | 0 | 0 | 0 |
| *PRO19* | 1 | 0 | 22 | 0 | 0 | 0 | 0 | 1 | 2 | 3 | 0 | 0 | 0 | 0 | 0 | 0 |
| *PRO20* | 1 | 0 | 22 | 0 | 0 | 0 | 0 | 0 | 2 | 3 | 0 | 0 | 0 | 0 | 0 | 0 |
| *PRO21* | 3 | 0 | 26 | 0 | 0 | 1 | 0 | 1 | 2 | 3 | 0 | 0 | 0 | 0 | 0 | 0 |
| *PRO22* | 0 | 0 | 22 | 0 | 0 | 0 | 0 | 1 | 2 | 3 | 0 | 0 | 0 | 0 | 0 | 0 |
| *PRO23* | 0 | 0 | 12 | 2 | 0 | 0 | 1 | 1 | 1 | 1 | 0 | 0 | 0 | 0 | 0 | 0 |
| *PRO24* | 2 | 0 | 10 | 0 | 0 | 0 | 0 | 2 | 0 | 0 | 0 | 0 | 0 | 0 | 0 | 0 |
| *PRO25* | 1 | 1 | 14 | 2 | 0 | 0 | 0 | 0 | 1 | 1 | 0 | 0 | 0 | 0 | 0 | 0 |
| 0 | 113 | 23 | 962 | 12 | 12 | 14 | 14 | 82 | 52 | 63 | 5 | 2 | 1 | 1 | 1 | 1 |

Specific to Albumin
Specific to Globulin
Specific to Glutelin
Specific to Prolamin

Table S7: Distribution and area of concentration of seed-specific cis-elements in *SSP* promoter sequences. The promoter sequence of each SSP has been divided into 100 bp bins labelled from 1-20 (5' to 3'). The distribution of each seed-specific element in each bin was counted. For each bin, the table lists the names of SSPs in front of the corresponding element. Rows for albumins are in pink, globulins are in gray, glutelins are in yellow and prolamins are in green.

| Element | 1 | 2 | 3 | 4 | 5 | 6 | 7 | 8 | 9 | 10 | 11 | 12 | 13 | 14 | 15 | 16 | 17 | 18 | 19 | 20 |
|---|---|---|---|---|---|---|---|---|---|---|---|---|---|---|---|---|---|---|---|---|
| 2SSEEDPROTBANAPA | ALB7 | ALB5 | | | ALB11 ALB15 | | | | ALB13 | | | ALB13 | ALB11 | | | | | ALB3 | | ALB6 |
| 2SSEEDPROTBANAPA | | | | | | | | | | | | | | | | | | | | |
| 2SSEEDPROTBANAPA | | GLU5 | | GLU3 | | | GLU18 | | | | GLU1 | | | | | | | | GLU5 | |
| 2SSEEDPROTBANAPA | PRO4 PRO21 | PRO7 | | | | PRO3 PRO9 | PRO10 | PRO3 PRO20 PRO19 | PRO5 PRO20 PRO21 | PRO11 PRO13 PRO14 PRO19 | PRO4 PRO9 | PRO7 PRO10 | PRO23 | | | | | | PRO22 PRO25 | |
| -300CORE | | | | | | ALB15 | | | | | | | ALB5 | | ALB16 | | | | | |
| -300CORE | | | | | | | | | | | | | | | | | | | | |
| -300CORE | GLU12 | | | | GLU11 | GLU10 GLU12 | | GLU2 | | | GLU16 | | | | | | GLU5 | GLU16 GLU20 | | |
| -300CORE | | | | | | PRO2 | PRO2 | | PRO23 | PRO23 | | | | | PRO17 PRO18 | PRO25 | PRO16 PRO25 | | | |
| -300ELEMENT | | | | ALB3 | ALB14 | | ALB3 | ALB5 ALB6 ALB10 | ALB8 ALB10 | | ALB4 | ALB7 ALB9 ALB10 | | | ALB5 ALB12 ALB16 | ALB15 | ALB16 | | ALB12 ALB16 | |
| -300ELEMENT | GLB3 | | | | | | | GLB1 | | | | | | | | GLB2 | | | | |
| -300ELEMENT | GLU6 GLU14 GLU10 | | GLU15 | GLU15 | GLU6 GLU11 GLU15 | GLU9 GLU10 | GLU1 GLU7 GLU8 | GLU1 GLU2 | GLU20 | | GLU15 | | GLU2 | | GLU4 GLU5 GLU7 GLU8 GLU9 GLU16 | GLU17 GLU19 | GLU4 GLU5 GLU12 GLU15 | GLU2 GLU9 GLU13 GLU16 GLU20 | GLU7 GLU5 GLU8 | GLU5 |
| -300ELEMENT | PRO8 PRO16 | PRO14 PRO24 | | | PRO4 | PRO2 PRO7 PRO17 | PRO14 | PRO21 | PRO23 | PRO4 PRO11 PRO13 PRO14 PRO22 PRO23 | PRO7 PRO9 PRO10 PRO23 | | | PRO11 PRO17 | PRO6 PRO17 PRO18 | PRO25 | PRO16 PRO17 PRO25 | | | PRO16 |
| -300MOTIFZMZEIN | | | | | | | | | | | | | | | GLU1 | | | | ALB3 | |
| -300MOTIFZMZEIN | | | | | | | | | | | | | | | | | | | | |
| -300MOTIFZMZEIN | | | | | | | | GLU11 | | GLU10 | | | | | | | GLU16 | | | |
| -300MOTIFZMZEIN | | | PRO22 | PRO12 | | | PRO12 | | | | | PRO23 | | | PRO1 PRO4 PRO9 PRO16 | PRO3 PRO7 PRO8 PRO10 PRO13 PRO14 | PRO3 PRO5 PRO16 PRO20 PRO21 | PRO4 PRO17 PRO18 PRO19 | PRO5 PRO6 PRO7 PRO8 PRO11 PRO13 PRO14 PRO17 PRO18 PRO19 PRO20 PRO21 PRO22 PRO25 | |
| AACACOREOSGLUB1 | | | | ALB16 | ALB15 | | | ALB15 | | ALB13 | | ALB2 | ALB4 | ALB10 | | | | ALB11 | | |
| AACACOREOSGLUB1 | | | | | | | | | | | | | | GLB4 | | | | | | |
| AACACOREOSGLUB1 | GLU7 GLU8 GLU9 | | GLU11 | GLU10 | | GLU3 GLU11 | | GLU10 GLU20 | GLU11 | GLU10 | GLU1 GLU2 | | GLU16 GLU17 GLU19 | | GLU9 | GLU1 GLU5 GLU7 GLU8 GLU20 GLU12 GLU13 | GLU4 GLU9 | GLU5 GLU7 GLU8 | GLU2 GLU5 GLU7 GLU8 GLU9 GLU12 GLU13 GLU16 | GLU6 |
| AACACOREOSGLUB1 | PRO4 PRO5 PRO21 | PRO7 PRO21 | | | | PRO3 | | PRO19 PRO20 | | | PRO23 | | PRO23 | | | PRO3 PRO16 | PRO5 PRO18 PRO20 PRO21 PRO25 | PRO19 | PRO22 PRO25 | |
| AACAOSGLUB1 | | | | | | | | | | | | | | | | | | | | |
| AACAOSGLUB1 | | | | | | | | | | | | | | | | | | | | |
| AACAOSGLUB1 | | | | | | | | | | | | | | | | | | | GLU7 GLU9 GLU8 | |
| AACAOSGLUB1 | | | | | | | | | | | | | | | | | | | | |
| ACGTABOX | | ALB3 | ALB11 | ALB15 | ALB5 | | | ALB3 | ALB5 | | | ALB10 ALB12 | | | | | | ALB6 | | |
| ACGTABOX | GLB4 | | | | | | | | | | | | | | GLB2 | | | | | |
| ACGTABOX | | | | | | | | GLU4 | | | GLU4 GLU18 | | | GLU6 | | | GLU15 | | | |
| ACGTABOX | PRO13 | | | PRO21 | PRO21 | PRO2 PRO9 | PRO10 | | | | PRO4 | | PRO7 | | | PRO16 | | | | |
| ACGTOSGLUB1 | ALB12 | | | | | | | ALB3 ALB6 | | | | ALB7 | | | | | | | | |
| ACGTOSGLUB1 | | | | | | | | GLB3 | | | | | | | | | | | GLB4 | |
| ACGTOSGLUB1 | | GLU13 | | | | | | | | | | | | | | | | | | |
| ACGTOSGLUB1 | | | | PRO16 PRO21 | | PRO21 | | | | | | PRO16 | | | PRO15 | PRO13 PRO14 | | | GLU7 GLU8 GLU9 | |
| ACGTSEED3 | | | | | | | | | | | | | | | | | | | | |
| ACGTSEED3 | | | | | | | | GLB3 | | | | | | | | | | | GLB4 | |
| ACGTSEED3 | | | | | | | | | | | | | | | | | | | | |
| ACGTSEED3 | | | | | | | | | | | | | | | | | | | | |
| AMYBOX1 | | | ALB8 | ALB6 | | | ALB5 | ALB6 ALB7 | | | | ALB3 ALB7 | | ALB10 | ALB16 | | ALB11 | ALB9 ALB10 ALB11 ALB12 | ALB12 | |
| AMYBOX1 | GLB4 | | | | | | | | | | | | | | | | | | | |
| AMYBOX1 | | GLU14 | GLU11 | GLU10 | | | | | | GLU13 | | | GLU12 | | GLU9 GLU12 | GLU1 GLU4 GLU7 GLU8 | GLU9 GLU18 | GLU5 GLU7 GLU8 GLU12 GLU20 | GLU2 GLU5 GLU12 GLU13 GLU16 | GLU6 |
| AMYBOX1 | PRO1 PRO22 | PRO8 PRO18 | PRO12 | PRO1 | PRO1 PRO18 PRO24 | PRO9 PRO10 | | | | | PRO23 | | | | PRO1 PRO2 PRO13 PRO14 | PRO16 | PRO5 PRO18 PRO21 | PRO9 PRO10 | PRO17 PRO18 | |
| AMYBOX2 | | ALB5 ALB11 | | | | | | | | | | | | | | | | ALB11 | | |
| AMYBOX2 | | | | | | | | | | | | | | | | | | | | |
| AMYBOX2 | | | GLU5 GLU12 | GLU5 | GLU14 | GLU18 | GLU11 | | GLU10 | | GLU5 | | | | GLU6 | | GLU9 | GLU2 GLU7 GLU8 GLU16 | GLU2 | |
| AMYBOX2 | | PRO12 | PRO11 | PRO11 PRO14 | | | | | PRO11 PRO13 PRO14 PRO15 | | | | PRO1 PRO3 | PRO5 PRO21 PRO20 | PRO19 | | | | | |
| ARE1 | | | | | | | | | | | | | | | | | | | | |
| ARE1 | | | | | | | | | GLB4 | | | | | | | | | | | |
| ARE1 | | | | | | | | | | | | | | | | | | | | |
| ARE1 | | | | | | | | | | | | | | | | | | | | |
| CAATBOX1 | ALB2 ALB6 ALB7 ALB8 ALB10 ALB11 ALB12 ALB13 ALB14 | ALB2 ALB6 ALB8 ALB11 ALB12 ALB13 | ALB6 ALB11 ALB12 ALB16 | ALB2 ALB3 ALB4 ALB5 ALB6 ALB8 ALB13 ALB14 ALB15 ALB16 | ALB2 ALB3 ALB4 ALB5 ALB6 ALB7 ALB8 ALB10 ALB11 ALB12 ALB13 ALB14 ALB15 ALB16 | ALB3 ALB5 ALB6 ALB7 ALB9 ALB10 ALB11 ALB12 ALB15 ALB16 | ALB4 ALB5 ALB6 ALB7 ALB10 ALB12 ALB13 ALB16 | ALB4 ALB5 ALB7 ALB11 ALB12 ALB13 ALB15 ALB16 | ALB3 ALB4 ALB5 ALB6 ALB7 ALB8 ALB10 ALB11 ALB12 ALB13 ALB16 | ALB5 ALB8 ALB9 ALB10 ALB13 ALB16 | ALB2 ALB4 ALB5 ALB6 ALB7 ALB8 ALB9 ALB10 ALB13 ALB16 | ALB2 ALB3 ALB6 ALB9 ALB10 ALB16 | ALB2 ALB3 ALB4 ALB5 ALB6 ALB7 ALB10 ALB11 ALB12 ALB13 ALB16 | ALB2 ALB3 ALB11 ALB12 ALB13 | ALB2 ALB3 ALB5 ALB6 ALB7 ALB8 ALB12 ALB15 ALB16 | ALB2 ALB3 ALB7 ALB9 ALB10 ALB11 ALB12 ALB13 ALB16 | ALB2 ALB3 ALB5 ALB6 ALB7 ALB9 ALB10 ALB11 ALB12 ALB13 ALB15 | ALB2 ALB3 ALB5 ALB6 ALB10 ALB12 ALB13 ALB15 ALB16 | ALB2 ALB3 ALB9 ALB10 ALB11 ALB12 ALB13 | ALB2 ALB3 ALB5 ALB6 ALB9 ALB10 ALB11 ALB13 ALB16 |
| CAATBOX1 | GLB2 GLB3 | GLB1 GLB2 GLB3 GLU4 | GLB3 GLB4 | GLB1 | GLB2 | GLB1 GLB2 GLB3 | GLB2 | GLB1 GLB2 | GLB2 GLB3 | GLB2 | GLB2 GLB4 | GLB2 GLB4 | GLB4 | GLB4 | GLB2 | GLB2 GLB4 | GLB2 GLB4 | GLB1 GLB4 | | GLB4 |
| CAATBOX1 | GLU3 GLU5 GLU6 GLU9 GLU10 GLU11 GLU12 GLU15 GLU16 GLU18 GLU20 | GLU2 GLU3 GLU6 GLU7 GLU8 GLU10 GLU11 GLU15 GLU16 GLU18 | GLU2 GLU3 GLU5 GLU6 GLU9 GLU10 GLU12 GLU15 GLU16 GLU17 GLU18 GLU19 GLU20 | GLU1 GLU3 GLU5 GLU6 GLU7 GLU8 GLU9 GLU11 GLU12 GLU15 GLU16 GLU17 GLU19 GLU20 | GLU3 GLU7 GLU8 GLU10 GLU11 GLU12 GLU15 GLU16 GLU18 GLU20 | GLU1 GLU2 GLU6 GLU10 GLU12 GLU13 GLU15 GLU16 GLU18 GLU20 | GLU2 GLU6 GLU9 GLU11 GLU12 GLU15 GLU16 GLU18 GLU20 | GLU2 GLU3 GLU4 GLU6 GLU7 GLU8 GLU9 GLU10 GLU11 GLU12 GLU13 GLU14 GLU16 GLU17 GLU18 GLU19 | GLU4 GLU5 GLU6 GLU7 GLU8 GLU9 GLU10 GLU11 GLU13 GLU14 GLU15 GLU16 GLU17 GLU18 GLU19 | GLU1 GLU2 GLU3 GLU5 GLU6 GLU7 GLU8 GLU10 GLU11 GLU12 GLU13 GLU15 GLU16 GLU17 GLU19 GLU20 | GLU1 GLU2 GLU4 GLU5 GLU6 GLU9 GLU10 GLU12 GLU13 GLU18 | GLU1 GLU2 GLU5 GLU6 GLU7 GLU8 GLU12 GLU13 GLU16 | GLU3 GLU4 GLU5 GLU6 GLU9 GLU12 GLU13 GLU16 | GLU1 GLU2 GLU3 GLU4 GLU5 GLU6 GLU7 GLU8 GLU9 GLU15 GLU17 GLU18 GLU19 | GLU1 GLU3 GLU4 GLU5 GLU6 GLU7 GLU8 GLU9 GLU12 GLU16 GLU20 | GLU1 GLU2 GLU5 GLU6 GLU7 GLU8 GLU12 GLU13 GLU18 GLU20 | GLU1 GLU5 GLU13 GLU15 GLU16 GLU20 | GLU1 GLU2 GLU3 GLU4 GLU6 GLU9 GLU12 GLU13 GLU15 GLU16 GLU17 GLU18 GLU19 | GLU1 GLU2 GLU4 GLU5 GLU6 GLU7 GLU8 GLU9 GLU13 GLU18 GLU20 | GLU1 GLU2 GLU3 GLU4 GLU5 GLU6 GLU7 GLU8 GLU9 GLU13 GLU16 GLU17 GLU19 |
| CAATBOX1 | PRO3 PRO4 PRO7 PRO8 PRO10 PRO13 PRO14 PRO16 PRO17 PRO19 PRO20 PRO22 PRO23 PRO25 | PRO1 PRO3 PRO4 PRO5 PRO7 PRO8 PRO11 PRO12 PRO14 PRO16 PRO18 PRO19 PRO20 PRO22 PRO23 PRO25 | PRO1 PRO3 PRO4 PRO6 PRO7 PRO11 PRO13 PRO15 PRO16 PRO18 PRO19 PRO20 PRO21 PRO22 PRO23 PRO24 PRO25 | PRO1 PRO4 PRO7 PRO8 PRO11 PRO12 PRO14 PRO15 PRO16 PRO19 PRO20 PRO22 PRO23 PRO25 | PRO1 PRO2 PRO3 PRO4 PRO7 PRO9 PRO10 PRO11 PRO12 PRO15 PRO16 PRO17 PRO18 PRO19 PRO20 PRO22 PRO23 PRO24 | PRO1 PRO2 PRO3 PRO7 PRO8 PRO9 PRO13 PRO14 PRO15 PRO16 PRO19 PRO20 PRO21 PRO22 PRO24 PRO25 | PRO1 PRO3 PRO5 PRO8 PRO10 PRO12 PRO13 PRO14 PRO15 PRO16 PRO19 PRO20 PRO22 PRO23 PRO24 PRO25 | PRO3 PRO4 PRO5 PRO12 PRO13 PRO15 PRO16 PRO17 PRO19 PRO20 PRO21 PRO22 PRO24 PRO25 | PRO2 PRO3 PRO4 PRO5 PRO7 PRO8 PRO15 PRO19 PRO20 PRO21 PRO23 PRO25 | PRO3 PRO7 PRO9 PRO10 PRO15 PRO16 PRO17 PRO19 PRO21 PRO23 PRO24 PRO25 | PRO1 PRO2 PRO5 PRO6 PRO8 PRO15 PRO17 PRO18 PRO19 PRO20 PRO21 PRO22 PRO23 | PRO1 PRO2 PRO3 PRO4 PRO5 PRO9 PRO16 PRO18 PRO19 PRO20 PRO21 PRO23 PRO25 | PRO1 PRO3 PRO4 PRO5 PRO7 PRO8 PRO10 PRO11 PRO13 PRO14 PRO15 PRO16 PRO18 PRO20 PRO21 PRO22 PRO25 | PRO3 PRO5 PRO7 PRO9 PRO10 PRO11 PRO13 PRO14 PRO15 PRO16 PRO18 PRO19 PRO20 PRO21 PRO23 | PRO1 PRO2 PRO3 PRO4 PRO5 PRO6 PRO8 PRO9 PRO10 PRO15 PRO16 PRO17 PRO18 PRO20 PRO21 | PRO1 PRO4 PRO5 PRO6 PRO7 PRO8 PRO11 PRO13 PRO14 PRO15 PRO16 PRO17 PRO18 PRO19 PRO20 PRO21 PRO22 PRO24 PRO25 | PRO1 PRO5 PRO6 PRO7 PRO8 PRO9 PRO10 PRO11 PRO13 PRO14 PRO16 PRO17 PRO18 PRO20 PRO21 PRO24 PRO25 | PRO2 PRO3 PRO4 PRO16 PRO17 PRO18 PRO19 PRO22 PRO24 PRO25 | PRO1 PRO2 PRO3 PRO4 PRO6 PRO7 PRO8 PRO9 PRO10 PRO11 PRO13 PRO14 PRO16 PRO22 | PRO2 PRO5 PRO6 PRO7 PRO8 PRO11 PRO13 PRO14 PRO17 PRO18 PRO19 PRO20 PRO21 PRO22 PRO25 |
| CANBNNAPA | ALB7 | ALB5 | | ALB15 | ALB11 ALB15 | | | | ALB13 | ALB4 | | ALB13 | ALB3 ALB11 | | | ALB12 ALB13 | | ALB3 ALB6 | | ALB6 |
| CANBNNAPA | | | | | | | | | | | | GLB4 | | | | | GLB4 | | GLB4 | GLB4 |
| CANBNNAPA | | GLU5 GLU14 | | GLU3 | | | GLU18 | GLU5 GLU14 | GLU12 | | GLU1 GLU2 | GLU16 | GLU16 | GLU2 | | GLU6 | | | GLU5 | GLU15 |

(1-20): In (1-20): Indicates a bin of 100bp each

| | | | | | | | | | | | | | | | | | | | | |
|---|---|---|---|---|---|---|---|---|---|---|---|---|---|---|---|---|---|---|---|---|
| CANBNNAPA | PRO4 PRO21 | PRO7 | | | PRO15 | PRO3 PRO9 | PRO10 PRO12 | PRO3 PRO20 PRO19 | PRO5 PRO20 PRO21 | PRO1 PRO11 PRO13 PRO14 PRO19 | PRO4 PRO9 | PRO7 PRO10 PRO13 PRO14 | PRO23 | | PRO1 | | PRO3 | PRO4 PRO9 | PRO5 PRO6 PRO7 PRO8 PRO10 PRO11 PRO13 PRO17 PRO18 PRO19 PRO20 PRO21 PRO22 PRO25 | |
| CAREOSREP1 | ALB4 | | ALB16 | | ALB9 | | ALB13 | | | | ALB2 ALB16 | ALB10 | | ALB10 | | ALB5 ALB8 ALB15 | | ALB2 | | |
| CAREOSREP1 | GLB4 | | GLB1 | | | | GLB1 GLB2 | GLB3 | GLB1 GLB2 | | | | | | | | | | | |
| CAREOSREP1 | | GLU5 GLU20 | GLU6 GLU13 GLU16 | | | | GLU5 | | | | | GLU9 | GLU7 GLU8 | | GLU16 | | | | GLU4 | GLU5 GLU12 |
| CAREOSREP1 | | | PRO8 PRO17 | PRO24 | | | PRO13 | | PRO3 | | PRO5 PRO20 PRO21 | PRO19 | | PRO22 PRO23 | | | | | | PRO22 PRO25 |
| CEREGLUBOX1PSLEGA | | | | | | | | | | | | | | | | | | ALB11 | | |
| CEREGLUBOX1PSLEGA | | | | | | | | | | | | | | | | | | | | |
| CEREGLUBOX1PSLEGA | | | | | | | | | | | | | | | | | | | | |
| CEREGLUBOX1PSLEGA | | | | | | | | | | | | | | | | | | | | |
| CEREGLUBOX2PSLEGA | | | | | | | | | | | | | | | | | | | | |
| CEREGLUBOX2PSLEGA | | | | | | | | | | | | | | | | | | | | |
| CEREGLUBOX2PSLEGA | | | | | | | | | | | | | | | | | | | | |
| CEREGLUBOX2PSLEGA | | | | | PRO18 | | | | | | | | PRO19 | | | PRO18 | | | | |
| CEREGLUBOX3PSLEGA | | | | | | | ALB3 | ALB5 | | | | | | | | | | | | |
| CEREGLUBOX3PSLEGA | | | | | | | | | | | | | | | | | | | | |
| CEREGLUBOX3PSLEGA | | | | | | | | | | | | | | | | | | | | |
| CEREGLUBOX3PSLEGA | | | | | | | | | | | | | | | | | | | | |
| DOFCOREZM | ALB3 ALB6 ALB8 ALB10 ALB11 ALB13 ALB14 ALB10 | ALB1 ALB3 ALB5 ALB6 ALB7 ALB8 ALB11 ALB13 ALB14 | ALB3 ALB6 ALB8 ALB11 ALB12 ALB14 | ALB2 ALB3 ALB4 ALB5 ALB6 ALB8 ALB11 ALB13 ALB14 ALB15 | ALB3 ALB5 ALB6 ALB7 ALB8 ALB11 ALB12 ALB13 ALB14 | ALB5 ALB6 ALB7 ALB8 ALB10 ALB15 ALB16 | ALB2 ALB3 ALB6 ALB7 ALB8 ALB10 ALB11 ALB12 ALB15 ALB16 | ALB3 ALB4 ALB5 ALB6 ALB7 ALB8 ALB9 ALB10 ALB11 ALB12 ALB13 | ALB3 ALB4 ALB6 ALB7 ALB8 ALB9 ALB10 ALB11 ALB12 ALB13 ALB15 | ALB3 ALB4 ALB7 ALB8 ALB9 ALB10 ALB11 ALB12 ALB13 ALB15 | ALB3 ALB4 ALB5 ALB6 ALB7 ALB8 ALB9 ALB10 ALB11 ALB16 | ALB3 ALB5 ALB6 ALB7 ALB8 ALB9 ALB10 ALB11 ALB12 | ALB3 ALB4 ALB5 ALB6 ALB7 ALB8 ALB9 ALB10 ALB12 ALB13 ALB15 | ALB2 ALB5 ALB6 ALB9 ALB10 ALB11 ALB16 | ALB2 ALB5 ALB6 ALB7 ALB9 ALB10 ALB11 ALB12 ALB13 ALB15 ALB16 | ALB5 ALB7 ALB9 ALB10 ALB11 ALB13 ALB15 ALB16 | ALB3 ALB5 ALB6 ALB7 ALB9 ALB10 ALB11 ALB12 ALB13 ALB15 ALB16 | ALB3 ALB6 ALB9 ALB10 ALB11 ALB12 ALB13 ALB15 ALB16 | ALB2 ALB3 ALB5 ALB9 ALB10 ALB12 ALB13 ALB16 | ALB6 ALB9 ALB10 ALB11 ALB12 ALB13 |
| DOFCOREZM | GLB3 GLB4 | GLB2 GLB3 GLB4 | GLB1 GLB3 | GLB1 GLB2 GLB3 GLB4 | GLB1 GLB2 GLB4 | GLB1 GLB3 GLB4 | GLB4 | GLB1 GLB2 | GLB1 GLB3 | GLB1 GLB2 | GLB1 GLB2 | GLB1 GLB2 GLB4 | GLB1 GLB4 | GLB1 GLB4 | GLB1 GLB2 | GLB1 GLB2 GLB4 | GLB1 | | GLB2 | GLB2 |
| DOFCOREZM | GLU2 GLU3 GLU4 GLU5 GLU6 GLU7 GLU8 GLU10 GLU11 GLU12 GLU14 GLU15 GLU16 GLU17 GLU18 GLU19 GLU20 | GLU2 GLU3 GLU5 GLU6 GLU9 GLU10 GLU11 GLU14 GLU15 GLU18 | GLU2 GLU3 GLU4 GLU6 GLU7 GLU8 GLU9 GLU10 GLU12 GLU13 GLU14 GLU15 GLU17 GLU18 GLU19 | GLU2 GLU3 GLU4 GLU5 GLU7 GLU8 GLU9 GLU11 GLU12 GLU13 GLU14 GLU15 GLU16 GLU17 GLU18 GLU19 GLU20 | GLU1 GLU2 GLU4 GLU5 GLU6 GLU7 GLU8 GLU9 GLU10 GLU11 GLU12 GLU13 GLU15 GLU16 GLU17 GLU18 GLU19 | GLU3 GLU5 GLU9 GLU10 GLU11 GLU12 GLU13 GLU14 GLU15 GLU17 GLU18 GLU19 | GLU1 GLU2 GLU4 GLU6 GLU7 GLU8 GLU9 GLU10 GLU11 GLU12 GLU13 GLU14 GLU16 GLU17 GLU18 GLU19 | GLU2 GLU5 GLU7 GLU8 GLU9 GLU10 GLU11 GLU12 GLU13 GLU14 GLU15 GLU17 GLU19 GLU20 | GLU1 GLU2 GLU3 GLU4 GLU5 GLU6 GLU7 GLU8 GLU9 GLU10 GLU11 GLU12 GLU13 GLU14 GLU15 GLU16 GLU18 GLU20 | GLU1 GLU2 GLU3 GLU4 GLU5 GLU6 GLU7 GLU8 GLU9 GLU10 GLU11 GLU12 GLU13 GLU18 | GLU1 GLU2 GLU5 GLU6 GLU7 GLU8 GLU10 GLU12 GLU13 GLU15 GLU16 GLU20 | GLU2 GLU3 GLU6 GLU9 GLU13 GLU15 | GLU1 GLU2 GLU3 GLU5 GLU7 GLU8 GLU9 GLU12 GLU13 GLU15 GLU16 | GLU1 GLU2 GLU3 GLU4 GLU6 GLU9 GLU13 GLU15 GLU16 GLU17 GLU18 GLU19 | GLU2 GLU4 GLU5 GLU6 GLU7 GLU8 GLU9 GLU12 GLU13 GLU16 GLU18 GLU20 | GLU1 GLU3 GLU4 GLU5 GLU6 GLU9 GLU12 GLU13 GLU17 GLU19 GLU20 | GLU1 GLU2 GLU4 GLU5 GLU6 GLU7 GLU8 GLU9 GLU12 GLU13 GLU16 GLU17 GLU19 GLU20 | GLU2 GLU4 GLU5 GLU6 GLU7 GLU8 GLU9 GLU12 GLU13 GLU16 GLU18 GLU20 | GLU3 GLU4 GLU5 GLU6 GLU7 GLU8 GLU9 GLU12 GLU13 GLU16 GLU17 GLU18 GLU19 GLU20 | GLU1 GLU2 GLU4 GLU5 GLU6 GLU7 GLU8 GLU9 GLU12 GLU13 GLU16 GLU20 |
| DOFCOREZM | PRO1 PRO2 PRO4 PRO5 PRO6 PRO8 PRO14 PRO15 PRO16 PRO17 PRO18 PRO19 PRO20 PRO22 PRO23 PRO24 | PRO2 PRO3 PRO4 PRO5 PRO6 PRO7 PRO8 PRO12 PRO14 PRO17 PRO18 PRO22 PRO24 | PRO1 PRO2 PRO5 PRO6 PRO7 PRO8 PRO11 PRO13 PRO14 PRO16 PRO18 PRO25 | PRO2 PRO3 PRO4 PRO5 PRO9 PRO10 PRO11 PRO12 PRO14 PRO15 PRO16 PRO17 PRO18 PRO20 PRO24 PRO25 | PRO2 PRO4 PRO7 PRO8 PRO9 PRO10 PRO11 PRO15 PRO16 PRO17 PRO18 PRO19 PRO22 PRO23 PRO24 PRO25 | PRO1 PRO2 PRO3 PRO4 PRO6 PRO7 PRO8 PRO9 PRO10 PRO11 PRO12 PRO14 PRO15 PRO16 PRO17 PRO19 PRO20 PRO21 PRO22 PRO23 PRO24 PRO25 | PRO2 PRO3 PRO7 PRO9 PRO10 PRO11 PRO12 PRO13 PRO15 PRO17 PRO20 PRO22 PRO23 PRO24 PRO25 | PRO3 PRO4 PRO5 PRO12 PRO14 PRO15 PRO17 PRO18 PRO19 PRO20 PRO21 PRO23 PRO24 | PRO1 PRO2 PRO3 PRO4 PRO5 PRO7 PRO8 PRO9 PRO10 PRO11 PRO13 PRO14 PRO15 PRO17 PRO18 PRO19 PRO20 PRO21 PRO23 PRO24 | PRO3 PRO4 PRO5 PRO7 PRO9 PRO10 PRO11 PRO13 PRO14 PRO18 PRO20 PRO22 PRO23 | PRO3 PRO5 PRO6 PRO7 PRO9 PRO10 PRO11 PRO13 PRO14 PRO16 PRO17 PRO19 PRO20 PRO21 PRO22 PRO23 PRO25 | PRO1 PRO3 PRO4 PRO5 PRO8 PRO11 PRO13 PRO14 PRO15 PRO17 PRO20 PRO21 PRO23 PRO25 | PRO1 PRO2 PRO4 PRO5 PRO7 PRO8 PRO9 PRO10 PRO15 PRO17 PRO18 PRO19 PRO20 PRO21 PRO23 PRO24 | PRO1 PRO2 PRO5 PRO6 PRO7 PRO8 PRO9 PRO10 PRO11 PRO13 PRO14 PRO15 PRO16 PRO17 PRO19 PRO20 PRO21 PRO22 PRO23 PRO25 | PRO1 PRO2 PRO3 PRO4 PRO6 PRO8 PRO9 PRO16 PRO17 PRO18 PRO21 PRO25 | PRO1 PRO2 PRO3 PRO4 PRO5 PRO7 PRO8 PRO10 PRO11 PRO13 PRO15 PRO16 PRO17 PRO18 PRO20 PRO21 PRO22 PRO25 | PRO1 PRO2 PRO3 PRO4 PRO9 PRO16 PRO17 PRO18 PRO19 PRO20 PRO22 PRO24 PRO25 | PRO3 PRO4 PRO5 PRO6 PRO7 PRO8 PRO9 PRO10 PRO11 PRO13 PRO14 PRO16 PRO17 PRO18 PRO19 PRO20 PRO21 PRO22 PRO24 PRO25 | PRO2 PRO3 PRO4 PRO5 PRO6 PRO7 PRO8 PRO9 PRO10 PRO11 PRO13 PRO14 PRO16 PRO17 PRO18 PRO19 PRO20 PRO21 PRO22 PRO25 | PRO1 PRO5 PRO6 PRO7 PRO8 PRO9 PRO10 PRO11 PRO13 PRO14 PRO17 PRO19 PRO20 PRO21 PRO22 PRO25 |
| DPBFCOREDCDC3 | | ALB2 | ALB2 ALB3 ALB15 | ALB16 | ALB2 ALB13 ALB15 | ALB6 ALB11 ALB15 | | | ALB8 ALB16 | ALB7 | ALB2 | ALB9 ALB13 | | ALB15 | | ALB11 | ALB2 ALB16 | ALB6 | | ALB6 ALB16 |
| DPBFCOREDCDC3 | GLB2 | GLB2 | | | GLB3 | | GLB3 | | GLB1 | | | GLB2 | GLB2 | GLB2 | GLB1 | | GLB4 | GLB2 GLB4 | | |
| DPBFCOREDCDC3 | GLU11 | GLU2 GLU10 GLU14 GLU16 | GLU17 GLU19 | GLU4 | GLU4 GLU7 GLU8 GLU9 GLU17 GLU19 | GLU2 | GLU13 GLU17 GLU19 | | GLU9 | GLU5 GLU7 GLU8 GLU15 | | GLU20 | GLU6 | GLU2 GLU12 | GLU6 GLU18 | GLU6 GLU16 | GLU1 | | GLU18 | GLU1 GLU17 GLU19 |
| DPBFCOREDCDC3 | PRO19 PRO21 PRO24 | | PRO17 | PRO1 PRO6 PRO14 | PRO21 PRO25 | PRO8 PRO12 | PRO8 | PRO2 PRO12 PRO13 PRO14 | PRO2 | | PRO18 PRO24 | | | | PRO15 | PRO1 | PRO18 | PRO11 PRO13 | PRO3 PRO4 PRO16 | PRO5 PRO6 PRO7 PRO8 PRO9 PRO10 PRO11 PRO13 PRO14 PRO20 PRO21 |
| DRE2COREZMRAB17 | ALB1 | ALB9 | | | ALB2 | ALB2 | | | | ALB13 | | | | | | | | | ALB8 | ALB5 |
| DRE2COREZMRAB17 | | | GLB1 | GLB2 | | | | | | | | | | | | | | | | |
| DRE2COREZMRAB17 | | | GLU17 GLU19 | | | | | | GLU3 | | | GLU18 | GLU20 | | | | | | GLU4 | |
| DRE2COREZMRAB17 | PRO12 | PRO11 | PRO13 PRO17 | PRO25 | | | | | | PRO18 | | | | | | | | PRO2 | | |
| EBOXBNNAPA | ALB1 ALB2 ALB3 ALB4 ALB9 ALB13 ALB14 ALB16 | ALB2 ALB7 ALB8 ALB12 | ALB2 ALB5 ALB6 ALB10 ALB11 ALB12 ALB14 ALB15 | ALB3 ALB4 ALB7 | ALB2 ALB4 ALB13 ALB16 | ALB4 ALB12 ALB13 ALB15 ALB16 | ALB2 ALB7 ALB13 ALB15 | ALB2 ALB6 ALB10 ALB12 | | ALB3 ALB10 ALB12 ALB16 | | ALB6 ALB7 ALB8 ALB9 | ALB2 ALB5 ALB12 ALB13 | ALB2 ALB6 ALB11 ALB15 | ALB6 ALB16 | ALB7 ALB16 | ALB2 ALB13 | ALB2 ALB3 ALB15 | ALB5 ALB6 ALB9 ALB13 | ALB3 ALB8 ALB13 ALB16 |
| EBOXBNNAPA | GLB3 | GLB3 GLB4 | GLB2 | GLB2 GLB4 | GLB1 GLB3 | GLB4 | | | GLB1 GLB4 | | GLB2 | GLB2 | GLB2 | GLB2 | | | GLB2 | GLB1 GLB2 GLB4 | GLB4 | |
| EBOXBNNAPA | GLU6 GLU7 GLU8 GLU9 GLU10 GLU11 GLU14 | GLU2 GLU7 GLU8 GLU15 | GLU1 GLU5 GLU7 GLU8 GLU9 GLU10 GLU11 GLU15 GLU17 GLU19 GLU20 | GLU4 GLU9 GLU10 GLU12 GLU18 | GLU1 GLU4 GLU7 GLU8 GLU9 GLU11 GLU14 GLU16 GLU20 | GLU7 GLU8 GLU9 GLU16 GLU17 GLU18 GLU19 | GLU3 GLU4 GLU5 GLU10 GLU15 GLU17 GLU18 GLU19 | GLU3 GLU6 GLU12 GLU13 | GLU2 GLU3 GLU4 GLU5 | GLU1 GLU2 GLU9 GLU12 GLU13 GLU17 GLU18 GLU19 GLU20 | GLU6 GLU7 GLU8 GLU16 GLU17 GLU19 GLU20 | GLU1 GLU3 GLU4 | GLU1 GLU7 GLU8 GLU9 GLU13 GLU17 GLU19 | GLU2 GLU4 GLU5 GLU17 GLU19 | GLU1 GLU6 GLU9 GLU15 GLU17 GLU19 | GLU2 GLU5 GLU7 GLU8 GLU13 GLU15 GLU16 GLU20 | GLU1 GLU5 GLU6 GLU9 GLU18 | GLU1 GLU2 GLU5 GLU7 GLU8 GLU16 | GLU3 GLU6 GLU13 GLU17 GLU18 GLU19 GLU20 | GLU2 GLU4 GLU5 GLU17 GLU18 GLU19 GLU20 |
| EBOXBNNAPA | PRO1 PRO3 PRO4 PRO10 PRO13 PRO14 PRO15 PRO16 PRO18 PRO20 PRO21 PRO22 PRO23 PRO24 | PRO4 PRO5 PRO6 PRO7 PRO13 PRO16 PRO19 PRO20 PRO21 PRO23 PRO24 | PRO3 PRO5 PRO7 PRO8 PRO14 PRO15 PRO17 PRO19 PRO21 PRO25 | PRO1 PRO3 PRO4 PRO7 PRO16 PRO20 PRO21 PRO22 PRO23 PRO24 | PRO2 PRO8 PRO7 PRO11 PRO12 PRO14 PRO15 PRO17 PRO19 PRO21 | PRO1 PRO3 PRO11 PRO12 PRO13 PRO16 PRO18 PRO19 PRO20 PRO21 | PRO3 PRO5 PRO9 PRO12 PRO15 PRO17 PRO21 | PRO1 PRO8 PRO10 PRO16 PRO19 PRO20 PRO25 | PRO1 PRO3 PRO4 PRO9 PRO15 PRO18 PRO19 PRO20 PRO21 PRO23 PRO24 PRO25 | PRO2 PRO4 PRO7 PRO9 PRO10 PRO11 PRO13 PRO14 PRO15 PRO18 PRO22 PRO23 | PRO5 PRO7 PRO10 PRO17 PRO18 PRO20 PRO22 PRO25 | PRO8 PRO16 PRO22 PRO23 | PRO4 PRO9 PRO15 PRO16 PRO17 PRO18 | PRO3 PRO7 PRO9 PRO10 PRO11 PRO13 PRO14 PRO15 | PRO15 PRO16 PRO22 PRO25 | PRO4 PRO5 PRO17 PRO19 PRO20 PRO21 PRO22 | PRO2 PRO3 PRO4 PRO6 PRO7 PRO8 PRO9 PRO10 PRO11 PRO13 PRO14 PRO17 PRO18 | PRO1 PRO2 PRO6 PRO9 PRO10 PRO11 PRO13 PRO15 PRO16 PRO19 PRO22 PRO25 | PRO1 PRO2 PRO5 PRO6 PRO7 PRO8 PRO11 PRO13 PRO14 PRO15 PRO19 PRO20 PRO21 PRO22 | PRO16 |
| EMHVCHORD | | | | | | | | | | | | | | | | | | | | |
| EMHVCHORD | | | | | | | | | | | | | | | | | | | | |
| EMHVCHORD | | | | | GLU11 | GLU10 | | GLU2 | | | | | | | | | GLU5 | GLU20 | | |
| EMHVCHORD | | | | | | PRO2 | | | | PRO23 | | | | | PRO17 | PRO25 | PRO16 PRO25 | | | |
| GADOWNAT | | | | | | | | ALB3 | | | | | | | | | | | | |
| GADOWNAT | | | | | | | | | | | | | | | | | | GLB2 | | |
| GADOWNAT | | | | | | | | | | | | GLU18 | | | | GLU18 | | | | |
| GADOWNAT | | | PRO4 | PRO7 | | | | | | | | | | PRO4 PRO9 | PRO7 PRO10 PRO11 PRO13 PRO14 | | | | | |
| GARE1OSREP1 | | | | ALB6 | | | ALB5 | ALB6 ALB7 | | | | ALB7 | | | | | | | | |

| | | | | | | | | | | | | | | | | | | | | |
|---|---|---|---|---|---|---|---|---|---|---|---|---|---|---|---|---|---|---|---|---|
| GARE1OSREP1 | | | | | | | | | | | | | | | | | | | | |
| GARE1OSREP1 | | GLU14 | | | | | | | | | | | | | | | | | | |
| GARE1OSREP1 | | | | | PRO1<br>PRO18 | | | | | | | | | | PRO1<br>PRO2 | | PRO5<br>PRO21 | | PRO17<br>PRO18 | |
| GARE2OSREP1 | | | ALB14 | | | | | | | | | | | ALB3 | | | | | | |
| GARE2OSREP1 | | | | | | | | | | | | | | | | | | | | |
| GARE2OSREP1 | | | | GLU16 | | | | | | | | | | | | | GLU17<br>GLU19 | | | |
| GARE2OSREP1 | | PRO23 | | PRO12 | | | | | | | | | | | PRO4<br>PRO9 | PRO7<br>PRO8<br>PRO10<br>PRO11<br>PRO13<br>PRO14 | | | | |
| GAREAT | | | ALB8 | | ALB10 | ALB8 | ALB10 | ALB6 | ALB9 | | | ALB3<br>ALB7<br>ALB8 | | ALB10 | ALB9<br>ALB16 | ALB10 | ALB11 | ALB9<br>ALB10<br>ALB11<br>ALB12 | ALB12 | ALB3 |
| GAREAT | GLB4 | | | | | | | | | | | | | | | | | | | |
| GAREAT | | | GLU11 | GLU10 | | | GLU12 | GLU17<br>GLU19 | | GLU13 | | | GLU12 | | GLU9<br>GLU12 | GLU1<br>GLU4<br>GLU7<br>GLU8 | GLU9<br>GLU18 | GLU5<br>GLU7<br>GLU8<br>GLU12<br>GLU13<br>GLU20 | GLU2<br>GLU4<br>GLU5<br>GLU12<br>GLU13<br>GLU16 | GLU6 |
| GAREAT | PRO1<br>PRO22 | PRO6<br>PRO8<br>PRO15<br>PRO18 | PRO12 | PRO1<br>PRO4 | PRO7<br>PRO24 | PRO9<br>PRO10 | | PRO16<br>PRO24 | | PRO5<br>PRO21 | PRO19<br>PRO23 | PRO4<br>PRO9<br>PRO10 | PRO7<br>PRO11<br>PRO13<br>PRO14 | | PRO8<br>PRO13<br>PRO14 | PRO16 | PRO18 | PRO9<br>PRO10 | | |
| GCN4OSGLUB1 | | | | | | | | | | | | | | | | ALB16 | | | ALB3<br>ALB11 | |
| GCN4OSGLUB1 | | | | | | | | | | | | | | | | | | | | |
| GCN4OSGLUB1 | | | GLU15 | | | | | GLU11 | | GLU10 | | | | | GLU1 | GLU2 | GLU16 | GLU5<br>GLU7<br>GLU8<br>GLU9 | GLU2<br>GLU13<br>GLU16 | |
| GCN4OSGLUB1 | | | PRO22 | PRO12 | | | PRO12 | | | PRO15 | | PRO23 | | | PRO1<br>PRO4<br>PRO9<br>PRO16 | PRO3<br>PRO7<br>PRO8<br>PRO10<br>PRO13<br>PRO14 | PRO3<br>PRO5<br>PRO16<br>PRO21<br>PRO20 | PRO4<br>PRO17<br>PRO18<br>PRO19 | PRO5<br>PRO6<br>PRO7<br>PRO8<br>PRO11<br>PRO13<br>PRO14<br>PRO17<br>PRO18<br>PRO19<br>PRO20<br>PRO21<br>PRO22<br>PRO25 | |
| GLMHVCHORD | | | | | | | | | | | | | | | | | | | ALB3 | |
| GLMHVCHORD | | | | | | | | | | | | | | | | | | | | |
| GLMHVCHORD | | | | | | | | GLU11 | | GLU10 | | | | | GLU1 | | GLU16 | | GLU2<br>GLU16 | |
| GLMHVCHORD | | | PRO22 | PRO12 | | | PRO12 | | | | | PRO23 | | | PRO1<br>PRO4<br>PRO9<br>PRO16 | PRO3<br>PRO7<br>PRO8<br>PRO10<br>PRO13<br>PRO14 | PRO3<br>PRO5<br>PRO16<br>PRO20<br>PRO21 | PRO4<br>PRO17<br>PRO18<br>PRO19 | PRO5<br>PRO6<br>PRO7<br>PRO8<br>PRO11<br>PRO13<br>PRO14<br>PRO17<br>PRO18<br>PRO19<br>PRO20<br>PRO21<br>PRO22<br>PRO25 | |
| GLUTAACAOS | | | | | | | | | | | | | | | | | | | | |
| GLUTAACAOS | | | | | | | | | | | | | | | | | | | | |
| GLUTAACAOS | | | | | | | | | GLU11 | GLU10 | | | | | | | | | GLU2<br>GLU13<br>GLU16 | |
| GLUTAACAOS | | | | | | | | | | | | | | | | | | | | |
| GLUTEBOX1OSGT2 | | | | | | | | | | | | | | | | | | | | |
| GLUTEBOX1OSGT2 | | | | | | | | | | | | | | | | | | | | |
| GLUTEBOX1OSGT2 | | | | | | | | | | | | | | | | | | | GLU2<br>GLU16 | |
| GLUTEBOX1OSGT2 | | | | | | | | | | | | | | | | | | | | |
| GLUTEBOX1OSGT3 | | | | | | | | | | | | | | | | | | | | |
| GLUTEBOX1OSGT3 | | | | | | | | | | | | | | | | | | | | |
| GLUTEBOX1OSGT3 | | | | | | | | | | | | | | | | | | | GLU13 | |
| GLUTEBOX1OSGT3 | | | | | | | | | | | | | | | | | | | | |
| GLUTEBOX2OSGT2 | | | | | | | | | | | | | | | | | | | | |
| GLUTEBOX2OSGT2 | | | | | | | | | | | | | | | | | | | | |
| GLUTEBOX2OSGT2 | | | | | | | | | | | | | | | | | | | GLU2 | |
| GLUTEBOX2OSGT2 | | | | | | | | | | | | | | | | | | | | |
| GLUTEBOX2OSGT3 | | | | | | | | | | | | | | | | | | | | |
| GLUTEBOX2OSGT3 | | | | | | | | | | | | | | | | | | | | |
| GLUTEBOX2OSGT3 | | | | | | | | | | | | | | | | | | | GLU13 | |
| GLUTEBOX2OSGT3 | | | | | | | | | | | | | | | | | | | | |
| GLUTEBP1OS | | | | | | | | | | | | | | | | | | | | |
| GLUTEBP1OS | | | | | | | | | | | | | | | | | | | | |
| GLUTEBP1OS | | | | | | | | | | | | | | | | | GLU13 | | | |
| GLUTEBP1OS | | | | | | | | | | | | | | | | | | | | |
| GLUTEBP2OS | | | | | | | | | | | | | | | | | | | | |
| GLUTEBP2OS | | | | | | | | | | | | | | | | | | | | |
| GLUTEBP2OS | | | | | | | | | | | | GLU13 | | | | | | | | |
| GLUTEBP2OS | | | | | | | | | | | | | | | | | | | | |
| GLUTECOREOS | | | | | | | | | | | | | | | | | | | | |
| GLUTECOREOS | | | | | | | | | | | | | | | | | | | | |
| GLUTECOREOS | | | | | | | | | | | | | | | | | | | GLU16 | |
| GLUTECOREOS | | | | | | | | | | | | | | | | | | | | |
| MYBGAHV | | | ALB8 | | | | | | | | | ALB3 | | ALB10 | ALB16 | | ALB11 | ALB10<br>ALB11 | ALB12 | |
| MYBGAHV | GLB4 | | | | | | | | | | | | | | | | | | | |
| MYBGAHV | | | GLU11 | GLU10 | PRO24 | | | | | GLU13 | | | GLU12 | | GLU9<br>GLU12 | GLU1<br>GLU4<br>GLU7<br>GLU8 | GLU9<br>GLU18 | GLU5<br>GLU7<br>GLU8<br>GLU12<br>GLU20 | GLU2<br>GLU5<br>GLU12<br>GLU13<br>GLU16 | GLU6 |
| MYBGAHV | PRO1<br>PRO22 | PRO8<br>PRO18 | PRO12 | PRO1 | | PRO9<br>PRO10 | | | | | PRO23 | | | | PRO13<br>PRO14 | PRO16 | PRO18 | PRO9<br>PRO10 | | |
| NAPINMOTIFBN | | | ALB2 | | | | | | ALB16 | ALB15 | | | | | | | ALB2<br>ALB13 | | | |
| NAPINMOTIFBN | | | | | | | | | | | | | | GLB4 | | | | | | |
| NAPINMOTIFBN | | | | | | | GLU4 | GLU4<br>GLU5 | GLU4 | GLU1 | | GLU3 | | GLU5 | GLU2<br>GLU4 | | GLU16 | | GLU2 | |
| NAPINMOTIFBN | | | PRO17 | | | | | | | PRO23 | | | | | | PRO25 | | PRO16 | | |
| NTBBF1ARROLB | ALB8 | ALB6<br>ALB8 | ALB6 | ALB2<br>ALB8 | ALB12<br>ALB14 | ALB7<br>ALB16 | ALB7<br>ALB8 | | ALB9 | | | ALB8<br>ALB12 | ALB3 | ALB2<br>ALB9 | ALB13 | | ALB9 | ALB6<br>ALB11 | ALB9 | ALB9<br>ALB10<br>ALB11 |
| NTBBF1ARROLB | GLB4 | | | | | | | | | | | | | | | GLB1 | GLB1 | | | |
| NTBBF1ARROLB | | | | GLU2<br>GLU5 | GLU11<br>GLU18 | GLU10<br>GLU12 | GLU1<br>GLU11 | GLU2<br>GLU10 | GLU2<br>GLU13 | | GLU2 | GLU3 | GLU12 | GLU2<br>GLU18 | GLU13 | GLU1<br>GLU4<br>GLU9 | GLU2<br>GLU4<br>GLU5<br>GLU7<br>GLU8<br>GLU13<br>GLU20 | GLU4<br>GLU12<br>GLU20 | GLU4<br>GLU20 | GLU2<br>GLU13 |
| NTBBF1ARROLB | | PRO22 | PRO18<br>PRO25 | PRO12 | PRO25 | PRO1<br>PRO2<br>PRO16<br>PRO25 | PRO11<br>PRO15 | PRO15 | PRO23 | PRO23 | PRO6<br>PRO23<br>PRO25 | PRO4<br>PRO9<br>PRO23 | PRO7<br>PRO10<br>PRO15 | PRO17 | PRO3<br>PRO4<br>PRO9<br>PRO17<br>PRO25 | PRO5<br>PRO7<br>PRO8<br>PRO10<br>PRO11<br>PRO14<br>PRO16<br>PRO20<br>PRO21<br>PRO25 | PRO16<br>PRO17<br>PRO18<br>PRO19<br>PRO22<br>PRO25 | PRO16<br>PRO17<br>PRO18<br>PRO22<br>PRO24<br>PRO25 | | |
| OPAQUE2ZMB32 | | | | | | | | | | | | | | | | | | | | |
| OPAQUE2ZMB32 | | | | | | | | | | | | | | | | | | | | |
| OPAQUE2ZMB32 | | | | | | | | | | | | | | | | | | | | |
| OPAQUE2ZMB32 | | PRO12 | | | | | | | | | | | | | | | | | | |
| POLASIG1 | | ALB3<br>ALB8<br>ALB10 | ALB5<br>ALB6<br>ALB8 | ALB2<br>ALB6<br>ALB8<br>ALB13 | ALB3<br>ALB4<br>ALB5<br>ALB6 | ALB3<br>ALB10<br>ALB11<br>ALB15 | ALB3<br>ALB7<br>ALB8 | ALB5<br>ALB7<br>ALB8<br>ALB15<br>ALB16 | ALB3<br>ALB7<br>ALB8<br>ALB10 | ALB3<br>ALB8<br>ALB12<br>ALB15 | ALB4<br>ALB9<br>ALB10<br>ALB11<br>ALB12 | ALB5<br>ALB6<br>ALB9<br>ALB10 | ALB5<br>ALB9<br>ALB10<br>ALB11<br>ALB15 | ALB9<br>ALB16 | ALB5<br>ALB12<br>ALB13<br>ALB15 | ALB2<br>ALB7<br>ALB15 | ALB9<br>ALB11<br>ALB15 | ALB3<br>ALB9<br>ALB16 | ALB6<br>ALB9<br>ALB11<br>ALB13<br>ALB16 | |
| POLASIG1 | GLB4 | | | | GLB1 | GLB1 | GLB1 | GLB1 | GLB1 | | GLB4 | | | | GLB4 | | | | | |
| POLASIG1 | GLU10<br>GLU13 | GLU3<br>GLU15 | GLU1<br>GLU4<br>GLU11<br>GLU6 | GLU10<br>GLU11 | GLU1<br>GLU5<br>GLU10<br>GLU15<br>GLU18 | GLU2<br>GLU9<br>GLU20 | GLU6<br>GLU7<br>GLU8<br>GLU11 | GLU1<br>GLU3<br>GLU5<br>GLU9<br>GLU10 | GLU5<br>GLU6<br>GLU7<br>GLU8<br>GLU15<br>GLU18<br>GLU20 | GLU1<br>GLU6<br>GLU7<br>GLU8<br>GLU9<br>GLU20 | GLU2<br>GLU6<br>GLU20 | GLU2<br>GLU16 | GLU1<br>GLU3<br>GLU9<br>GLU12 | GLU1<br>GLU3<br>GLU7<br>GLU8<br>GLU9<br>GLU12<br>GLU16<br>GLU17<br>GLU19 | GLU3<br>GLU5<br>GLU7<br>GLU8<br>GLU13 | GLU5 | GLU4<br>GLU7<br>GLU8<br>GLU9<br>GLU12<br>GLU18 | GLU4<br>GLU13 | GLU6<br>GLU9<br>GLU20 | GLU1<br>GLU4<br>GLU7<br>GLU8 |
| POLASIG1 | PRO4<br>PRO6<br>PRO7<br>PRO8<br>PRO22 | PRO7<br>PRO18<br>PRO22 | PRO1<br>PRO6<br>PRO18 | PRO2<br>PRO3<br>PRO5<br>PRO14<br>PRO23 | PRO2<br>PRO9<br>PRO15<br>PRO18<br>PRO24<br>PRO25 | PRO3<br>PRO4<br>PRO10<br>PRO19<br>PRO20<br>PRO24 | PRO4<br>PRO7<br>PRO9<br>PRO10<br>PRO11<br>PRO16 | PRO4<br>PRO5<br>PRO7<br>PRO9<br>PRO10<br>PRO18<br>PRO19<br>PRO20<br>PRO21<br>PRO23<br>PRO24 | PRO3<br>PRO7<br>PRO11<br>PRO13<br>PRO14<br>PRO17<br>PRO24 | PRO15<br>PRO16<br>PRO22<br>PRO25 | PRO3<br>PRO5<br>PRO6<br>PRO16<br>PRO19<br>PRO20<br>PRO21 | PRO3<br>PRO5<br>PRO8<br>PRO20<br>PRO21<br>PRO22<br>PRO23 | PRO1<br>PRO3<br>PRO4<br>PRO15<br>PRO16<br>PRO19 | PRO3<br>PRO5<br>PRO6<br>PRO7<br>PRO9<br>PRO10<br>PRO11<br>PRO13<br>PRO14<br>PRO16<br>PRO17<br>PRO18<br>PRO19<br>PRO20<br>PRO21<br>PRO25 | PRO2<br>PRO6<br>PRO19 | PRO2<br>PRO16<br>PRO17<br>PRO19<br>PRO20 | PRO16 | PRO22<br>PRO24<br>PRO25 | | |
| POLASIG2 | ALB11<br>ALB14 | ALB6 | ALB8 | ALB4<br>ALB5<br>ALB6 | ALB11<br>ALB12 | ALB3<br>ALB6<br>ALB7 | ALB8 | ALB4<br>ALB7 | ALB6 | ALB7 | | ALB11<br>ALB15 | ALB7<br>ALB11 | ALB9 | ALB5<br>ALB10 | | ALB5 | ALB9<br>ALB12 | ALB10<br>ALB12 | ALB13 |
| POLASIG2 | | | | | | | | GLB1 | | | GLB1 | | | | | | | | | |
| POLASIG2 | GLU4<br>GLU11 | GLU2<br>GLU10 | GLU4 | GLU11<br>GLU18 | GLU18 | GLU10 | PRO9<br>PRO10 | | | GLU18 | | GLU5<br>GLU12 | GLU7<br>GLU8<br>GLU9 | GLU1<br>GLU15 | GLU3<br>GLU13<br>GLU18 | GLU3<br>GLU17<br>GLU18<br>GLU19 | | GLU20 | GLU20 | |

| | | | | | | | | | | | | | | | | | | | | |
|---|---|---|---|---|---|---|---|---|---|---|---|---|---|---|---|---|---|---|---|---|
| POLASIG2 | PRO17 | | PRO6 PRO8 PRO15 | PRO24 | | | | PRO16 | PRO8 PRO24 | PRO17 | PRO15 | | PRO5 PRO15 | | PRO6 PRO18 | PRO6 PRO16 | | | | |
| PROLAMINBOXOSGLUB1 | ALB3 | | | | | | | | | | ALB4 | | | | | | | | ALB12 ALB16 | |
| PROLAMINBOXOSGLUB1 | | | | | | | GLB4 | | | | | | | | | | | | | |
| PROLAMINBOXOSGLUB1 | | | GLU3 | | | | | GLU20 | GLU11 | GLU10 | | | GLU2 | | GLU16 | | | GLU9 | GLU7 GLU8 | |
| PROLAMINBOXOSGLUB1 | | PRO24 | PRO5 PRO18 | | PRO4 PRO8 PRO18 | PRO7 | | | | PRO11 PRO13 PRO14 | | | | | PRO6 PRO18 | | | | | |
| PROXBBNNAPA | ALB7 | ALB5 | | | | | | | ALB13 | | | | | | | | | ALB3 | | |
| PROXBBNNAPA | | | | | | | | | | | | | | | | | | | | |
| PROXBBNNAPA | | | | | | | | | | | | | | | | | | | | |
| PROXBBNNAPA | | | | | | | | | | | | | | | | | | | | |
| PYRIMIDINEBOXHVEPB1 | | | ALB6 | | | | ALB7 | | | | | | ALB4 | ALB3 | ALB12 | ALB13 | | | | |
| PYRIMIDINEBOXHVEPB1 | | | | | | | | | | GLB2 | | | | | | GLB4 | | | | |
| PYRIMIDINEBOXHVEPB1 | | | | | | | GLU13 | | | GLU12 | | | | | GLU18 | | | | | |
| PYRIMIDINEBOXHVEPB1 | | PRO18 | | GLU2 GLU16 | | | | | PRO18 | | | | | | | | | | | |
| PYRIMIDINEBOXOSRAMY1A | | ALB11 | ALB3 | ALB3 ALB5 ALB13 | ALB13 | ALB5 | ALB10 | ALB4 ALB5 ALB9 | ALB6 ALB8 | | ALB4 ALB5 | ALB3 ALB5 ALB9 ALB10 ALB11 | ALB8 ALB7 | ALB10 | ALB11 ALB12 | ALB13 | ALB5 | | ALB3 ALB5 | ALB12 |
| PYRIMIDINEBOXOSRAMY1A | | | | | | GLB3 | | | | | GLU15 | | | | | GLB2 | | | | |
| PYRIMIDINEBOXOSRAMY1A | GLU3 GLU6 GLU11 | GLU9 GLU10 | GLU4 GLU7 GLU8 | GLU5 | GLU6 GLU12 | GLU15 | GLU11 GLU14 | GLU10 GLU17 GLU19 GLU20 | GLU11 GLU12 | GLU7 GLU8 GLU9 GLU10 | | GLU3 | GLU16 | GLU1 GLU6 GLU17 GLU19 | GLU2 GLU4 GLU5 GLU6 | | | GLU9 GLU13 GLU20 | GLU5 GLU7 GLU8 GLU12 GLU13 | |
| PYRIMIDINEBOXOSRAMY1A | PRO4 PRO6 | PRO5 PRO7 PRO14 | PRO1 PRO2 PRO6 | | PRO22 | PRO9 PRO10 PRO17 | PRO9 PRO10 PRO13 PRO22 PRO24 | | PRO3 PRO4 PRO9 PRO10 | PRO4 PRO7 | PRO5 PRO7 PRO9 PRO10 PRO19 PRO20 PRO21 PRO23 | | | PRO6 | | PRO16 | PRO25 | PRO16 | | GLU1 GLU4 GLU5 GLU12 GLU13 GLU20 |
| RYREPEATBNNAPA | ALB5 ALB15 | ALB2 | ALB5 ALB8 ALB10 ALB15 | ALB5 | ALB15 | ALB16 | ALB11 | ALB13 | ALB10 | ALB12 | | | ALB12 ALB16 | ALB2 | ALB6 | ALB15 | | ALB5 ALB9 ALB10 | | |
| RYREPEATBNNAPA | | | | GLB3 | GLB1 | GLB4 | | GLB4 | | GLB4 | | | GLB2 | | | | | GLB2 | | |
| RYREPEATBNNAPA | GLU14 | | | | | GLU13 | GLU12 GLU13 | GLU1 GLU2 GLU18 | GLU4 GLU12 | GLU13 | GLU16 | GLU12 GLU20 | GLU2 GLU13 | | GLU1 GLU16 GLU20 | GLU2 GLU9 | GLU3 GLU4 GLU7 GLU8 GLU15 | GLU2 GLU6 | GLU3 | GLU2 GLU15 |
| RYREPEATBNNAPA | PRO19 PRO20 | PRO4 | PRO4 PRO5 PRO18 PRO24 PRO25 | PRO7 PRO8 PRO21 PRO23 | PRO18 PRO21 PRO25 | PRO21 | | PRO3 | PRO1 PRO5 PRO11 PRO20 PRO21 | PRO3 PRO5 PRO11 PRO13 PRO14 PRO18 PRO19 PRO21 PRO25 | | PRO5 PRO19 PRO20 PRO21 | PRO16 | PRO3 PRO4 PRO9 | PRO3 PRO7 PRO10 PRO11 PRO13 PRO14 | PRO2 PRO5 PRO20 PRO21 | PRO1 PRO5 PRO19 PRO20 PRO21 | | PRO3 | |
| RYREPEATGMGY2 | ALB5 | | | | | ALB16 | | | | | | | | ALB2 | ALB6 | | | ALB10 | | |
| RYREPEATGMGY2 | | | | | | | | GLB4 | | | | | | | | | | GLB2 | | |
| RYREPEATGMGY2 | GLU14 | | | | | | GLU12 GLU13 | GLU1 | GLU12 | | | | GLB2 | | GLU1 GLU16 | | | GLU2 GLU6 | | GLU15 GLU2 |
| RYREPEATGMGY2 | | PRO4 | PRO18 PRO25 | PRO7 PRO23 | PRO21 PRO25 | PRO21 | | | PRO1 PRO11 | PRO18 PRO25 | | | GLU2 | PRO3 PRO4 PRO9 | PRO7 PRO10 PRO11 PRO13 PRO14 | PRO2 PRO5 PRO20 PRO21 | PRO19 | | | |
| RYREPEATLEGUMINBOX | ALB5 | | | | | ALB16 | | ALB13 | | | | | | ALB2 | ALB6 | | | ALB5 ALB10 | | |
| RYREPEATLEGUMINBOX | | | | | GLB1 | | | GLB4 | | | | | GLB2 | | | | | GLB2 | | |
| RYREPEATLEGUMINBOX | GLU14 | | | | | GLU13 | GLU12 GLU13 | GLU1 | GLU12 | | | | GLU2 | | GLU1 GLU16 | | GLU3 | GLU2 GLU6 | | GLU2 GLU15 |
| RYREPEATLEGUMINBOX | | PRO4 | PRO5 PRO18 PRO24 PRO25 | PRO7 PRO8 PRO21 PRO23 | PRO25 PRO21 | PRO21 | | | PRO1 PRO11 | PRO18 PRO25 | | | | PRO3 PRO4 PRO9 | PRO7 PRO10 PRO11 PRO13 PRO14 | PRO2 PRO5 PRO20 PRO21 | PRO19 | | | |
| RYREPEATVFLEB4 | | | | | | | | | | | | | | | | | | | | |
| RYREPEATVFLEB4 | | | | | | | | GLB4 | | | | | | | | | | GLB2 | | |
| RYREPEATVFLEB4 | | | | | | | | | | | | | | | GLU1 | | | GLU2 | | GLU15 |
| RYREPEATVFLEB4 | | | PRO18 | | PRO21 | | | | PRO11 | | | | | | | | | | | |
| SEF1MOTIF | | | ALB6 | | | | ALB7 | ALB6 ALB8 ALB15 | ALB5 | | ALB6 ALB10 ALB11 ALB15 | ALB7 ALB11 ALB15 | ALB9 ALB15 | | | ALB7 | | | ALB6 | |
| SEF1MOTIF | | | | | | | | | GLB1 | | | | | | GLB1 | | | | | |
| SEF1MOTIF | | | | | GLU5 GLU12 GLU15 | | | GLU1 GLU4 | GLU15 | | | | | | | | GLU7 GLU8 GLU9 | | | |
| SEF1MOTIF | PRO22 | | PRO15 | PRO3 | PRO2 PRO15 | PRO16 PRO19 PRO20 | | | | PRO15 | | | | PRO6 | | | | | | |
| SEF3MOTIFGM | ALB5 | ALB5 ALB7 | ALB15 ALB12 | ALB6 | ALB9 | | | ALB7 | | ALB5 ALB15 | ALB3 ALB12 | | | ALB15 | ALB11 | ALB8 ALB15 | | | ALB5 | |
| SEF3MOTIFGM | | | | | | | | | | | | | | | | | GLB4 | | | |
| SEF3MOTIFGM | GLU12 | GLU6 | GLU9 | GLU7 GLU8 GLU16 | | GLU18 | | GLU6 GLU12 | GLU4 GLU5 | GLU2 GLU6 GLU16 | | GLU16 | GLU4 GLU20 | GLU13 | GLU13 | GLU3 GLU16 | | GLU15 GLU20 | GLU15 | |
| SEF3MOTIFGM | PRO12 | PRO6 PRO21 PRO22 | | PRO18 PRO22 | PRO12 | | PRO2 PRO18 | PRO8 PRO24 | | PRO8 | PRO4 PRO8 PRO16 | PRO2 PRO7 PRO11 PRO13 PRO14 | PRO23 | PRO15 PRO23 | PRO8 | | PRO13 PRO18 | PRO11 PRO14 | PRO25 PRO22 | PRO22 |
| SEF4MOTIFGM7S | ALB13 ALB14 | ALB10 ALB14 | ALB10 ALB11 ALB13 ALB16 | ALB6 ALB10 | ALB3 ALB15 | ALB3 ALB8 ALB9 ALB16 | | ALB5 | ALB3 ALB5 ALB7 ALB8 | ALB4 ALB8 ALB9 ALB12 | ALB2 ALB3 ALB5 ALB8 ALB9 ALB10 ALB11 ALB12 | ALB4 ALB9 ALB10 ALB12 ALB15 ALB16 | ALB2 ALB15 | ALB5 ALB12 | ALB8 ALB9 ALB13 | ALB5 | ALB12 ALB13 | | ALB2 ALB3 ALB12 ALB13 | ALB11 ALB12 |
| SEF4MOTIFGM7S | | GLB4 | GLB4 | | | GLB3 | GLB1 | | GLB1 | | GLB1 GLB4 | | | | GLB4 | GLB1 | | | | |
| SEF4MOTIFGM7S | GLU5 GLU18 | GLU2 GLU4 | GLU4 | GLU11 GLU14 GLU18 GLU20 | GLU1 GLU3 GLU4 GLU10 GLU13 GLU15 | GLU1 GLU3 GLU12 GLU16 | GLU1 GLU17 GLU19 | GLU17 GLU19 | GLU1 GLU15 GLU18 | GLU2 GLU6 GLU15 | GLU6 GLU15 | GLU2 GLU3 GLU5 GLU7 GLU8 GLU9 GLU16 GLU20 | GLU3 GLU4 GLU7 GLU8 GLU9 GLU12 GLU20 | GLU1 GLU3 GLU5 GLU17 GLU19 GLU20 | GLU3 GLU9 GLU12 GLU18 GLU20 | GLU2 GLU3 GLU7 GLU8 GLU9 GLU17 GLU18 GLU18 GLU19 | GLU2 GLU4 GLU5 GLU7 GLU8 GLU16 | GLU12 GLU15 GLU17 GLU19 | GLU6 GLU18 | GLU6 GLU13 |
| SEF4MOTIFGM7S | PRO16 PRO19 | PRO5 PRO6 PRO7 | PRO25 | PRO5 PRO8 PRO12 PRO15 PRO18 PRO23 | PRO1 PRO4 PRO9 PRO10 PRO16 PRO25 | PRO7 PRO9 PRO10 | PRO7 PRO9 PRO10 | PRO18 PRO24 | PRO15 PRO16 PRO23 | PRO4 PRO9 PRO13 PRO16 PRO25 | PRO7 PRO10 PRO11 PRO15 PRO18 | PRO1 PRO5 PRO15 | PRO5 PRO25 | PRO11 PRO18 | PRO4 PRO6 PRO17 PRO25 | PRO9 PRO10 PRO16 PRO17 PRO22 | PRO6 PRO7 PRO8 PRO11 PRO16 | PRO3 | PRO17 PRO18 PRO22 | PRO5 PRO20 PRO21 |
| SP8BFIBSP8BIB | ALB13 | | | | | ALB15 | | ALB15 | ALB11 | | | | ALB5 | | | ALB6 | | | | |
| SP8BFIBSP8BIB | | | | | | | | | | | | | | GLB4 | | | | | | |
| SP8BFIBSP8BIB | | | | | | GLU12 | | GLU5 | | | | | GLU9 GLU17 GLU19 | GLU7 GLU8 | GLU6 | | | GLU15 | | |
| SP8BFIBSP8BIB | PRO23 | | | PRO2 PRO11 | | | | PRO25 | | | | | | | | | | | | |
| SPHCOREZMC1 | | | | | | | | | | | | | | | | | | | | |
| SPHCOREZMC1 | | | | | | | | | | | | | | | | | | | | |
| SPHCOREZMC1 | | | | | | | GLU13 | | | | | | | | | | | GLU2 | | |
| SPHCOREZMC1 | | | | | | | | | | | | | | | | | | | | |
| SURE1STPAT21 | | | ALB13 | | | | | ALB16 | | | | | | | | ALB2 | | | | |
| SURE1STPAT21 | | | | | | | | | | | | | | | | | | | | |
| SURE1STPAT21 | | | | | GLU11 GLU16 | GLU10 | | | GLU11 | GLU10 | GLU5 | | | | | | | | | |
| SURE1STPAT21 | PRO22 | | | | | | | | | | | | | | | | | | | |
| TATABOX1 | | | | | | | | | | | | | | | | | | | | |
| TATABOX1 | | | | | | | | | | | | | | | | | | | | |
| TATABOX1 | | | | | | | | | | | | | | | | | | | | |
| TATABOX1 | | | | | | | | | | | | | | | | | | | PRO16 | |
| TATABOX2 | ALB11 | ALB3 | ALB11 | | | ALB3 | | ALB6 ALB15 | ALB5 | ALB15 | | ALB7 ALB15 | ALB3 ALB11 | ALB3 ALB11 | | | | ALB12 | ALB3 ALB12 | ALB16 |
| TATABOX2 | GLB4 | | | | GLB1 | | | GLB1 | GLB3 | | | | | | | | | | | |
| TATABOX2 | GLU2 GLU5 GLU9 | GLU7 GLU8 GLU16 | | | GLU5 GLU15 GLU12 | GLU7 GLU8 GLU9 GLU16 | GLU9 GLU14 | GLU4 GLU7 GLU8 | | GLU12 | GLU1 GLU9 | GLU7 GLU8 | GLU17 GLU19 | | | | | GLU17 GLU19 | GLU17 GLU18 GLU19 | GLU2 GLU3 GLU15 GLU16 |
| TATABOX2 | PRO25 | PRO11 PRO23 PRO25 | | PRO2 PRO3 | PRO2 | PRO3 PRO16 PRO19 PRO20 | PRO5 PRO12 PRO20 PRO21 | PRO16 PRO19 PRO25 | PRO9 PRO10 | PRO25 | | | PRO23 | | | PRO1 | | PRO3 PRO4 | PRO9 PRO10 PRO16 | PRO5 PRO6 PRO7 PRO8 PRO11 PRO13 PRO14 PRO17 PRO18 PRO20 PRO21 PRO22 PRO25 |
| TATABOX3 | ALB14 | ALB8 ALB14 | | ALB11 | | | | | ALB15 | | ALB10 | ALB11 | ALB15 | | ALB11 ALB13 | | | | ALB9 | |
| TATABOX3 | | | | | | | | | GLB1 | | | | | | GLB4 | | | | | |
| TATABOX3 | GLU5 | | GLU4 GLU6 GLU20 | | GLU3 GLU20 | | | GLU4 GLU5 GLU9 | GLU7 GLU8 GLU9 | GLU1 GLU18 | | | GLU3 | GLU20 | | GLU3 | | | | |

| | | | | | | | | | | | | | | | | | | | | |
|---|---|---|---|---|---|---|---|---|---|---|---|---|---|---|---|---|---|---|---|---|
| TATABOX3 | | | PRO1 | PRO4 | PRO7<br>PRO11<br>PRO14 | PRO16<br>PRO21 | PRO15<br>PRO16 | PRO15 | PRO3<br>PRO16 | PRO5<br>PRO15<br>PRO16<br>PRO20<br>PRO21 | PRO8<br>PRO11<br>PRO13<br>PRO14<br>PRO19 | PRO15 | PRO15 | | | PRO18 | | | | |
| TATABOX4 | ALB8 | ALB3<br>ALB6 | ALB3<br>ALB11 | ALB5 | ALB14 | ALB7<br>ALB8 | ALB6 | | | | ALB3<br>ALB7<br>ALB11 | | ALB13<br>ALB15 | ALB11 | ALB5<br>ALB9<br>ALB10 | ALB3<br>ALB12 | | | | ALB9<br>ALB10 |
| TATABOX4 | | | | | GLB1 | GLB1 | | | | | | GLB1 | | | | GLB1 | | | | |
| TATABOX4 | GLU9<br>GLU10 | GLU7<br>GLU8 | | | GLU5<br>GLU15 | GLU7<br>GLU9<br>GLU8 | GLU4 | | GLU9 | GLU7<br>GLU8<br>GLU11<br>GLU12 | GLU10<br>GLU13 | | GLU5 | GLU7<br>GLU8<br>GLU9<br>GLU17<br>GLU19 | | | | | GLU9 | GLU5<br>GLU7<br>GLU8<br>GLU12<br>GLU20 |
| TATABOX4 | | PRO16<br>PRO20 | PRO15 | | PRO14<br>PRO18 | GLU13<br>GLU16 | PRO16 | PRO15 | PRO6 | | | | | | | PRO17<br>PRO18<br>PRO24 | | | | |
| TATABOX5 | ALB3<br>ALB10 | ALB10 | ALB8<br>ALB11 | ALB2<br>ALB6<br>ALB13 | ALB4<br>ALB5<br>ALB6<br>ALB10 | ALB3<br>ALB6<br>ALB10 | ALB3<br>ALB12 | ALB7<br>ALB8 | ALB3<br>ALB7<br>ALB8<br>ALB9<br>ALB10 | ALB7<br>ALB9<br>ALB11<br>ALB12 | ALB4<br>ALB8<br>ALB10<br>ALB12 | ALB6<br>ALB8<br>ALB9<br>ALB10<br>ALB11<br>ALB15 | ALB6<br>ALB9<br>ALB10<br>ALB11<br>ALB12<br>ALB15<br>ALB16 | ALB10<br>ALB16 | ALB5<br>ALB10<br>ALB15 | ALB7<br>ALB12<br>ALB15 | ALB7<br>ALB11 | ALB9<br>ALB10 | ALB3<br>ALB6<br>ALB10<br>ALB16 | ALB2 |
| TATABOX5 | GLB2<br>GLB4 | | | | | | GLB1 | GLB1 | GLB1 | | GLB4 | | | GLB4 | | | | | | |
| TATABOX5 | GLU5<br>GLU7<br>GLU8<br>GLU10 | GLU3<br>GLU15 | GLU1<br>GLU4<br>GLU11 | GLU10<br>GLU16<br>GLU18 | GLU1<br>GLU2<br>GLU9<br>GLU13<br>GLU15<br>GLU18 | GLU2<br>GLU5<br>GLU7<br>GLU8<br>GLU11<br>GLU12<br>GLU18 | GLU10 | GLU1<br>GLU2<br>GLU3<br>GLU5<br>GLU9<br>GLU17<br>GLU19 | GLU5<br>GLU7<br>GLU8<br>GLU9<br>GLU20 | GLU16<br>GLU18<br>GLU20 | GLU6<br>GLU13<br>GLU20 | GLU2<br>GLU5<br>GLU16<br>GLU20 | GLU1<br>GLU3<br>GLU9<br>GLU12 | GLU1<br>GLU3<br>GLU7<br>GLU8<br>GLU12<br>GLU16<br>GLU18 | GLU1<br>GLU3<br>GLU5<br>GLU16<br>GLU18 | GLU3<br>GLU5<br>GLU7<br>GLU8<br>GLU9<br>GLU17<br>GLU18<br>GLU19 | GLU4<br>GLU5<br>GLU7<br>GLU8<br>GLU9<br>GLU12<br>GLU18 | GLU13 | GLU1 | |
| TATABOX5 | PRO8<br>PRO22 | PRO8<br>PRO15 | PRO4<br>PRO18<br>PRO22 | PRO3<br>PRO4<br>PRO5<br>PRO7 | PRO2<br>PRO9<br>PRO10<br>PRO24<br>PRO25 | PRO7<br>PRO10<br>PRO19<br>PRO20<br>PRO24 | PRO9<br>PRO10<br>PRO15<br>PRO21 | PRO16<br>PRO21<br>PRO23 | PRO15<br>PRO16<br>PRO17 | PRO3<br>PRO16<br>PRO17<br>PRO22 | PRO15<br>PRO16<br>PRO17 | PRO5<br>PRO19<br>PRO20<br>PRO21<br>PRO22 | PRO16 | PRO4<br>PRO6<br>PRO16<br>PRO17<br>PRO18<br>PRO22<br>PRO25 | PRO6<br>PRO7<br>PRO9<br>PRO10<br>PRO11<br>PRO13<br>PRO14<br>PRO18 | PRO2<br>PRO11<br>PRO15<br>PRO16<br>PRO17 | PRO5<br>PRO18<br>PRO19<br>PRO21 | PRO17<br>PRO18 | PRO1<br>PRO14<br>PRO17<br>PRO18 | |
| TATCCAOSAMY | ALB13 | ALB3<br>ALB5<br>ALB11 | | ALB13 | | ALB8<br>ALB11 | | ALB6 | | ALB10 | | ALB7<br>ALB16 | | | ALB9 | ALB10<br>ALB12 | | ALB11<br>ALB13 | | ALB2 |
| TATCCAOSAMY | | | | | GLB4 | | | | | GLB1 | | | | | | | | | | |
| TATCCAOSAMY | GLU12<br>GLU20 | | GLU12<br>GLU5 | GLU5 | | GLU10<br>GLU18 | GLU11<br>GLU20 | GLU4<br>GLU11 | GLU10<br>GLU12 | | GLU5 | GLU1 | | GLU5 | GLU1<br>GLU6 | | GLU9 | GLU2<br>GLU7<br>GLU8<br>GLU16 | GLU2<br>GLU13 | GLU1 |
| TATCCAOSAMY | PRO11 | PRO12<br>PRO21 | PRO11 | PRO1<br>PRO11<br>PRO14 | GLU11<br>GLU14 | | | PRO4 | PRO7<br>PRO11<br>PRO13<br>PRO14<br>PRO15 | PRO17 | PRO4 | PRO2<br>PRO7<br>PRO9<br>PRO10<br>PRO11<br>PRO13<br>PRO14 | PRO1<br>PRO2<br>PRO3<br>PRO23 | PRO5<br>PRO20<br>PRO21 | PRO19 | | | PRO9 | PRO10<br>PRO17<br>PRO18<br>PRO22<br>PRO25 | |
| TATCCAYMOTIFOSRAMY3D | | ALB3<br>ALB5<br>ALB11 | | | | | | | | | | | | | | | | ALB11 | | |
| TATCCAYMOTIFOSRAMY3D | | | | | | | | | | GLB1 | | | | | | | | | | |
| TATCCAYMOTIFOSRAMY3D | GLU20 | | GLU5<br>GLU12 | GLU5 | GLU14 | GLU18 | GLU11<br>GLU20 | | GLU10 | | GLU5 | GLU1 | | | GLU1<br>GLU6 | | | GLU2<br>GLU7<br>GLU8<br>GLU16 | GLU2<br>GLU13 | |
| TATCCAYMOTIFOSRAMY3D | | PRO12 | PRO11 | PRO11<br>PRO14 | | | | | PRO11<br>PRO13<br>PRO14<br>PRO15 | | | PRO2 | PRO1<br>PRO2<br>PRO3 | PRO5<br>PRO20<br>PRO21 | PRO19 | | GLU9 | | | |
| TGACGTVMAMY | | | ALB11 | ALB15 | ALB4 | ALB4 | | ALB10 | ALB4 | | | ALB12 | ALB6 | | | | ALB7 | | | |
| TGACGTVMAMY | | | | | | | | GLB4 | | | | | | | | | | | | |
| TGACGTVMAMY | | | | GLU1<br>GLU13 | | | | | GLU11 | | GLU10 | GLU1 | | | | GLU13 | | | GLU20 | |
| TGACGTVMAMY | PRO13 | PRO12 | | | | | | PRO23 | | PRO5 | | | PRO11<br>PRO13<br>PRO14 | | PRO25 | | | | | |
| WBOXHVISO1 | | ALB14 | ALB16 | ALB14 | ALB4<br>ALB12 | ALB15 | ALB11<br>ALB13<br>ALB15 | ALB4 | ALB9<br>ALB10<br>ALB12<br>ALB13<br>ALB15 | ALB11<br>ALB16 | ALB2 | ALB8<br>ALB15 | ALB5 | ALB3 | ALB3<br>ALB12<br>ALB15 | ALB3<br>ALB13<br>ALB16 | ALB16 | ALB2 | ALB3<br>ALB6<br>ALB11<br>ALB15 | |
| WBOXHVISO1 | | GLB4 | | GLB4 | | GLB3 | GLB1 | GLB4 | | | | GLB1 | GLB1 | GLB1 | GLB1 | GLB1 | GLB2 | | | |
| WBOXHVISO1 | GLU2<br>GLU20 | GLU2<br>GLU9<br>GLU18 | GLU2<br>GLU4<br>GLU7<br>GLU8<br>GLU15 | GLU1 | GLU1<br>GLU16 | GLU20 | GLU3<br>GLU6<br>GLU12<br>GLU18 | GLU1<br>GLU2<br>GLU5<br>GLU7<br>GLU8<br>GLU9<br>GLU11 | GLU1<br>GLU6<br>GLU7<br>GLU8<br>GLU9<br>GLU16<br>GLU17<br>GLU18<br>GLU19<br>GLU20 | GLU7<br>GLU8<br>GLU10<br>GLU18 | GLU1<br>GLU18 | GLU5<br>GLU15 | GLU2 | GLU2<br>GLU6<br>GLU7<br>GLU8<br>GLU9<br>GLU17<br>GLU19 | GLU1<br>GLU2<br>GLU4<br>GLU5<br>GLU6<br>GLU17<br>GLU19 | GLU2<br>GLU12<br>GLU13<br>GLU16<br>GLU17<br>GLU19<br>GLU20 | GLU1<br>GLU2<br>GLU13<br>GLU16 | GLU1<br>GLU3<br>GLU5<br>GLU7<br>GLU8<br>GLU9<br>GLU12 | GLU2<br>GLU3<br>GLU5<br>GLU13<br>GLU15<br>GLU16<br>GLU17<br>GLU19<br>GLU20 | GLU6<br>GLU20 |
| WBOXHVISO1 | PRO7<br>PRO18<br>PRO23 | PRO6<br>PRO22 | PRO22<br>PRO25 | PRO4<br>PRO11<br>PRO12<br>PRO21 | PRO6<br>PRO7<br>PRO11<br>PRO13<br>PRO23<br>PRO25 | PRO4<br>PRO5<br>PRO8<br>PRO21 | PRO7<br>PRO11<br>PRO12<br>PRO16<br>PRO18<br>PRO23<br>PRO24<br>PRO25 | PRO15<br>PRO24 | PRO15 | PRO15 | PRO3<br>PRO23<br>PRO25 | PRO2<br>PRO23 | PRO2<br>PRO16<br>PRO17<br>PRO19<br>PRO20<br>PRO21<br>PRO25 | | PRO1<br>PRO4<br>PRO6<br>PRO9<br>PRO15<br>PRO16<br>PRO22 | PRO3<br>PRO7<br>PRO8<br>PRO10<br>PRO13<br>PRO14 | PRO2<br>PRO3<br>PRO5<br>PRO16<br>PRO20<br>PRO21 | PRO1<br>PRO4<br>PRO9<br>PRO10<br>PRO17<br>PRO18<br>PRO19<br>PRO22<br>PRO25 | [illegible]<br>PRO6<br>PRO7<br>PRO8<br>PRO11<br>PRO13<br>PRO14<br>PRO17<br>PRO18<br>PRO19<br>PRO20<br>PRO21<br>PRO22<br>PRO25 | |
| WRKY71OS | ALB1<br>ALB3<br>ALB11 | ALB14<br>ALB16 | ALB8<br>ALB10<br>ALB11<br>ALB12<br>ALB15<br>ALB16 | ALB4<br>ALB12<br>ALB13<br>ALB14<br>ALB15<br>ALB16 | ALB2<br>ALB4<br>ALB9<br>ALB12<br>ALB13 | ALB2<br>ALB4<br>ALB5<br>ALB10<br>ALB13<br>ALB15 | ALB4<br>ALB5<br>ALB10<br>ALB11<br>ALB12<br>ALB13<br>ALB15<br>ALB16 | ALB2<br>ALB4<br>ALB8<br>ALB9<br>ALB10<br>ALB12<br>ALB15<br>ALB16 | ALB4<br>ALB8<br>ALB9<br>ALB10<br>ALB11<br>ALB12<br>ALB13<br>ALB15 | ALB2<br>ALB4<br>ALB10<br>ALB11<br>ALB13<br>ALB16 | ALB2<br>ALB4<br>ALB5<br>ALB11<br>ALB13<br>ALB15 | ALB5<br>ALB8<br>ALB9<br>ALB10<br>ALB12<br>ALB15 | ALB5<br>ALB6<br>ALB8<br>ALB9<br>ALB15<br>ALB16 | ALB3<br>ALB6 | ALB2<br>ALB3<br>ALB6<br>ALB12<br>ALB15 | ALB2<br>ALB3<br>ALB8<br>ALB13<br>ALB16 | ALB6<br>ALB7<br>ALB12<br>ALB16 | ALB2<br>ALB5<br>ALB9<br>ALB10<br>ALB11<br>ALB13 | ALB3<br>ALB6<br>ALB11<br>ALB15 | ALB3<br>ALB5 |
| WRKY71OS | GLB1<br>GLB3<br>GLB4 | GLB4 | GLB2 | GLB1<br>GLB4 | GLB2<br>GLB3 | GLB1<br>GLB2<br>GLB3 | GLB1<br>GLB3 | GLB4 | GLB2<br>GLB4 | | GLB2<br>GLB4 | GLB1<br>GLB2 | GLB1<br>GLB4 | GLB1 | GLB1 | GLB1<br>GLB2<br>GLB4 | GLB2 | | GLB2 | |
| WRKY71OS | GLU2<br>GLU17<br>GLU19<br>GLU20 | GLU2<br>GLU9<br>GLU11<br>GLU13<br>GLU14<br>GLU18 | GLU1<br>GLU2<br>GLU4<br>GLU6<br>GLU7<br>GLU8<br>GLU10<br>GLU11<br>GLU12<br>GLU15<br>GLU16<br>GLU17<br>GLU19 | GLU1<br>GLU2<br>GLU4<br>GLU9<br>GLU13<br>GLU20 | GLU1<br>GLU7<br>GLU8<br>GLU9<br>GLU10<br>GLU13<br>GLU14<br>GLU16<br>GLU17<br>GLU19 | GLU2<br>GLU5<br>GLU7<br>GLU8<br>GLU9<br>GLU11<br>GLU13<br>GLU14<br>GLU15<br>GLU18<br>GLU20 | GLU1<br>GLU3<br>GLU5<br>GLU6<br>GLU10<br>GLU11<br>GLU12<br>GLU18 | GLU1<br>GLU2<br>GLU5<br>GLU6<br>GLU7<br>GLU8<br>GLU9<br>GLU10<br>GLU11<br>GLU12 | GLU1<br>GLU2<br>GLU3<br>GLU6<br>GLU7<br>GLU8<br>GLU9<br>GLU10<br>GLU11<br>GLU17<br>GLU18<br>GLU19 | GLU5<br>GLU7<br>GLU8<br>GLU10<br>GLU11<br>GLU15<br>GLU16<br>GLU17<br>GLU18<br>GLU19<br>GLU20 | GLU1<br>GLU3<br>GLU4<br>GLU10<br>GLU16<br>GLU18 | GLU1<br>GLU4<br>GLU5<br>GLU6<br>GLU15<br>GLU17<br>GLU18<br>GLU19 | GLU2<br>GLU15 | GLU1<br>GLU2<br>GLU4<br>GLU6<br>GLU9<br>GLU12<br>GLU13<br>GLU15<br>GLU17<br>GLU19 | GLU1<br>GLU2<br>GLU4<br>GLU5<br>GLU6<br>GLU7<br>GLU8<br>GLU9<br>GLU12<br>GLU16<br>GLU17<br>GLU19<br>GLU20 | GLU2<br>GLU5<br>GLU6<br>GLU7<br>GLU8<br>GLU9<br>GLU12<br>GLU13<br>GLU16<br>GLU17<br>GLU18<br>GLU19<br>GLU20 | GLU1<br>GLU2<br>GLU4<br>GLU5<br>GLU7<br>GLU8<br>GLU9<br>GLU13<br>GLU16<br>GLU17<br>GLU18<br>GLU19<br>GLU20 | GLU1<br>GLU2<br>GLU3<br>GLU5<br>GLU7<br>GLU8<br>GLU9<br>GLU12<br>GLU16<br>GLU17<br>GLU19 | GLU2<br>GLU3<br>GLU5<br>GLU13<br>GLU15<br>GLU16<br>GLU17<br>GLU18<br>GLU19<br>GLU20 | GLU6<br>GLU20 |
| WRKY71OS | PRO1<br>PRO5<br>PRO7<br>PRO13<br>PRO18<br>PRO20<br>PRO22<br>PRO23<br>PRO25 | PRO1<br>PRO3<br>PRO6<br>PRO8<br>PRO9<br>PRO12<br>PRO13<br>PRO14<br>PRO19<br>PRO22<br>PRO25 | PRO1<br>PRO2<br>PRO4<br>PRO6<br>PRO10<br>PRO13<br>PRO18<br>PRO22<br>PRO23<br>PRO25 | PRO3<br>PRO4<br>PRO6<br>PRO7<br>PRO11<br>PRO12<br>PRO19<br>PRO20<br>PRO21<br>PRO22<br>PRO25 | PRO1<br>PRO6<br>PRO7<br>PRO11<br>PRO12<br>PRO13<br>PRO16<br>PRO17<br>PRO23<br>PRO24<br>PRO25 | PRO3<br>PRO4<br>PRO5<br>PRO8<br>PRO12<br>PRO15<br>PRO18<br>PRO19<br>PRO20<br>PRO21<br>PRO22 | PRO1<br>PRO4<br>PRO7<br>PRO8<br>PRO9<br>PRO11<br>PRO12<br>PRO13<br>PRO14<br>PRO16<br>PRO17<br>PRO18<br>PRO21<br>PRO23<br>PRO24<br>PRO25 | PRO1<br>PRO3<br>PRO4<br>PRO7<br>PRO8<br>PRO10<br>PRO11<br>PRO13<br>PRO15<br>PRO19<br>PRO20<br>PRO23<br>PRO24 | PRO1PR<br>O2PRO3<br>PRO5PR<br>O6PRO7<br>PRO11P<br>RO13PR<br>O14PRO<br>15PRO1<br>8PRO20<br>PRO21P<br>RO23PR<br>O25 | PRO5<br>PRO15<br>PRO16<br>PRO17<br>PRO18<br>PRO19<br>PRO20<br>PRO23<br>PRO24 | PRO1<br>PRO3<br>PRO6<br>PRO17<br>PRO20<br>PRO21<br>PRO22<br>PRO23<br>PRO24<br>PRO25 | PRO2<br>PRO3<br>PRO6<br>PRO16<br>PRO17<br>PRO18<br>PRO23<br>PRO25 | PRO2<br>PRO4<br>PRO5<br>PRO6<br>PRO11<br>PRO13<br>PRO14<br>PRO16<br>PRO17<br>PRO19<br>PRO20<br>PRO21<br>PRO22<br>PRO25 | PRO3<br>PRO4<br>PRO5<br>PRO8<br>PRO9<br>PRO11<br>PRO13<br>PRO14<br>PRO19<br>PRO20<br>PRO21<br>PRO22<br>PRO25 | PRO1<br>PRO4<br>PRO5<br>PRO6<br>PRO7<br>PRO9<br>PRO10<br>PRO11<br>PRO13<br>PRO14<br>PRO15<br>PRO16<br>PRO22<br>PRO24<br>PRO25 | [illegible]<br>PRO4<br>PRO5<br>PRO6<br>PRO7<br>PRO8<br>PRO9<br>PRO10<br>PRO11<br>PRO13<br>PRO14<br>PRO16<br>PRO17<br>PRO19<br>PRO20<br>PRO21<br>PRO22<br>PRO25 | PRO1<br>PRO2<br>PRO3<br>PRO5<br>PRO6<br>PRO7<br>PRO8<br>PRO11<br>PRO13<br>PRO14<br>PRO16<br>PRO17<br>PRO20<br>PRO21<br>PRO22<br>PRO25 | PRO1<br>PRO4<br>PRO5<br>PRO9<br>PRO10<br>PRO15<br>PRO17<br>PRO18<br>PRO19<br>PRO20<br>PRO21<br>PRO22<br>PRO25 | PRO5<br>PRO6<br>PRO7<br>PRO8<br>PRO11<br>PRO13<br>PRO14<br>PRO15<br>PRO17<br>PRO18<br>PRO19<br>PRO20<br>PRO21<br>PRO22<br>PRO25 | PRO15 |

**Table S8. Unique probe sets with respective MSU IDs and gene names of 36 SSPs are listed with their fold change values in five seed developmental stages (S1-S5).**

| SSP | LOC | Probe set | S1 | S2 | S3 | S4 | S5 |
|---|---|---|---|---|---|---|---|
| *ALB1* | LOC_Os03g55730 | Os.11614.1.S1_s_at | -1.03475 | -1.26476 | -1.64841 | -1.654157 | -1.238751 |
| *ALB2* | LOC_Os03g55734 | Os.13536.1.S1_at | -0.07126 | 9.647662 | 9.136682 | 7.8231458 | 2.6143481 |
| *ALB4* | LOC_Os05g41970 | Os.10338.1.S1_x_at | -0.05443 | 11.30982 | 11.98644 | 12.197425 | 12.2312 |
| *ALB5* | LOC_Os07g11310 | Os.55229.1.S1_s_at | -0.16641 | 3.185514 | 9.469239 | 10.139349 | 10.540231 |
| *ALB6* | LOC_Os07g11320 | OsAffx.16210.1.S1_at | -0.01985 | 0.229898 | 2.624386 | 4.7061758 | 5.1677227 |
| *ALB7* | LOC_Os07g11330 | Os.8224.1.S1_at | -0.05738 | 9.475899 | 12.02263 | 12.306412 | 12.396999 |
| *ALB8* | LOC_Os07g11360 | Os.12884.1.S1_at | -0.12369 | 7.206985 | 11.64983 | 12.198414 | 12.320983 |
| *ALB9* | LOC_Os07g11380 | Os.5529.5.A1_at | -0.01855 | -0.00208 | 0.286129 | 0.8126423 | 0.4190397 |
| *ALB10* | LOC_Os07g11410 | Os.7449.2.S1_x_at | -0.07941 | 9.288882 | 11.95109 | 12.302156 | 12.375307 |
| *ALB11* | LOC_Os07g11510 | Os.39040.3.S1_x_at | 0.021761 | 11.38038 | 12.38187 | 12.45132 | 12.49292 |
| *ALB12* | LOC_Os07g11630 | Os.6445.1.S1_x_at | 0.10267 | 6.106025 | 8.181385 | 8.5129748 | 8.5278893 |
| *ALB13* | LOC_Os07g11650 | Os.8005.2.S1_x_at | -1.54407 | 8.900621 | 9.491825 | 9.6409245 | 9.6480274 |
| *ALB14* | LOC_Os07g12080 | Os.5494.1.S1_at | -0.30259 | 7.177462 | 9.382535 | 10.366676 | 10.688448 |
| *ALB15* | LOC_Os07g12090 | OsAffx.16221.1.A1_s_at | -0.0302 | 1.605257 | 3.075981 | 2.6674123 | 1.4675654 |
| *ALB16* | LOC_Os11g33000 | Os.11519.1.S1_a_at | 0.010641 | 12.42583 | 12.14739 | 12.104479 | 12.088413 |
| *GLB1* | LOC_Os03g10110 | Os.6372.1.S1_at | -0.68343 | 7.644902 | 10.32048 | 10.661998 | 10.788648 |
| *GLB3* | LOC_Os03g46100 | Os.5503.2.S1_at | -0.17854 | 6.999302 | 9.534412 | 10.172911 | 10.11273 |
| *GLB4* | LOC_Os03g57960 | Os.8208.1.S1_at | 0.626856 | 8.819532 | 11.54093 | 11.596232 | 11.634596 |
| *GLU1* | LOC_Os01g55630 | OsAffx.11615.1.S1_at | -0.0857 | 4.307417 | 7.45145 | 7.8700667 | 6.1748448 |
| *GLU2* | LOC_Os01g55690 | Os.22346.1.S1_x_at | -0.25063 | 11.12851 | 11.37265 | 11.391493 | 11.388573 |
| *GLU3* | LOC_Os01g74480 | Os.18214.1.S1_at | 0.03281 | 0.026125 | 0.038729 | -0.016479 | 0.0264409 |
| *GLU4* | LOC_Os02g14600 | Os.38686.1.S1_at | -0.02315 | 10.71269 | 11.15177 | 11.189519 | 10.981429 |
| *GLU5* | LOC_Os02g15070 | Os.13907.1.S1_at | -0.10272 | 10.85664 | 11.92767 | 11.874043 | 11.179099 |
| *GLU6* | LOC_Os02g15090 | Os.14012.1.S1_at | -0.10181 | 11.004 | 11.89669 | 11.920729 | 11.764939 |
| *GLU12* | LOC_Os02g25640 | Os.9822.3.S1_x_at | -0.06743 | 11.94787 | 12.52192 | 12.461752 | 12.477262 |
| *GLU13* | LOC_Os02g25860 | Os.20396.1.A1_s_at | -0.09858 | 8.100305 | 9.8729 | 9.4340087 | 3.9645734 |
| *GLU14* | LOC_Os03g31360 | Os.8013.1.S1_at | -1.09894 | 10.48669 | 10.61918 | 10.512416 | 10.276435 |
| *GLU15* | LOC_Os05g02520 | Os.28047.1.S1_at | 2.323789 | 2.460086 | 2.545184 | 1.8328743 | -0.790479 |
| *GLU16* | LOC_Os08g03410 | Os.12816.1.S1_at | 3.820055 | 4.194497 | 9.935343 | 10.804815 | 10.69122 |
| *GLU18* | LOC_Os09g37967 | Os.55779.1.S1_at | -0.07211 | -0.06966 | -0.05827 | 0.0337545 | 0.0613673 |
| *GLU19* | LOC_Os09g37976 | Os.55742.1.S1_at | 2.961718 | 2.800813 | 2.924411 | 1.4490619 | 1.7686415 |
| *GLU20* | LOC_Os10g26060 | Os.12807.1.S1_x_at | -1.46994 | 9.926479 | 10.32753 | 10.165247 | 10.138838 |
| *PRO15* | LOC_Os06g31060 | Os.5517.1.S1_at | -0.08799 | 7.341345 | 10.54316 | 10.895781 | 10.819704 |
| *PRO16* | LOC_Os06g31070 | Os.5918.1.S1_at | -0.09413 | 11.61038 | 12.18646 | 12.257232 | 12.371306 |
| *PRO17* | LOC_Os07g10570 | Os.11099.2.S1_x_at | -0.03993 | 11.84916 | 12.9172 | 12.937485 | 12.963907 |
| *PRO18* | LOC_Os07g10580 | Os.8177.1.S1_a_at | -0.06204 | 11.19938 | 11.74155 | 11.706398 | 11.753168 |

**Table S9. List of primer sequences used for qRT-PCR analysis of rice *SSP*.**

| S.No. | LOCUS ID | SSP name | Forward Primer | Reverse Primer |
|---|---|---|---|---|
| 1 | LOC_Os03g55740 | ALB3 | AAACTTTGGGCATGGGTAGCT | GCAGGAGAGCAATTGGTTGTG |
| 2 | LOC_Os02g15150 | GLU7 | GGTAACAACAATAGGGTTCAACAAG | TCTGATCATTTTGGCTCTGCAG |
| 3 | LOC_Os02g15169/ LOC_Os02g15178 | GLU8,9 | CAATTGTTGAAACCGACTTTGAC | GTTGTATGAATCAGCACGACTAGGA |
| 4 | LOC_Os02g16820/ LOC_Os02g16830 | GLU10,11 | CAACAGGGAGCAACAAATGTATG | AACAGTTTAAGGCCATTCTTTACC |
| 5 | LOC_Os05g26240 | PRO1 | GGGACGGGGCCAAGGTGAGGAAA | TACTCCCTCCGTTTCACCTTCCG |
| 6 | LOC_Os05g26359 | PRO2 | CCACCACCACAAGCCATCACCGCC | AAAGGGGCGCTGGTGCTCCGGT |
| 7 | LOC_Os05g26350 | PRO3 | GCAACAGCTCAGGCTGGTGGCGC | CTGCTGGAGGTGTAGCTGCTGCGC |
| 8 | LOC_Os05g26368 | PRO4 | GTGGCGCAACAATCTCACTATC | ACTGGAGTTGTAGCTGCTGCG |
| 9 | LOC_Os05g26377/ LOC_Os05g26620/ LOC_Os05g26460 | PRO5,7,10 | GTGGCGCAACAATCTCACTATC | GGTGCACCATAGTACCTAGGGTAG |
| 10 | LOC_Os05g26480 | PRO8 | CATGTTTTAGCTGTGTTTCAACTGA | AACAATGTTAATGTCCTGATAGTGAGA |
| 11 | LOC_Os05g26690 | PRO11 | ATGTTTTAGCTGCGTTTCAACTG | TTGTAGCTGCTGCGCTATGG |
| 12 | LOC_Os05g26720 | PRO12 | GATCGTTCACACAGTTCAAGCAT | CAGCTGATATTGCCTATAACTTTGAC |
| 13 | LOC_Os05g26750 | PRO13 | GGTGGGTGGAGGGCGACGTTTGC | GCTCCCGCCACCGCGAGATGA |
| 14 | LOC_Os05g26770 | PRO14 | TTCAAGCATTATACAGTAAAAAAGAAAG | CAGCTGATATTGCCTATAACTTTGAC |
| 15 | LOC_Os07g11900/ LOC_Os07g11910/ LOC_Os07g11920 | PRO19,20,21 | TTCGTCTTTGCTCTCCTTGCTA | GAGAGGCGACTGCACCTGAT |
| 16 | LOC_Os12g16880 | PRO22 | CCCTGTAGCGCATTCGACGGTCGGT | CGGCCTGGCAGTGAGACTGTTGCG |
| 17 | LOC_Os12g16890 | PRO23 | ATCACCCGTGTTTCAACTGAGA | TCGAAGTAGACGCTAGCAAACTG |
| 18 | LOC_Osg12g17010 | PRO24 | TCATCGTGTGTGACAAGTTGAAAC | CTTAGAACCAAATGTAGGAACACATC |
| 19 | LOC_Osg12g17030 | PRO25 | GCGCAGCTACAGCTACAACAGT | CTACAGGGAGCAGTGTTGTAGCTT |

Table S10. Fold change values of *SSPs* in five stages of rice seed

| SSP | S1 | S2 | S3 | S4 | S5 |
|---|---|---|---|---|---|
| *ALB3* | 2.287714 | 8.441406 | 8.895065 | 9.904669 | 11.98704 |
| *GLU7* | 0.078281 | 7.405641 | 9.868073 | 12.91975 | 12.78957 |
| *PRO1* | 2.140931 | -2.076881 | 0.338745 | 2.438862 | 10.51609 |
| *PRO2* | -1.716179 | -3.054952 | -1.267264 | 2.027692 | 5.241789 |
| *PRO3* | 1.427234 | 2.129469 | 4.767318 | 9.480331 | 4.604453 |
| *PRO4* | -2.831367 | -0.458904 | 8.841946 | 13.26569 | 13.67402 |
| *PRO8* | -3.004018 | -0.520239 | 10.49985 | 14.79457 | 15.00255 |
| *PRO11* | -0.948499 | 1.394483 | 11.70157 | 15.03512 | 15.36591 |
| *PRO12* | -2.318572 | -0.425879 | 9.177881 | 13.84659 | 13.21294 |
| *PRO13* | -0.727743 | -2.540377 | 1.025192 | 1.709432 | 10.03028 |
| *PRO14* | -3.754084 | -1.312353 | 9.506977 | 14.16365 | 13.6633 |
| *PRO22* | -5.902214 | -4.713594 | 3.181314 | 7.486963 | 9.470546 |
| *PRO23* | -2.74985 | 1.28564 | 10.40716 | 14.40732 | 15.25907 |
| *PRO24* | -0.404587 | 1.84159 | 6.789933 | 9.088482 | 9.325067 |
| *PRO25* | -3.401215 | -2.634295 | -2.909177 | -0.477651 | 0.897611 |
| *GLU8,9* | -2.662161 | 4.235087 | 10.54881 | 13.8347 | 13.27147 |
| *GLU10,11* | -8.133274 | -3.591566 | 5.136018 | 8.144802 | 0.853647 |
| *PRO5,7, 10* | -8.240623 | -5.537846 | 3.973587 | 8.556809 | 13.05103 |
| *PRO19,20,21* | -1.703276 | -0.761973 | 8.537558 | 13.49309 | 13.7103 |

**Table S11. Support Vector Machine (SVM) prot analysis for SVM classification of a protein into a functional family from its primary sequence.**

| Query | R value | P value (%) |
|---|---|---|
| Albumin | | |
| Query=ALB1 | | |
| Length=340 amino acids | | |
| Transmembrane | 4.9 | 98.6 |
| All lipid-binding proteins | 2.3 | 88.1 |
| Iron-binding | 1.4 | 71.3 |
| Metal-binding | 1 | 58.6 |
| Photosystem I | 1 | 58.6 |
| Query=ALB2 | | |
| Length=68 amino acids | | |
| Lipid synthesis | 1 | 58.6 |
| Calcium-binding | 1 | 58.6 |
| Photoreceptor | 1 | 58.6 |
| EC 1.9.-.-: Oxidoreductases - Acting on a heme group of donors | 1 | 58.6 |
| TC 1.C. Channels/Pores - Pore-forming toxins (proteins and peptides) | 1 | 58.6 |
| Outer membrane | 1 | 58.6 |
| Sodium-binding | 1 | 58.6 |
| Query=ALB3 | | |
| Length=134 amino acids | | |
| Metal-binding | 1 | 58.6 |
| Transmembrane | 1 | 58.6 |
| TC 1.C. Channels/Pores - Pore-forming toxins (proteins and peptides) | 1 | 58.6 |
| EC 4.2.-.-: Lyases - Carbon-Oxygen Lyases | 1 | 58.6 |
| Query=ALB4 | | |
| Length=186 amino acids | | |
| All lipid-binding proteins | 2.3 | 88.1 |
| Outer membrane | 1 | 58.6 |
| Metal-binding | 1 | 58.6 |
| DNA repair | 1 | 58.6 |
| Calcium-binding | 1 | 58.6 |
| Magnesium-binding | 1 | 58.6 |
| Query=ALB5 | | |
| Length=154 amino acids | | |
| All lipid-binding proteins | 1.3 | 68.5 |
| EC 2.7.-.-: Transferases - Transferring Phosphorus-Containing Groups | 1.2 | 65.4 |
| Iron-binding | 1.1 | 62.2 |
| Outer membrane | 1 | 58.6 |
| Chlorophyll biosynthesis | 1 | 58.6 |
| TC 3.A.3 P-type ATPase (P-ATPase) family | 1 | 58.6 |
| Magnesium-binding | 1 | 58.6 |
| Query=ALB6 | | |
| Length=167 amino acids | | |
| Zinc-binding | 1.2 | 65.4 |
| Nuclear Receptors | 1.1 | 62.2 |
| Iron-binding | 1 | 58.6 |
| Metal-binding | 1 | 58.6 |
| Calcium-binding | 1 | 58.6 |
| TC 2.A.6 Resistance-nodulation-cell division (RND) family | 1 | 58.6 |
| Lipid degradation | 1 | 58.6 |
| TC 3.A.3 P-type ATPase (P-ATPase) family | 1 | 58.6 |
| Query=ALB7 | | |
| Length=160 amino acids | | |
| Transmembrane | 2.7 | 92.1 |
| All lipid-binding proteins | 1.6 | 76.2 |
| Photosystem I | 1 | 58.6 |
| EC 2.1.-.-: Transferases - Transferring One-Carbon Groups | 1 | 58.6 |
| Query=ALB8 | | |
| Length=163 amino acids | | |
| Transmembrane | 3 | 94.2 |
| All lipid-binding proteins | 2.1 | 85.4 |
| Zinc-binding | 1.9 | 82.2 |
| EC 3.4.-.-: Hydrolases - Acting on peptide bonds (Peptidases) | 1.4 | 71.3 |
| EC 3.1.-.-: Hydrolases - Acting on Ester Bonds | 1.1 | 62.2 |
| TC 1.C. Channels/Pores - Pore-forming toxins (proteins and peptides) | 1 | 58.6 |
| Query=ALB9 | | |
| Length=188 amino acids | | |
| All lipid-binding proteins | 2 | 83.9 |
| Transmembrane | 1.5 | 73.8 |
| Lipoprotein | 1.5 | 73.8 |
| EC 3.4.-.-: Hydrolases - Acting on peptide bonds (Peptidases) | 1.2 | 65.4 |
| Iron-binding | 1.1 | 62.2 |
| Metal-binding | 1 | 58.6 |
| Query=ALB10 | | |
| Length=178 amino acids | | |
| All lipid-binding proteins | 2.5 | 90.3 |
| Lipoprotein | 1.6 | 76.2 |
| Transmembrane | 1.5 | 73.8 |
| EC 3.4.-.-: Hydrolases - Acting on peptide bonds (Peptidases) | 1.1 | 62.2 |
| All DNA-binding | 1 | 58.6 |
| Chlorophyll biosynthesis | 1 | 58.6 |
| Metal-binding | 1 | 58.6 |
| Lipopolysaccharide biosynthesis | 1 | 58.6 |
| EC 4.1.-.-: Lyases - Carbon-Carbon Lyases | 1 | 58.6 |
| Copper-binding | 1 | 58.6 |
| EC 2.3.-.-: Transferases - Acyltransferases | 1 | 58.6 |
| EC 4.2.-.-: Lyases - Carbon-Oxygen Lyases | 1 | 58.6 |
| Query=ALB11 | | |
| Length=160 amino acids | | |
| Transmembrane | 2.4 | 89.3 |
| All lipid-binding proteins | 2.2 | 86.8 |
| Zinc-binding | 1.3 | 68.5 |
| EC 3.4.-.-: Hydrolases - Acting on peptide bonds (Peptidases) | 1.2 | 65.4 |
| DNA repair | 1 | 58.6 |
| EC 3.5.-.-: Hydrolases - Acting on Carbon-Nitrogen Bonds, other than Peptide Bonds | 1 | 58.6 |
| TC 1.C. Channels/Pores - Pore-forming toxins (proteins and peptides) | 1 | 58.6 |
| Query=ALB12 | | |
| Length=158 amino acids | | |
| Transmembrane | 2.5 | 90.3 |
| All lipid-binding proteins | 1.8 | 80.4 |
| Zinc-binding | 1.6 | 76.2 |
| EC 3.4.-.-: Hydrolases - Acting on peptide bonds (Peptidases) | 1.5 | 73.8 |
| EC 3.1.-.-: Hydrolases - Acting on Ester Bonds | 1.3 | 68.5 |
| EC 2.7.-.-: Transferases - Transferring Phosphorus-Containing Groups | 1.3 | 68.5 |
| Photosynthesis | 1.1 | 62.2 |
| Photorespiration | 1.1 | 62.2 |
| Chlorophyll biosynthesis | 1 | 58.6 |
| Metal-binding | 1 | 58.6 |
| EC 4.1.-.-: Lyases - Carbon-Carbon Lyases | 1 | 58.6 |
| Query=ALB13 | | |
| Length=159 amino acids | | |
| Transmembrane | 2.2 | 86.8 |
| EC 3.4.-.-: Hydrolases - Acting on peptide bonds (Peptidases) | 1.6 | 76.2 |
| Photorespiration | 1.1 | 62.2 |
| EC 4.1.-.-: Lyases - Carbon-Carbon Lyases | 1.1 | 62.2 |
| Photosynthesis | 1 | 58.6 |

| | | |
|---|---|---|
| Chlorophyll biosynthesis | 1 | 58.6 |
| Metal-binding | 1 | 58.6 |
| Magnesium-binding | 1 | 58.6 |
| Query=ALB14 | | |
| Length=148 amino acids | | |
| Transmembrane | 1.7 | 78.4 |
| All lipid-binding proteins | 1.6 | 76.2 |
| Zinc-binding | 1.5 | 73.8 |
| Iron-binding | 1.2 | 65.4 |
| EC 2.7.-.-: Transferases - Transferring Phosphorus-Containing Groups | 1.2 | 65.4 |
| Chlorophyll biosynthesis | 1 | 58.6 |
| Metal-binding | 1 | 58.6 |
| EC 1.3.-.-: Oxidoreductases - Acting on the CH-CH group of donors | 1 | 58.6 |
| Outer membrane | 1 | 58.6 |
| DNA repair | 1 | 58.6 |
| Magnesium-binding | 1 | 58.6 |
| Query=ALB15 | | |
| Length=100 amino acids | | |
| All lipid-binding proteins | 1.2 | 65.4 |
| All DNA-binding | 1 | 58.6 |
| Zinc-binding | 1 | 58.6 |
| Outer membrane | 1 | 58.6 |
| EC 3.6.-.-: Hydrolases - Acting on Acid Anhydrides | 1 | 58.6 |
| RNA-binding Proteins | 1 | 58.6 |
| Magnesium-binding | 1 | 58.6 |
| Query=ALB16 | | |
| Length=135 amino acids | | |
| Photosynthesis | 1 | 58.6 |
| Antigen | 1 | 58.6 |
| Nuclear Receptors | 1 | 58.6 |
| EC 1.9.-.-: Oxidoreductases - Acting on a heme group of donors | 1 | 58.6 |
| EC 1.2.-.-: Oxidoreductases - Acting on the aldehyde or oxo group of donors | 1 | 58.6 |
| TC 1.C. Channels/Pores - Pore-forming toxins (proteins and peptides) | 1 | 58.6 |
| Chlorophyll | 1 | 58.6 |
| Sodium-binding | 1 | 58.6 |
| EC 1.10.-.-: Oxidoreductases - Acting on diphenols and related substances as donor | 1 | 58.6 |
| Glutelin | | |
| Query=GLU1 | | |
| Length=236 amino acids | | |
| All lipid-binding proteins | 2 | 83.9 |
| Outer membrane | 1 | 58.6 |
| Metal-binding | 1 | 58.6 |
| DNA repair | 1 | 58.6 |
| TC 3.A.5 Type II (general) secretory pathway (IISP) family | 1 | 58.6 |
| Query=GLU2 | | |
| Length=499 amino acids | | |
| Transmembrane | 7.1 | 99 |
| All lipid-binding proteins | 5.1 | 98.8 |
| TC 8.A. Accessory Factors Involved in Transport - Auxiliary transport proteins | 1.6 | 76.2 |
| Outer membrane | 1 | 58.6 |
| Actin binding | 1 | 58.6 |
| Query=GLU3 | | |
| Length=377 amino acids | | |
| EC 3.1.-.-: Hydrolases - Acting on Ester Bonds | 3.2 | 95.2 |
| All lipid-binding proteins | 1.6 | 76.2 |
| Metal-binding | 1.3 | 68.5 |
| Iron-binding | 1.1 | 62.2 |
| EC 3.4.-.-: Hydrolases - Acting on peptide bonds (Peptidases) | 1.1 | 62.2 |
| Outer membrane | 1 | 58.6 |
| Actin binding | 1 | 58.6 |
| Copper-binding | 1 | 58.6 |
| Magnesium-binding | 1 | 58.6 |
| Query=GLU4 | | |
| Length=499 amino acids | | |
| Transmembrane | 5.6 | 98.9 |
| EC 3.1.-.-: Hydrolases - Acting on Ester Bonds | 2.2 | 86.8 |
| All lipid-binding proteins | 1.8 | 80.4 |
| Iron-binding | 1.5 | 73.8 |
| EC 2.7.-.-: Transferases - Transferring Phosphorus-Containing Groups | 1.1 | 62.2 |
| Calcium-binding | 1 | 58.6 |
| Outer membrane | 1 | 58.6 |
| Actin binding | 1 | 58.6 |
| DNA repair | 1 | 58.6 |
| TC 3.A.5 Type II (general) secretory pathway (IISP) family | 1 | 58.6 |
| Magnesium-binding | 1 | 58.6 |
| Query=GLU5 | | |
| Length=495 amino acids | | |
| Transmembrane | 8.8 | 99.2 |
| All lipid-binding proteins | 2.2 | 86.8 |
| EC 2.7.-.-: Transferases - Transferring Phosphorus-Containing Groups | 1.9 | 82.2 |
| Zinc-binding | 1.7 | 78.4 |
| Coat protein | 1.4 | 71.3 |
| Iron-binding | 1.2 | 65.4 |
| Copper-binding | 1 | 58.6 |
| TC 3.A.5 Type II (general) secretory pathway (IISP) family | 1 | 58.6 |
| Query=GLU6 | | |
| Length=484 amino acids | | |
| All lipid-binding proteins | 3.9 | 97.5 |
| Transmembrane | 1.2 | 65.4 |
| Metal-binding | 1 | 58.6 |
| Outer membrane | 1 | 58.6 |
| Actin binding | 1 | 58.6 |
| DNA repair | 1 | 58.6 |
| Copper-binding | 1 | 58.6 |
| Query=GLU7 | | |
| Length=495 amino acids | | |
| Transmembrane | 10 | 99.2 |
| Zinc-binding | 2.9 | 93.6 |
| All lipid-binding proteins | 2.9 | 93.6 |
| EC 3.1.-.-: Hydrolases - Acting on Ester Bonds | 1.3 | 68.5 |
| Calcium-binding | 1 | 58.6 |
| DNA repair | 1 | 58.6 |
| Photosystem I | 1 | 58.6 |
| Copper-binding | 1 | 58.6 |
| Query=GLU8 | | |
| Length=499 amino acids | | |
| Transmembrane | 9.6 | 99.2 |
| Zinc-binding | 3.2 | 95.2 |
| All lipid-binding proteins | 2.2 | 86.8 |
| EC 3.1.-.-: Hydrolases - Acting on Ester Bonds | 1.8 | 80.4 |
| Calcium-binding | 1 | 58.6 |
| DNA repair | 1 | 58.6 |
| Photosystem I | 1 | 58.6 |
| Query=GLU9 | | |
| Length=499 amino acids | | |
| Transmembrane | 9.6 | 99.2 |
| Zinc-binding | 3.2 | 95.2 |
| All lipid-binding proteins | 2.2 | 86.8 |

| | | |
|---|---|---|
| EC 3.1.-.-: Hydrolases - Acting on Ester Bonds | 1.8 | 80.4 |
| Calcium-binding | 1 | 58.6 |
| DNA repair | 1 | 58.6 |
| Photosystem I | 1 | 58.6 |
| Query=GLU10 | | |
| Length=500 amino acids | | |
| Transmembrane | 4.7 | 98.5 |
| EC 3.1.-.-: Hydrolases - Acting on Ester Bonds | 2.4 | 89.3 |
| Iron-binding | 2.2 | 86.8 |
| TC 8.A. Accessory Factors Involved in Transport - Auxiliary transport proteins | 1.5 | 73.8 |
| Structural protein (Matrix protein,Core protein,Viral occlusion body,Keratin) | 1.3 | 68.5 |
| All DNA-binding | 1.1 | 62.2 |
| Zinc-binding | 1 | 58.6 |
| Calcium-binding | 1 | 58.6 |
| Outer membrane | 1 | 58.6 |
| DNA repair | 1 | 58.6 |
| Photosystem I | 1 | 58.6 |
| TC 3.A.5 Type II (general) secretory pathway (IISP) family | 1 | 58.6 |
| Query=GLU11 | | |
| Length=500 amino acids | | |
| Transmembrane | 4.3 | 98.1 |
| EC 3.1.-.-: Hydrolases - Acting on Ester Bonds | 2.4 | 89.3 |
| Iron-binding | 2.2 | 86.8 |
| Structural protein (Matrix protein,Core protein,Viral occlusion body,Keratin) | 1.5 | 73.8 |
| TC 8.A. Accessory Factors Involved in Transport - Auxiliary transport proteins | 1.4 | 71.3 |
| Zinc-binding | 1.3 | 68.5 |
| All DNA-binding | 1.1 | 62.2 |
| All lipid-binding proteins | 1.1 | 62.2 |
| Calcium-binding | 1 | 58.6 |
| Outer membrane | 1 | 58.6 |
| DNA repair | 1 | 58.6 |
| Photosystem I | 1 | 58.6 |
| Transmembrane | 7.4 | 99 |
| Iron-binding | 4.3 | 98.1 |
| EC 3.1.-.-: Hydrolases - Acting on Ester Bonds | 1.2 | 65.4 |
| Calcium-binding | 1 | 58.6 |
| Outer membrane | 1 | 58.6 |
| DNA repair | 1 | 58.6 |
| Query=GLU13 | | |
| Length=376 amino acids | | |
| All lipid-binding proteins | 4.4 | 98.2 |
| Transmembrane | 2.9 | 93.6 |
| EC 1.11.-.-: Oxidoreductases - Acting on a peroxide as acceptor | 2.3 | 88.1 |
| Zinc-binding | 2.2 | 86.8 |
| EC 3.4.-.-: Hydrolases - Acting on peptide bonds (Peptidases) | 1.3 | 68.5 |
| Actin binding | 1 | 58.6 |
| All lipid-binding proteins | 6.1 | 99 |
| TC 8.A. Accessory Factors Involved in Transport - Auxiliary transport proteins | 1.4 | 71.3 |
| Outer membrane | 1 | 58.6 |
| Actin binding | 1 | 58.6 |
| Query=GLU15 | | |
| Length=359 amino acids | | |
| All lipid-binding proteins | 3.8 | 97.3 |
| EC 3.1.-.-: Hydrolases - Acting on Ester Bonds | 1.9 | 82.2 |
| EC 3.5.-.-: Hydrolases - Acting on Carbon-Nitrogen Bonds, other than Peptide Bonds | 1.9 | 82.2 |
| Outer membrane | 1 | 58.6 |
| Photosystem I | 1 | 58.6 |
| Copper-binding | 1 | 58.6 |
| Query=GLU17 | | |
| Length=325 amino acids | | |
| EC 2.3.-.-: Transferases - Acyltransferases | 2.4 | 89.3 |
| Chlorophyll biosynthesis | 1 | 58.6 |
| Calcium-binding | 1 | 58.6 |
| Magnesium-binding | 1 | 58.6 |
| Query=GLU18 | | |
| Length=350 amino acids | | |
| EC 2.3.-.-: Transferases - Acyltransferases | 2.1 | 85.4 |
| EC 2.4.-.-: Transferases - Glycosyltransferases | 1.3 | 68.5 |
| EC 1.3.-.-: Oxidoreductases - Acting on the CH-CH group of donors | 1.1 | 62.2 |
| EC 1.2.-.-: Oxidoreductases - Acting on the aldehyde or oxo group of donors | 1.1 | 62.2 |
| Calcium-binding | 1 | 58.6 |
| All lipid-binding proteins | 1 | 58.6 |
| DNA repair | 1 | 58.6 |
| Query=GLU19 | | |
| Length=354 amino acids | | |
| EC 2.3.-.-: Transferases - Acyltransferases | 1.8 | 80.4 |
| EC 1.3.-.-: Oxidoreductases - Acting on the CH-CH group of donors | 1.4 | 71.3 |
| Calcium-binding | 1 | 58.6 |
| Magnesium-binding | 1 | 58.6 |
| Query=GLU20 | | |
| Length=499 amino acids | | |
| Transmembrane | 5.8 | 99 |
| All lipid-binding proteins | 4.3 | 98.1 |
| Photosynthesis | 1.1 | 62.2 |
| Metal-binding | 1.1 | 62.2 |
| Outer membrane | 1 | 58.6 |
| Actin binding | 1 | 58.6 |
| EC 2.1.-.-: Transferases - Transferring One-Carbon Groups | 1 | 58.6 |
| Globulin | | |
| Query=GLB1 | | |
| Length=656 amino acids | | |
| All lipid-binding proteins | 1.4 | 71.3 |
| Zinc-binding | 1.3 | 68.5 |
| Calcium-binding | 1 | 58.6 |
| Query=GLB2 | | |
| Length=565 amino acids | | |
| EC 4.2.-.-:Lyases-Carbon-Oxygen Lyases | 1 | 58.6 |
| EC 2.4.-.-: Transferases - Glycosyltransferases | 4.3 | 98.1 |
| EC 1.2.-.-: Oxidoreductases - Acting on the aldehyde or oxo group of donors | 3.1 | 94.7 |
| EC 1.1.-.-: Oxidoreductases - Acting on the CH-OH group of donors | 2.1 | 85.4 |
| All lipid-binding proteins | 2.1 | 85.4 |
| Metal-binding | 2 | 83.9 |
| EC 3.4.-.-: Hydrolases - Acting on peptide bonds (Peptidases) | 1.6 | 76.2 |
| EC 1.7.-.-: Oxidoreductases - Acting on other nitrogenous compounds as donors | 1.5 | 73.8 |
| Zinc-binding | 1.2 | 65.4 |
| Transmembrane | 1.2 | 65.4 |
| EC 4.1.-.-: Lyases - Carbon-Carbon Lyases | 1.1 | 62.2 |
| Magnesium-binding | 1 | 58.6 |
| Query=GLB3 | | |
| Length=562 amino acids | | |
| Metal-binding | 1.1 | 62.2 |
| Query=GLB4 | | |
| Length=470 amino acids | | |
| All lipid-binding proteins | 3.9 | 97.5 |
| Metal-binding | 1.9 | 82.2 |
| Zinc-binding | 1.3 | 68.5 |
| Calcium-binding | 1 | 58.6 |
| DNA repair | 1 | 58.6 |

| | | |
|---|---|---|
| Prolamin | | |
| Query=PRO1 | | |
| Length=76 amino acids | | |
| All DNA-binding | 1 | 58.6 |
| Photosynthesis | 1 | 58.6 |
| snRNA-binding Proteins | 1 | 58.6 |
| Metal-binding | 1 | 58.6 |
| TC 3.A.15 The Outer Membrane Protein Secreting Main Terminal Branch (MTB) fam | 1 | 58.6 |
| Photosystem I | 1 | 58.6 |
| mRNA-binding Proteins | 1 | 58.6 |
| Calcium-binding | 1 | 58.6 |
| Photoreceptor | 1 | 58.6 |
| Query=PRO2 | | |
| Length=118 amino acids | | |
| Outer membrane | | |
| Chlorophyll biosynthesis | 1 | 58.6 |
| Magnesium-binding | 1 | 58.6 |
| Query=PRO3 | | |
| Length=150 amino acids | | |
| Calcium-binding | 1 | 58.6 |
| Photoreceptor | 1 | 58.6 |
| Repressor | 1 | 58.6 |
| DNA repair | 1 | 58.6 |
| Lipid degradation | | |
| Metal-binding | 1 | 58.6 |
| Structural protein (Matrix protein,Core protein,Viral occlusion body,Keratin) | 1 | 58.6 |
| EC 4.1.-.-: Lyases - Carbon-Carbon Lyases | 1 | 58.6 |
| EC 3.8.-.-: Hydrolases - Acting on Halide Bonds | 1 | 58.6 |
| Copper-binding | 1 | 58.6 |
| Calcium-binding | 1 | 58.6 |
| Query=PRO4 | | |
| Length=104 amino acids | | |
| Metal-binding | | |
| Query=PRO5 | | |
| Length=150 amino acids | | |
| Lipid degradation | | |
| Metal-binding | 1 | 58.6 |
| Structural protein (Matrix protein,Core protein,Viral occlusion body,Keratin) | 1 | 58.6 |
| EC 4.1.-.-: Lyases - Carbon-Carbon Lyases | 1 | 58.6 |
| EC 3.8.-.-: Hydrolases - Acting on Halide Bonds | 1 | 58.6 |
| Copper-binding | 1 | 58.6 |
| Calcium-binding | 1 | 58.6 |
| Query=PRO6 | | |
| Length=148 amino acids | | |
| Transmembrane | | |
| EC 2.5.-.-: Transferases - Transferring Alkyl or Aryl Groups, Other than Methyl Grou | 1.2 | 65.4 |
| All lipid-binding proteins | 1.1 | 62.2 |
| EC 3.4.-.-: Hydrolases - Acting on peptide bonds (Peptidases) | 1.1 | 62.2 |
| Calcium-binding | 1 | 58.6 |
| Outer membrane | 1 | 58.6 |
| Copper-binding | 1 | 58.6 |
| Magnesium-binding | 1 | 58.6 |
| Query=PRO7 | | |
| Length=150 amino acids | | |
| Metal-binding | | |
| EC 3.8.-.-: Hydrolases - Acting on Halide Bonds | 1 | 58.6 |
| Calcium-binding | 1 | 58.6 |
| Lipid degradation | 1 | 58.6 |
| Structural protein (Matrix protein,Core protein,Viral occlusion body,Keratin) | 1 | 58.6 |
| EC 4.1.-.-: Lyases - Carbon-Carbon Lyases | 1 | 58.6 |
| Query=PRO8 | | |
| Length=150 amino acids | | |
| Metal-binding | | |
| EC 3.8.-.-: Hydrolases - Acting on Halide Bonds | 1 | 58.6 |
| Calcium-binding | 1 | 58.6 |
| Lipid degradation | 1 | 58.6 |
| Structural protein (Matrix protein,Core protein,Viral occlusion body,Keratin) | 1 | 58.6 |
| EC 4.1.-.-: Lyases - Carbon-Carbon Lyases | 1 | 58.6 |
| Copper-binding | 1 | 58.6 |
| Query=PRO9 | | |
| Length=148 amino acids | | |
| Transmembrane | | |
| EC 2.5.-.-: Transferases - Transferring Alkyl or Aryl Groups, Other than Methyl Grou | 2 | 83.9 |
| Calcium-binding | 1.4 | 71.3 |
| Hormone | 1 | 58.6 |
| Copper-binding | 1 | 58.6 |
| EC 5.3.-.-: Isomerases - Intramolecular Oxidoreductases | 1 | 58.6 |
| Query=PRO10 | | |
| Length=150 amino acids | | |
| Metal-binding | | |
| EC 3.8.-.-: Hydrolases - Acting on Halide Bonds | 1 | 58.6 |
| Calcium-binding | 1 | 58.6 |
| Lipid degradation | 1 | 58.6 |
| Structural protein (Matrix protein,Core protein,Viral occlusion body,Keratin) | 1 | 58.6 |
| EC 4.1.-.-: Lyases - Carbon-Carbon Lyases | 1 | 58.6 |
| Copper-binding | 1 | 58.6 |
| Query=PRO11 | | |
| Length=153 amino acids | | |
| All lipid-binding proteins | 1 | 58.6 |
| Query=PRO12 | | |
| Length=153 amino acids | | |
| Lipid degradation | 1 | 58.6 |
| Copper-binding | 1 | 58.6 |
| Query=PRO13 | | |
| Length=153 amino acids | | |
| Metal-binding | 1 | 58.6 |
| TC 3.A.5 Type II (general) secretory pathway (IISP) family | 1 | 58.6 |
| Query=Pro14 | | |
| Length=153 amino acids | | |
| Based on Support Vector Machine classification, | | |
| your protein may belong to the following families: | | |
| All lipid-binding proteins | 1 | 58.6 |
| Query=PRO15 | | |
| Length=115 amino acids | | |
| Nickel-binding | | |
| Cobalt-binding | 1 | 58.6 |
| All DNA-binding | 1 | 58.6 |
| Zinc-binding | 1 | 58.6 |
| Metal-binding | 1 | 58.6 |
| Cell adhesion | 1 | 58.6 |
| Magnesium-binding | 1 | 58.6 |
| Query=PRO16 | | |
| Length=149 amino acids | | |
| All lipid-binding proteins | | |
| Immune response | 1.1 | 62.2 |
| Photoreceptor | 1 | 58.6 |
| TC 3.A.1 ATP-binding cassette (ABC) family | 1 | 58.6 |
| Actin binding | 1 | 58.6 |

| | | |
|---|---|---|
| DNA repair | 1 | 58.6 |
| Query=PRO17 | | |
| Length=156 amino acids | | |
| Nuclear Receptors | | |
| TC 3.A.1 ATP-binding cassette (ABC) family | 1 | 58.6 |
| TC 1.C. Channels/Pores - Pore-forming toxins (proteins and peptides) | 1 | 58.6 |
| Outer membrane | 1 | 58.6 |
| Query=PRO18 | | |
| Length=156 amino acids | | |
| Metal-binding | | |
| EC 3.4.-.-: Hydrolases - Acting on peptide bonds (Peptidases) | 1 | 58.6 |
| Nuclear Receptors | 1 | 58.6 |
| TC 3.A.1 ATP-binding cassette (ABC) family | 1 | 58.6 |
| TC 1.C. Channels/Pores - Pore-forming toxins (proteins and peptides) | 1 | 58.6 |
| Outer membrane | 1 | 58.6 |
| Structural protein (Matrix protein,Core protein,Viral occlusion body,Keratin) | 1 | 58.6 |
| Query=PRO19 | | |
| Length=151 amino acids | | |
| Iron-binding | | |
| Metal-binding | 1 | 58.6 |
| EC 3.4.-.-: Hydrolases - Acting on peptide bonds (Peptidases) | 1 | 58.6 |
| Calcium-binding | 1 | 58.6 |
| TC 3.A.1 ATP-binding cassette (ABC) family | 1 | 58.6 |
| Potassium-binding | 1 | 58.6 |
| Lipid degradation | 1 | 58.6 |
| All lipid-binding proteins | 1 | 58.6 |
| Magnesium-binding | 1 | 58.6 |
| Query=PRO20 | | |
| Length=151 amino acids | | |
| Iron-binding | | |
| Metal-binding | 1 | 58.6 |
| EC 3.4.-.-: Hydrolases - Acting on peptide bonds (Peptidases) | 1 | 58.6 |
| Lipid degradation | 1 | 58.6 |
| All lipid-binding proteins | 1 | 58.6 |
| Query=PRO21 | | |
| Length=151 amino acids | | |
| Metal-binding | | |
| Calcium-binding | 1 | 58.6 |
| Lipid degradation | 1 | 58.6 |
| Query=PRO22 | | |
| Length=156 amino acids | | |
| Transmembrane | | |
| Immune response | 1.1 | 62.2 |
| Metal-binding | 1 | 58.6 |
| EC 3.4.-.-: Hydrolases - Acting on peptide bonds (Peptidases) | 1 | 58.6 |
| Nuclear Receptors | 1 | 58.6 |
| TC 3.A.1 ATP-binding cassette (ABC) family | 1 | 58.6 |
| Lipid degradation | 1 | 58.6 |
| Outer membrane | 1 | 58.6 |
| DNA repair | 1 | 58.6 |
| Copper-binding | 1 | 58.6 |
| Magnesium-binding | 1 | 58.6 |
| Query=PRO23 | | |
| Length=156 amino acids | | |
| Antigen | | |
| TC 3.A.1 ATP-binding cassette (ABC) family | 1 | 58.6 |
| TC 1.C. Channels/Pores - Pore-forming toxins (proteins and peptides) | 1 | 58.6 |
| Outer membrane | 1 | 58.6 |
| All lipid-binding proteins | 1 | 58.6 |
| Query=PRO24 | | |
| Length=155 amino acids | | |
| Photosynthesis | | |
| Zinc-binding | 1 | 58.6 |
| EC 3.1.-.-: Hydrolases - Acting on Ester Bonds | 1 | 58.6 |
| Photoreceptor | 1 | 58.6 |
| EC 1.2.-.-: Oxidoreductases - Acting on the aldehyde or oxo group of donors | 1 | 58.6 |
| EC 3.6.-.-: Hydrolases - Acting on Acid Anhydrides | 1 | 58.6 |
| All lipid-binding proteins | 1 | 58.6 |
| Repressor | 1 | 58.6 |
| Structural protein (Matrix protein,Core protein,Viral occlusion body,Keratin) | 1 | 58.6 |
| Photosystem I | 1 | 58.6 |

**Table S12. Distribution of different domains in rice seed storage proteins.**

| | Domain Name | Start | End | Domain Name | Start | End | Domain Name | Start | End | Domain Name | Start | End |
|---|---|---|---|---|---|---|---|---|---|---|---|---|
| **Albumin** | | | | | | | | | | | | |
| **ALB1** | Bifunctional inhibitor/plant lipid transfer protein/seed storage helical domain | 10 | 127 | | | | | | | | | |
| **ALB2** | | 8 | 65 | | | | | | | | | |
| **ALB3** | | 9 | 134 | | | | | | | | | |
| **ALB4** | | 34 | 183 | | | | | | | | | |
| **ALB5** | | 18 | 140 | | | | | | | | | |
| **ALB6** | | 59 | 163 | | | | | | | | | |
| **ALB7** | | 37 | 153 | | | | | | | | | |
| **ALB8** | | 36 | 156 | | | | | | | | | |
| **ALB9** | | 60 | 180 | | | | | | | | | |
| **ALB10** | | 59 | 177 | | | | | | | | | |
| **ALB11** | | 41 | 157 | | | | | | | | | |
| **ALB12** | | 31 | 152 | | | | | | | | | |
| **ALB13** | | 34 | 147 | | | | | | | | | |
| **ALB14** | | 30 | 138 | | | | | | | | | |
| **ALB15** | | 3 | 56 | | | | | | | | | |
| **ALB16** | | 10 | 127 | | | | | | | | | |
| **Globulin** | | | | | | | | | | | | |
| **GLB1** | Rml-C1 | 63 | 267 | Rml-C2 | 315 | 527 | Cupin1 | 107 | 212 | Cupin2 | 322 | 483 |
| **GLB2** | | 70 | 272 | | 350 | 540 | | 124 | 220 | | 363 | 507 |
| **GLB3** | | 102 | 293 | | 280 | 521 | | 114 | 238 | | 312 | 507 |
| **GLB4** | | 53 | 245 | | 232 | 460 | | 76 | 173 | | 261 | 428 |
| **Glutelin** | | | | | | | | | | | | |
| **GLU1** | Cupin1 | 107 | 207 | Cupin2 | - | - | Rmlc | 4 | 213 | 11s seed storage | - | - |
| **GLU2** | | 51 | 211 | | 320 | 468 | | 45 | 475 | | 306 | 328 |
| **GLU3** | | 21 | 169 | | 220 | 350 | | 190 | 371 | | - | - |
| **GLU4** | | 5 | 207 | | 312 | 460 | | 37 | 471 | | - | - |
| **GLU5** | | 50 | 206 | | 311 | 459 | | 44 | 467 | | 298 | 320 |
| **GLU6** | | 43 | 199 | | 299 | 446 | | 30 | 457 | | 285 | 307 |
| **GLU7** | | 50 | 206 | | 312 | 459 | | 44 | 470 | | 298 | 320 |
| **GLU8** | | 50 | 206 | | 316 | 463 | | 44 | 474 | | 302 | 324 |
| **GLU9** | | 50 | 206 | | 316 | 463 | | 44 | 474 | | 302 | 324 |
| **GLU10** | | 50 | 206 | | 316 | 464 | | 44 | 475 | | 303 | 325 |
| **GLU11** | | 50 | 206 | | 316 | 464 | | 44 | 475 | | 303 | 325 |
| **GLU12** | | 53 | 216 | | 333 | 478 | | 40 | 490 | | 330 | 347 |
| **GLU13** | | 56 | 206 | | 216 | 347 | | 53 | 248 | | - | - |
| **GLU14** | | 50 | 210 | | 319 | 466 | | 44 | 477 | | 305 | 327 |
| **GLU15** | | 7 | 157 | | 206 | 338 | | 176 | 357 | | - | - |
| | | | | | | | | 6 | 184 | | | |
| **GLU16** | | 47 | 234 | | 353 | 491 | | 46 | 502 | | - | - |
| **GLU17** | | 12 | 103 | | 220 | 316 | | 14 | 163 | | - | - |
| | | | | | | | | 134 | 325 | | | |
| **GLU18** | | 10 | 119 | | 192 | 344 | | 11 | 182 | | - | - |
| | | | | | | | | 208 | 347 | | | |
| **GLU19** | | 13 | 121 | | 195 | 348 | | 6 | 354 | | - | - |
| **GLU20** | | 51 | 211 | | 320 | 468 | | 45 | 475 | | 306 | 328 |
| **Prolamins** | | | | | | | | | | | | |
| **PROL1** | Bifunctional inhibitor/plant lipid transfer protein/seed storage helical domain/ Gliadin | 7 | 74 | Bifunctional inhibitor/plant lipid transfer protein/seed storage helical domain | - | - | | | | | | |
| **PROL2** | | 54 | 100 | | - | - | | | | | | |
| **PROL3** | | 33 | 99 | | 34 | 122 | | | | | | |
| **PROL4** | | 33 | 99 | | - | - | | | | | | |
| **PROL5** | | 33 | 99 | | 34 | 140 | | | | | | |
| **PROL6** | | 33 | 99 | | 34 | 125 | | | | | | |
| **PROL7** | | 33 | 99 | | 34 | 122 | | | | | | |
| **PROL8** | | 33 | 99 | | 34 | 140 | | | | | | |
| **PROL9** | | 34 | 99 | | 33 | 107 | | | | | | |
| **PROL10** | | 33 | 99 | | 33 | 124 | | | | | | |
| **PROL11** | | 33 | 99 | | 37 | 144 | | | | | | |
| **PROL12** | | 33 | 99 | | 37 | 144 | | | | | | |
| **PROL13** | | 32 | 97 | | 34 | 136 | | | | | | |
| **PROL14** | | 33 | 99 | | 37 | 144 | | | | | | |
| **PROL15** | | - | - | | 5 | 105 | | | | | | |
| **PROL16** | | - | - | | 5 | 139 | | | | | | |
| **PROL17** | | - | - | | 5 | 146 | | | | | | |
| **PROL18** | | - | - | | 5 | 146 | | | | | | |
| **PROL19** | | 32 | 98 | | 34 | 133 | | | | | | |
| **PROL20** | | 32 | 98 | | 34 | 133 | | | | | | |
| **PROL21** | | 32 | 99 | | 34 | 134 | | | | | | |
| **PROL22** | | - | - | | 5 | 146 | | | | | | |
| **PROL23** | | - | - | | 32 | 156 | | | | | | |
| **PROL24** | | - | - | | 31 | 155 | | | | | | |

**Table S13. List of rice SSPs with confidence score, retrieved from AlphaFold Structure Database.**

| Protein names | Uniprot Id | Sequence length | Confidence score |
|---|---|---|---|
| ALB1 | Q10EM0 | 229 | 86.94 |
| ALB2 | A0A0P0W8Q9 | 143 | 62.75 |
| ALB3 | Q7Y0D9 | 136 | 56.78 |
| ALB4 | P29835 | 188 | 61.81 |
| ALB5 | Q8H4M5 | 156 | 67.66 |
| ALB6 | A0A0P0X3X4 | 149 | 49.9 |
| ALB7 | Q0D7S6 | 162 | 76.62 |
| ALB11 | A3BHT7 | 162 | 77.06 |
| ALB12 | Q7X7E6 | 160 | 79.93 |
| ALB13 | Q7X8H9 | 161 | 80.02 |
| ALB14 | Q8GVI3 | 150 | 82.02 |
| ALB15 | Q10ET9 | 529 | 92.15 |
| ALB16 | Q2R363 | 137 | 51.66 |
| GLB1 | A0A0P0VUB4 | 626 | 74.53 |
| GLB2 | A0A0P0VY13 | 339 | 75.19 |
| GLB3 | Q75GX9 | 564 | 81.19 |
| GLB4 | Q852L2 | 472 | 81.33 |
| GLU1 | A0A0P0V8F8 | 133 | 82.88 |
| GLU2 | A2ZY31 | 501 | 81 |
| GLU3 | Q94CS6 | 379 | 93.34 |
| GLU4 | Q6ESW6 | 501 | 82.45 |
| GLU5 | Q0E2E0 | 497 | 81 |
| GLU7 | A0A5S6RCW4 | 497 | 80.81 |
| GLU9 | Q0E2D3 | 501 | 80.69 |
| GLU11 | A3A5D5 | 502 | 80.5 |
| GLU12 | A1YQH2 | 255 | 71.58 |
| GLU13 | A0A0P0VIM4 | 340 | 76.87 |
| GLU14 | Q0DR11 | 498 | 81.44 |
| GLU15 | Q65XA1 | 361 | 95.34 |
| GLU16 | A0A0N7KP80 | 533 | 79.56 |
| GLU18 | A0A0P0XQ36 | 352 | 92.56 |
| GLU19 | A0A0P0XQT5 | 356 | 92.38 |
| PRO1 | A0A0P0WKT5 | 83 | 66.47 |
| PRO2 | Q5W6A3 | 152 | 50.82 |
| PRO3 | E5D3L6 | 106 | 60.53 |
| PRO4 | A0A0N7KKJ8 | 155 | 51.31 |
| PRO6 | Q0DJ44 | 152 | 50.62 |
| PRO7 | A0A0P0WKY1 | 112 | 63.62 |
| PRO8 | A0A0P0WKT6 | 152 | 50.37 |
| PRO9 | Q5W755 | 152 | 50.82 |
| PRO10 | A0A0P0WKV9 | 123 | 49.58 |
| PRO11 | A1YQF0 | 155 | 49.67 |
| PRO12 | A0A0P0WKV0 | 155 | 44.97 |
| PRO13 | Q5W748 | 155 | 43.91 |
| PRO14 | C7J3Z0 | 160 | 61.68 |
| PRO15 | C7J3Z1 | 136 | 55.37 |
| PRO16 | Q0DBY5 | 151 | 58.03 |
| PRO17 | A1YQF4 | 158 | 48.28 |
| PRO18 | Q0D7V7 | 158 | 50.38 |
| PRO19 | Q8GVK9 | 153 | 49.03 |
| PRO21 | Q8GVK7 | 153 | 49.49 |
| PRO22 | Q2QUA9 | 158 | 56.51 |
| PRO23 | P20698 | 158 | 56.97 |
| PRO24 | A0A0P0Y8T9 | 80 | 47.78 |
| PRO25 | C7J9I2 | 80 | 52.87 |

**Table S14. List of five generated models of each protein with pLDDT values**

**ALB8**: reranking models by 'plddt' metric
rank_001_alphafold2_ptm_model_4_seed_000 pLDDT=79.5 pTM=0.697
rank_002_alphafold2_ptm_model_1_seed_000 pLDDT=79.1 pTM=0.687
rank_003_alphafold2_ptm_model_3_seed_000 pLDDT=78.2 pTM=0.678
rank_004_alphafold2_ptm_model_5_seed_000 pLDDT=76 pTM=0.664
rank_005_alphafold2_ptm_model_2_seed_000 pLDDT=74.4 pTM=0.662

**ALB9**: reranking models by 'plddt' metric
rank_001_alphafold2_ptm_model_3_seed_000 pLDDT=71 pTM=0.598
rank_002_alphafold2_ptm_model_2_seed_000 pLDDT=70.5 pTM=0.602
rank_003_alphafold2_ptm_model_4_seed_000 pLDDT=68.5 pTM=0.593
rank_004_alphafold2_ptm_model_1_seed_000 pLDDT=68 pTM=0.591
rank_005_alphafold2_ptm_model_5_seed_000 pLDDT=67.3 pTM=0.56

**ALB10**: reranking models by 'plddt' metric
rank_001_alphafold2_ptm_model_3_seed_000 pLDDT=75 pTM=0.605
rank_002_alphafold2_ptm_model_1_seed_000 pLDDT=74.5 pTM=0.609
rank_003_alphafold2_ptm_model_2_seed_000 pLDDT=73.7 pTM=0.615
rank_004_alphafold2_ptm_model_4_seed_000 pLDDT=73.6 pTM=0.61
rank_005_alphafold2_ptm_model_5_seed_000 pLDDT=71.1 pTM=0.571

**GLU6**: reranking models by 'plddt' metric
rank_001_alphafold2_ptm_model_3_seed_000 pLDDT=81.9 pTM=0.807
rank_002_alphafold2_ptm_model_4_seed_000 pLDDT=81.6 pTM=0.808
rank_003_alphafold2_ptm_model_5_seed_000 pLDDT=81.5 pTM=0.81
rank_004_alphafold2_ptm_model_2_seed_000 pLDDT=81.1 pTM=0.812
rank_005_alphafold2_ptm_model_1_seed_000 pLDDT=80.2 pTM=0.801

**GLU8**: reranking models by 'plddt' metric
rank_001_alphafold2_ptm_model_5_seed_000 pLDDT=81.1 pTM=0.808
rank_002_alphafold2_ptm_model_4_seed_000 pLDDT=80.8 pTM=0.805
rank_003_alphafold2_ptm_model_3_seed_000 pLDDT=80.7 pTM=0.798
rank_004_alphafold2_ptm_model_2_seed_000 pLDDT=79.6 pTM=0.801
rank_005_alphafold2_ptm_model_1_seed_000 pLDDT=79.4 pTM=0.796

**GLU10**: reranking models by 'plddt' metric
rank_001_alphafold2_ptm_model_5_seed_000 pLDDT=80.9 pTM=0.807
rank_002_alphafold2_ptm_model_3_seed_000 pLDDT=80.9 pTM=0.795
rank_003_alphafold2_ptm_model_4_seed_000 pLDDT=80.6 pTM=0.798
rank_004_alphafold2_ptm_model_2_seed_000 pLDDT=79.7 pTM=0.802
rank_005_alphafold2_ptm_model_1_seed_000 pLDDT=79.4 pTM=0.794

**GLU17**: reranking models by 'plddt' metric
rank_001_alphafold2_ptm_model_4_seed_000 pLDDT=88.2 pTM=0.859
rank_002_alphafold2_ptm_model_3_seed_000 pLDDT=88.2 pTM=0.852
rank_003_alphafold2_ptm_model_5_seed_000 pLDDT=87.7 pTM=0.852
rank_004_alphafold2_ptm_model_2_seed_000 pLDDT=86.7 pTM=0.851
rank_005_alphafold2_ptm_model_1_seed_000 pLDDT=86.6 pTM=0.845

**GLU20**: reranking models by 'plddt' metric
rank_001_alphafold2_ptm_model_3_seed_000 pLDDT=81.7 pTM=0.812
rank_002_alphafold2_ptm_model_5_seed_000 pLDDT=81.1 pTM=0.811
rank_003_alphafold2_ptm_model_4_seed_000 pLDDT=81 pTM=0.805
rank_004_alphafold2_ptm_model_2_seed_000 pLDDT=80.3 pTM=0.807
rank_005_alphafold2_ptm_model_1_seed_000 pLDDT=79.6 pTM=0.798

**PRO5**: reranking models by 'plddt' metric
rank_001_alphafold2_ptm_model_3_seed_000 pLDDT=45.1 pTM=0.137
rank_002_alphafold2_ptm_model_4_seed_000 pLDDT=44.7 pTM=0.137
rank_003_alphafold2_ptm_model_2_seed_000 pLDDT=44.1 pTM=0.149
rank_004_alphafold2_ptm_model_1_seed_000 pLDDT=43.8 pTM=0.17
rank_005_alphafold2_ptm_model_5_seed_000 pLDDT=40.4 pTM=0.125

**PRO20**: reranking models by 'plddt' metric
rank_001_alphafold2_ptm_model_3_seed_000 pLDDT=52.1 pTM=0.17
rank_002_alphafold2_ptm_model_1_seed_000 pLDDT=50.5 pTM=0.181
rank_003_alphafold2_ptm_model_4_seed_000 pLDDT=50.4 pTM=0.155
rank_004_alphafold2_ptm_model_5_seed_000 pLDDT=48.1 pTM=0.142
rank_005_alphafold2_ptm_model_2_seed_000 pLDDT=47.3 pTM=0.16
